# Cytochrome ‘Nanowires’ are Physically Limited to Sub-Picoamp Currents that Suffice for Cellular Respiration

**DOI:** 10.1101/2024.12.21.629920

**Authors:** Matthew J. Guberman-Pfeffer, Caleb L. Herron

**Author notes:** **Correspondence:** Matthew J. Guberman-Pfeffer.

## Abstract

Mineral-respiring microorganisms from hydrothermal vents to terrestrial soils express filaments that electrically connect intracellular respiration to extracellular geochemistry. Filaments dubbed “cytochrome nanowires” (CNs) have been resolved by CryoEM, but whether they are the two-decades-long sought-after physiological ‘nanowires’ remains unproven. To assess their functional competence, we analyzed biological redox conduction in all CNs by computing driving forces in the presence of redox anti-cooperativities, reorganization energies with electronic polarizability, and Marcus rates for diffusive and protein-limited flux models. The chain of heme cofactors in any CN must be densely packed to realize weak (≤0.01 eV) electronic coupling for electron transfer, as evidenced by a single Soret band produced from coincidental absorptions on multiple hemes. Dense packing, in turn, has three consequences: (1) limited driving forces (≤|0.3| eV) due to shared electrostatic microenvironments, (2) strong (≤0.12 eV) redox anti-cooperativities that would accentuate the free energy landscape if the linear heme arrangement did not dictate a contra-thermodynamic oxidation order, and (3) an entropic penalty that is offset by thioether ‘tethers’ of the hemes to the protein backbone. These linkages physically necessitate the rate-throttling T-stacked motif (10-fold slower than the other highly conserved slip-stacked motif). If the sequence of slip- and T-stacked hemes in the CNs had the fastest known nanosecond rates at every step, a micron-long filament would carry a diffusive 0.02 pA current at a physiological 0.1 V, or a protein-limited current of 0.2 pA. Actual CNs have sub-optimal (≤10^2^-fold lower), but sufficient conductivities for cellular respiration, with at most thousands of filaments needed for total cellular metabolic flux. Since cells likely discharge less than the 1.0 pA assumed here, and there are multiple pathways besides CNs for expelling electrons, the micro-to-milli-Siemens/cm conductivities are more than sufficient. Reported conductivities once used to argue for metallic-like pili against the cytochrome hypothesis and now illogically attributed to CNs remain inconsistent by 10^2^-10^5^-fold with the physical constraints imposed on biological redox conduction through multiheme architectures.

## 1. Introduction

Prokaryotes from hydrothermal vents to terrestrial soils exhale ∼10^6^ electrons/s/cell (∼100 fA/cell)^1-7^ through filaments^8, 9^ that electrify microbial communities^10^ and biotic-abiotic interfaces.^11^ This ‘rock breathing’ strategy for anaerobic life, known as extracellular electron transfer (EET) is ancient,^12^ ubiquitous,^13, 14^ environmentally significant,^15-18^ and holds promise for sustainable technologies,^19-24^ but only if its mechanistic underpinnings are elucidated.

Filaments from mineral-respiring microorganisms have recently been resolved by cryogenic electron microscopy (CryoEM) to be polymerized multiheme cytochromes.^8, 25-28^ Though the filaments are dubbed “cytochrome nanowires” (CNs), their physiological role as ‘nanowires’ has not been demonstrated. No experiment can currently measure reliably how well a filament of known identity conducts electrons between molecular redox partners under fully hydrated, physiological conditions.^8^ But from a theoretical vantage point, can the theory of biological electron transfer (*i*.*e*., redox conduction)^29^ connect the atomic structures of CNs to their proposed physiological role as mesoscopic electrical conductors?

All known CNs^8, 25-28^ have the basic anatomy summarized in Figure 1. A single type of cofactor—a bis-histidine ligated *c*-type heme—is repeated hundreds of times and stacked in highly conserved geometries^8^ to form a micron-long (linear or branched) spiraling chain, which is encased by a mostly unstructured (≥50% turns and loops) protein sheath. Each filament is constructed of a different Lego-like multi-heme cytochrome that polymerizes either exclusively through non-covalent interactions, or one or more coordination bonds between protomers.

**Figure 1.**
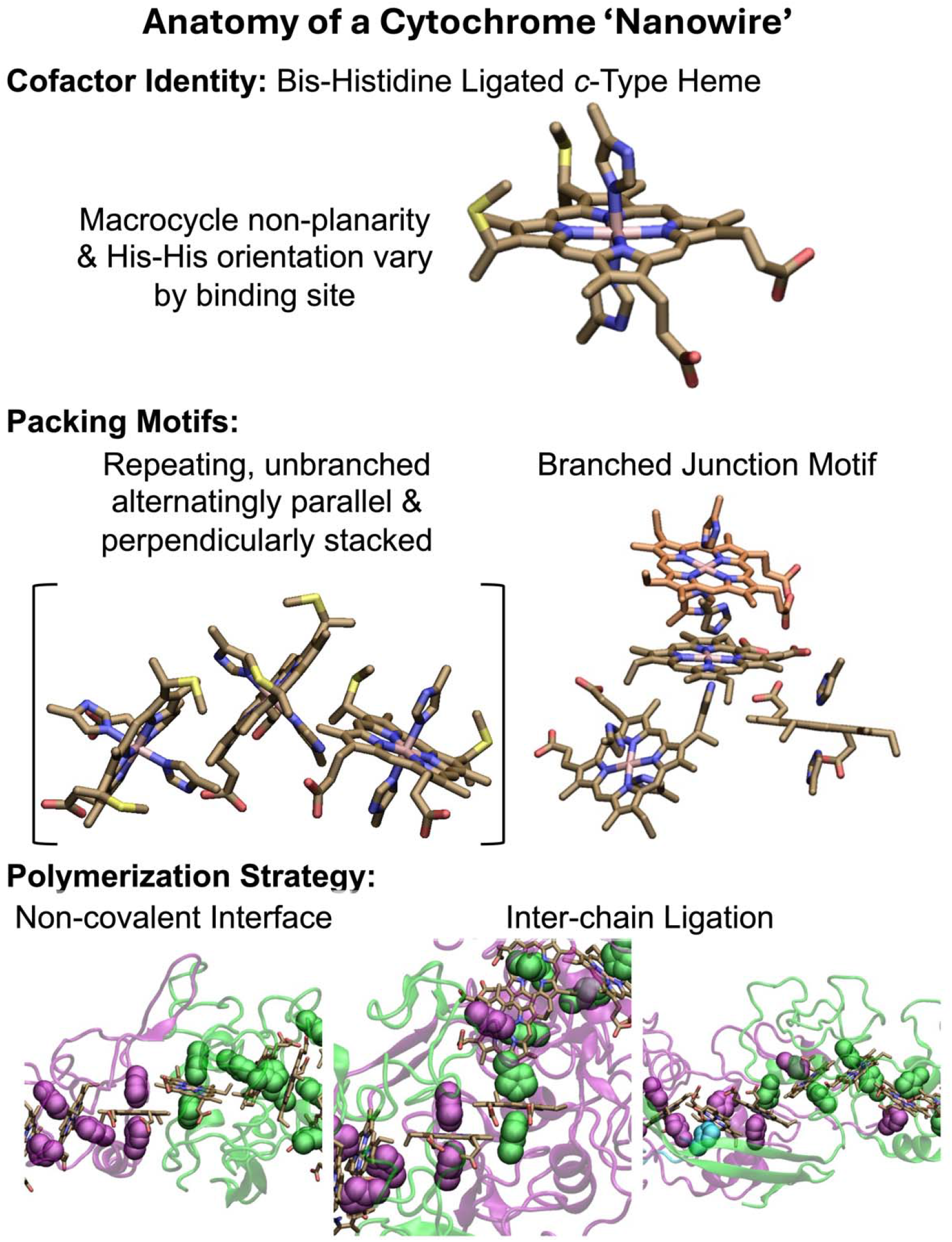
Anatomy of a cytochrome ‘nanowire.’ The heme cofactor, its packing motifs, and filament polymerization strategies are shown. Purple, green, and cyan indicate different protein chains. His ligands are shown in vdW representation. A heme that is coordinated by two differently colored His ligands receives one of them from the adjacent protein chain. The subunit interfaces shown at the bottom from left to right come from OmcZ (PDB 7LQ5),^28^ OmcS (PDB 6EF8),^26^ and F2KMU8 (PDB 8E5G).^8^ The figure was designed with VMD version 1.9.4a57.^137^

The filaments are homopolymers of a tetra-, hexa-, or octaheme protein. The filamentous outer-membrane cytochrome (Omc) types E, S, and Z from *Geobacter sulfurreducens* are respectively composed of these different types of multi-heme building blocks. Two other tetraheme-based filaments come from *Pyrobaculum calidifontis* and *Archaeoglobus veneficus*. Because the latter proteins have not yet been given convenient common names, they will be referred to by the UniProtKB accession codes A3MW92 and F2KMU8, respectively.

The stacking of hemes to form a chain, and the association of proteins to extend that chain is an architectural design found elsewhere in Nature: Chains of 4,^30, 31^ 5, ^32^ 12,^33^ 16,^34^ 20,^35^ 30,^36^ and 192^37^ hemes are formed through non-covalent association,^35, 36, 38, 39^ coordination chemistry,^31^ or genetic grafting and pruning,^33, 40^ of other multi-heme modules. CNs, however, are unique in that multi-heme proteins with two distinct binding interfaces are iteratively paired to form, in principle, an infinitely extendable, cylindrical assembly. That these assemblies extend for μm-lengths beyond the safety of the outer-membrane and contain ∼10^3^ cofactors, each with a metal ion (iron) of limited bioavailability, is remarkable from a biosynthetic cost-benefit perspective.

But the benefits to a microorganism from expressing CNs remain unclear. Why, for example, is a single bacterium, *Geobacter sulfurreducens*, capable of producing three of the five known CNs? Is it because the heme chain common to all CNs have different conductivities that match different respiratory rates, or does the non-homologous protein ‘packaging’ provide environmentally-customized interfaces?^41^

The CryoEM structures have been assumed to explain the molecular underpinnings of EET,^8^ but biochemical data has also been interpreted as implicating an altogether different filament of hypothetical structure.^42^ Solid-state electrical measurements have been assumed to be biologically relevant,^26^ even though the theory of biological electron transfer cannot explain them^43, 44^—an observation that once was used to argue that the filaments are not cytochromes.^45, 46^

Under pressure to validate the solid-state experiments, theoretical models have proposed biologically implausible electronic coherences over 20 nm,^44^ predicted spuriously negative redox potentials and a 0.4 V hysteresis^43, 47^ that was even known beforehand not to exist,^48, 49^ or have modeled biologically irrelevant electron transport (as opposed to transfer) through a molecular junction.^50^

None of these studies have answered the question: Are the CNs resolved by CryoEM *competent* to discharge the metabolic flux of electrons from a microbial cell? Guberman-Pfeffer recently proposed that a *minimum* of seven filaments would suffice for cellular respiration given the conductivity of a generic heme chain.^51^ How does this prediction hold up to computations on the actual structures?

Herein, we characterize the energetics and kinetics in all structurally resolved CNs, compare their conductivities to the needs of cellular respiration, and deduce physical constraints on the conductivity of any biological heme chain. In so doing, we demonstrate that the multiheme architecture physically limits CNs to sub-optimal but sufficient micro-to-millisiemens-per-centimeter conductivities for cellular respiration by EET. If the CNs were the only route for expelling electrons, which is *not* the case, tens of the tetrahemes, hundreds of the hexaheme, or thousands of the octaheme filaments would be needed.

## 2. Methods

All quantum chemical calculations were performed with the Gaussian 16 Rev. A.03 program.^52^ All other calculations were performed with Python packages available on GitHub that were written by the authors. These packages include: (1) BioDC V2.2 (https://github.com/Mag14011/BioDC/releases/tag/V2.2): A program to assess redox potentials, cooperativities, and conductivity in multiheme proteins; (2) ETAnalysis V1.0 (https://github.com/Mag14011/ETAnalysis/releases/tag/V1.0): A program to compute standard Marcus theory rates, charge diffusion constants, and protein-limited, steady-state multi-particle electron fluxes; (3) EFieldAnalysis V1.0 (https://github.com/Mag14011/ElectricFieldAnalysis/releases/tag/V1.0): A program to compute the electric field at multiple probe atoms in a protein over the course of molecular dynamics (MD) trajectories; and (4) PolReorg V1.0 (https://github.com/Mag14011/PolReorg/releases/tag/V1.0): A program to compute the effect of active site polarizability on outer-sphere reorganization energy for electron transfer.

### 2.1 Structure Preparation

Presently available entries in the Protein Data Bank (PDB) for CNs were prepared with BioDC. These include PDB ID 7TFS (OmcE), 8E5F (A3MW92), 8E5G (F2KMU8), 6EF8 and 6NEF (OmcS), and 7LQ5 and 8D9M (OmcZ). BioDC automatically detects the type (*b*/*c*) and axial ligation (His-His/His-Met) of each heme and assigns AMBER FF10 forcefield parameters for the chosen (oxidized/reduced) charge state. Standard protonation states were assigned, but BioDC can prepare each structure for constant pH molecular dynamics simulations (including for the propionic acid heme substituents) if desired. The program has an interactive interface that writes the selected settings to a file that can be used as an input file for reproducibility. Those files for the present analysis are provided in the GitHub repository.

### 2.2 Redox Potentials, Driving Forces, and Redox Cooperativities

BioDC was used to estimate the oxidation energy of heme *i* while all other hemes were in the reduced state (Eq. 1), and the change in the oxidation energy of heme *i* due to the oxidation of heme *j*, while all other hemes were in the reduced state (Eq 2).

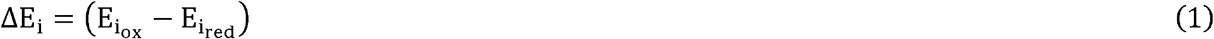

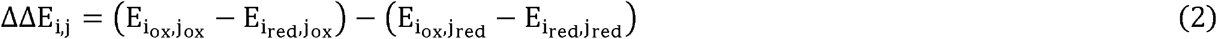

A more negative (positive) ΔEi indicates that the oxidized state is relatively stabilized (destabilized) and correlates with a more negative (positive) redox potential if the electrostatic contribution dominates the free energy of oxidation. In the context of electron transfer, the driving force is related to the difference in ΔEi between donating and accepting groups. Because those groups are chemically identical in the CNs, non-electrostatic contributions are expected to largely cancel.

Eq. 1 was used previously to estimate driving forces in OmcS;^53^ Eq. 2 is an extension of the idea to evaluate heme redox cooperativities as defined in.^54^

Consistent with this approach, electrostatic interactions were shown to dominate the redox potentials in Omc-E, S, and Z.^41^ Furthermore, the implementation of Eqs. 1 and 2 successfully reproduced the variation of redox potentials for hemes, with and without anti-cooperative interactions, in aqueous and membranous environments.^55^

To evaluate each energy term, BioDC interfaces with the Poisson-Boltzmann Solvation Area (PBSA) model of the AmberTools suite.^56^ The external dielectric constant was set to 78.2 for a bulk aqueous environment at 25°C and the implicit salt concentration was set to a physiologically relevant 0.15 M.

The internal dielectric constant of the protein was estimated from the total solvent accessible surface area (SASA) for each donor-acceptor pair by Eq. 3,

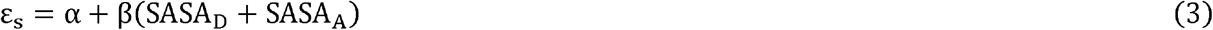

where α = 5.18 and β = 0.016 Å^-2^. These parameters were previously developed^57^ so that the Marcus continuum equation (Eq. 4 below) reproduces reorganization energies from MD simulations performed with a polarizable forcefield.

When computing heme redox anti-cooperativities by Eq. 2, the estimation of the interior dielectric constant was simplified in view of the fact that calculations were needed for every possible pair of hemes (not just adjacent donor-acceptor pairs). The internal dielectric constant was set for all heme pairs as the average of the values computed by Eq. 3 for adjacent heme pairs.

### 2.3. Reorganization Energy

#### 2.3.1. Empirically Parameterized Marcus Continuum Approach

BioDC estimates the outer-sphere reorganization energy (λ_out_) by assuming Δq units of charge are transferred between spherical donor and acceptor groups (radii R_D_ and R_A_) at an R center-to-center distance in a bulk medium characterized by static (ε_s_) and optical (ε_opt_) dielectric constants. The expression, Eq. 4, is known as the Marcus continuum equation.

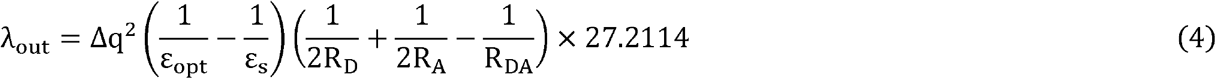

Eq. 4 was shown before^57^ to well reproduce λ_out_ from all-atom MD simulations conducted with an electronically polarizable forcefield if R_D_ = R_A_ = 4.6 Å for an effective heme radius, ε_opt_ = 1.84, and ε_s_ is assumed to be a linear function of the total (donor + acceptor) SASA (Eq. 3),

#### 2.3.2. Vertical Energy Distributions and Polarizability Corrections

The thermally-sampled distributions of instantaneous (vertical) changes in electrostatic interaction energy (ΔU) between the donor or acceptor and the rest of the environment upon oxidation/reduction The thermally-sampled distributions of instantaneous (vertical) changes in electrostatic interaction were used in a second approach to compute λ_out_.

If ΔU for the reactant (R) and product (P) states is Gaussian distributed and ergodically sampled according to Boltzmann statistics on the timescale of the electron transfer, λ_out_ is equivalently defined in terms of the average (⟨ΔU⟩_x_,x = R or P; Eq.5) or variance (σ_x_; Eq.6) of the vertical energy distributions in the two states. These definitions are respectively known as the Stokes-shift (λ^st^)and variance (λ^var^) reorganization energies; if they are not equivalent, the reaction reorganization energy (λ^rxn^) that should be used in Marcus theory is defined in terms of their ratio (Eq. 7).^58^ In Eqs. 5–7, k_b_ and T are respectively the Boltzmann constant and absolute temperature.

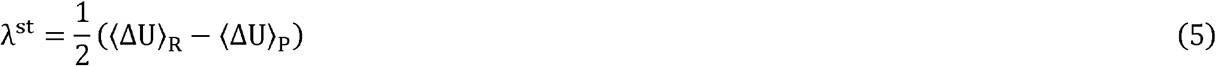

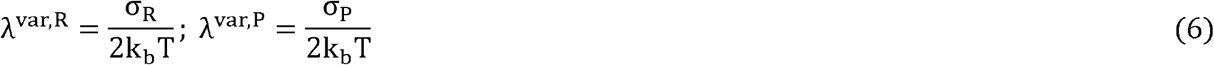

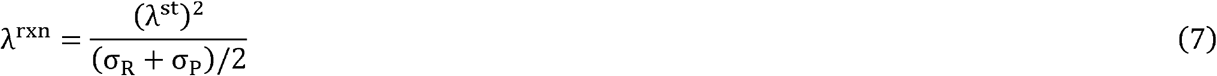

Because the inner-sphere contribution is much less than the outer-sphere contribution for heme groups, λ^rxn^ ≈ λ_out_ in this work. However, it will be noted in the results when λ^rxn^ is different from Because the inner-sphere contribution is much less than the outer-sphere contribution for heme the expectations of standard Marcus theory because λ^st^ ≠λ^var,R/P^.

##### 2.3.2.1. Coulombic Vertical Energies

ΔU is typically computed as the instantaneous Coulombic energy change (ΔU_coul_) for the donor-acceptor pair in the R or P state from classical MD simulations conducted using non-polarizable forcefields. ΔU_coul_ is given by the product of the change in partial atomic charge for each atom of the donor and acceptor (Δ_q_) with the electrostatic potential (ϕ) exerted by the environment on that atom, summed over all the atoms of the two groups (Eq. 8).

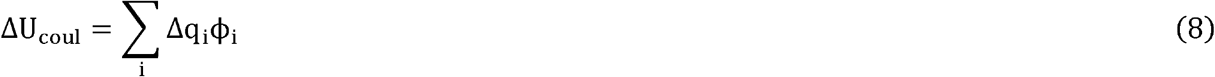

ΔU_coul_ was computed with the CPPTRAJ program^59^ of the AmberTools suite; an example CPPTRAJ input script is provided with the PolReorg package. Plots showing the distributions of ΔU_coul_ in the R and P states, as well as the convergence of λ^st^, λ^var,R^, and λ^var,P^, are shown for each electron transfer in Omc-E, S, and Z in Supplementary Figures 1–3.

##### 2.3.3.2. Perturbative Correction for Active Site Polarizability

ΔU was alternatively expressed for the R or P state by perturbation theory (Eq. 9) as the sum of a reference energy (ΔU_0_), the Coulombic term (ΔU_Coul_; Eq.8i, and the polarization energy (ΔU_pol_ ; Eq.10).^58^

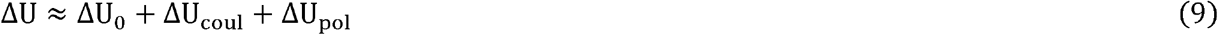

ΔU_pol_ for the donor or acceptor is related to the difference in the second-rank polarizability tensor between the oxidation states for that group (Δ **α**) and the electric field vector (**E**) acting on it (Eq. 10), which is approximated according to literature precedent^60, 61^(although over-simplistically) as the field acting on the Fe center of each heme. Note that because Δ always means final – initial, Δ **α** = (α^ox^− α^red^), where n = 1 for the donor and -1 for the acceptor.

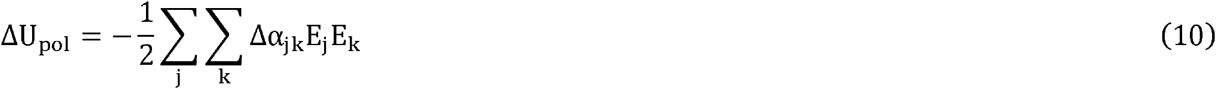

To evaluate Eq. 10, Δ **α** was computed with a variety of model chemistries for a His-His ligated heme group in both vacuum and configurations of the distinct binding sites of the OmcS filament sampled during MD simulations. To facilitate a comparison with prior work, Δ**α** was also computed for a His-Met ligated heme group in vacuum and configurations of cytochrome *c* from MD simulations. The benchmarking data is in Supplementary Tables 1 and 2.

The electric field at the heme-Fe centers was computed over the course of MD trajectories by the EFieldAnalysis program. EFieldAnalysis implements the methodology for a probe atom found in the TUPÂ package,^62^ but unlike TUPÂ, EFieldAnalysis wraps molecules instead of atoms at the periodic boundary to preserve molecular integrity. It also parallelizes the calculation of the total field vector over multiple CPUs for a more efficient assessment of long trajectories. Eqs. 9 and 10 were implemented in the PolReorg python package.

### 2.4. Electron Transfer Rates, Charge Diffusion Constants, and Steady-State Fluxes

Non-adiabatic Marcus theory electron transfer rates were computed with BioDC and used with (1) the analytical Derrida equation to obtain a charge diffusion constant,^63^ and (2) a multi-particle steady-state kinetics model to obtain the protein-limited electron flux.^53, 57, 64^ These two kinetic models are implemented in the ETAnalysis program with code graciously contributed by Fredrik Jansson (Derrida module) and Jochen Blumberger and Xiuyun Jiang (flux module). The latter module was modified to read the energy matrix of site and site-site interaction energies calculated by BioDC. With that information, it self-consistently updates the redox potentials by weighing the interactions according to the electron occupancies. The self-consistent procedure was improved by implementing an adaptive mixing scheme to aid convergence.

### 2.5. Monte Carlo Sampling of Energetic Parameters for Target Charge Diffusion Constants

The Parameter Explorer module of ETAnalysis was used to sample sets of electronic couplings, reaction free energies, and reorganization energies for a specified sequence of slip- and T-stacked heme pairs. A set is accepted if it gives a desired charge diffusion constant within a specified tolerance (here ±10%). A KD-tree algorithm was used to remove duplicate sets of parameters.

### 2.6. UV-vis Spectral Analysis

#### 2.6.1. Structure Preparation

The model of a heme cofactor shown in Supplementary Figure 4 was prepared by (1) cleaving and capping with hydrogens the C_β_-C_α_ bonds that connect the cofactor to the protein via axial His ligands and thioether bonds with Cys residues; (2) removing the propionic acid groups and treating them as part of the electrostatic environment,^65, 66^ and (3) for some computations, methyl substituents were replaced with hydrogen atoms. The latter truncation had a minimal spectral effect (Supplementary Figure 5).

The heme model was optimized in the closed-shell singlet reduced (formally Fe^2+^) state with the Becke, three-parameter, Lee–Yang–Parr (B3LYP) density functional^67, 68^ and a mixed basis set comprised of LANL2DZ for the Fe center^69^ and 6-31G(d) for H, C, N and S^70^). A local minimum was found, as confirmed by a harmonic vibrational analysis (*i*.*e*., no negative frequencies).

#### 2.6.2. Electrostatic Contribution to Spectral Tuning

The unmethylated heme model was superimposed on and replaced each of the six hemes in the central subunit of an OmcS trimer. The superpositions were performed on 216–300 configurations of the protein (sampling rate = 1 frame/ns) generated by previously published MD simulations^41^ in which the target heme was reduced while all other hemes were oxidized. This procedure leveraged the semi-rigid nature of the heme group to preserve the influence of the environment while avoiding the unreliability of MD-generated geometries in quantum mechanical calculations, as well as the intractability of 10^3^ quantum mechanical/molecular mechanical optimizations or dynamical simulations.

Time-dependent (TD)-DFT calculations were performed on these configurations with the superimposed hemes in the QM region and all other atoms converted to an electrostatic background. The CAM-B3LYP functional is known to produce a blue-shifted absorption spectrum for heme,^71^ and B3LYP does the same (Figure S6). On the hypothesis that the blue-shifting results from too much electron localization, a functional with a smaller fraction of Hartree-Fock exchange (specifically BLYP) was tested and found to give closer agreement with the expected position of the Soret absorption maximum (∼420 nm) for a reduced heme (Figure S6). The same mixed basis set used for geometry optimization was used for the spectral simulations.

#### 2.6.3. Conformational Contribution to Spectral Tuning

The heme cofactor was displaced in 0.1 Å increments from 0.0–1.0 Å along each of the ruffling and saddling deformation normal coordinates^72^ and the absorption spectrum was simulated for each conformer. The displaced geometries were prepared with the Normal Coordinate Structure program graciously provided by Christopher J. Kingsbury.^72^ The structures were optimized with all dihedrals fixed before simulating the spectra with TD-DFT.

#### 2.6.3. Excitonic Contribution to Spectral Tuning

The Frankel exciton model^73, 74^ implemented in Gaussian 16 Rev. A.03 was used to compute the UV-vis spectrum and to quantify excitonic couplings. For this calculation, the heme with methyl substituents was superimposed on the hemes in a configuration of the protein generated by MD and each heme was defined as a separate QM region. The protein-water environment was again included as an electrostatic background.

The calculation was performed with CIS and a mixed (LANL2DZ for Fe; 6-31G(d) for H, C, N, and S) basis set, as has been done previously^75^ to avoid the deficiencies of TD-DFT.^76^

## 3. Results and Discussion

Biological electron transfer in multiheme proteins is generally pictured in terms of a succession of reduction-oxidation (redox) reactions in which the electron takes multiple, incoherent ‘hops’ from site-to-site.^77^ In this view, an electron resides on a heme group until an infrequent thermal fluctuation overcomes the energy barrier for the electron to be on the adjacent heme, at which time the electron ‘hops’ to that heme with a probability less than unity (*i*.*e*., non-adiabatically).

The activating thermal fluctuation is rare because the energy barrier is usually several multiples of the available thermal energy. In the time spent waiting for the energetically propitious configuration to be realized, the electron transferred in the prior ‘hop’ is thermally equilibrated and loses coherence. Once the barrier for the next ‘hop’ is overcome, the probability for transferring the electron is small, because only the tails of the wavefunctions on adjacent hemes overlap in the intervening space due to the exponential decay of the wavefunctions with distance from the edge of the macrocycle.

The rate constant for each electron transfer (k_et_) is therefore the product of probabilities for thermal activation (ρ) and electronic tunneling (κ) which, in the high temperature (classical) limit, takes the form of standard Marcus theory:^78, 79^

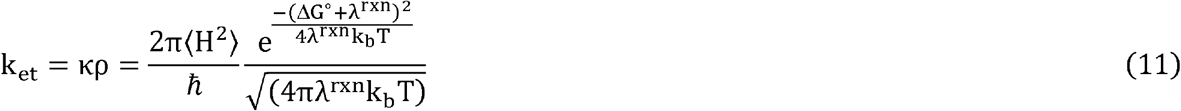

Apart from fundamental constants (*ћ*= reduced Plank constant; k_b_ = Boltzmann constant), k_et_ depends on three energetic parameters: (1) the thermally averaged electronic coupling squared (⟨H^2^⟩), which describes the overlap of wavefunctions on the donor and acceptor, (2) the reaction reorganization energy (λ^rxn^), which describes the energy penalty for rearranging the nuclei to be consistent with the product state before the electron is transferred, and (3) the reaction free energy (ΔG°), which is given by the redox potential difference between the charge donor and acceptor. The term 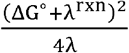 is defined as the activation energy (E_a_).

Each of these energetic terms is analyzed in turn for all structurally characterized CNs (Sections 3.1–3.4). The associated rates (Section 3.5) are assembled into diffusive and steady-state kinetic models of long-range electron transfer (Section 3.6). The computed conductivities are compared to solid-state electrical measurements and the demands of cellular respiration (Section 3.6).

### 3.1. Electronic versus Excitonic Coupling

Electronic coupling strengths for the heme packing geometries present in CNs have extensively and consistently been found to be sub-thermal energy at 300 K.^77^ And yet, UV-vis spectra for multiheme proteins were recently interpreted as suggesting an electronic band structure.^80^ This section clarifies that quantum mechanical electronic couplings in CNs are weak, whereas excitonic couplings are almost entirely classical and 10-fold larger. The excitonic couplings give insight into multiheme redox anti-cooperativities, which are important later.

#### 3.1.1. Heme-Heme Electronic Coupling Strengths are Sub-Thermal Energy

Electronic couplings are principally determined by the distance and orientation of the charge donating and accepting groups.^81^ The electrostatic environment causes only a ∼10% perturbation.^43, 82^

Adjacent hemes in the known CNs adopt two highly conserved packing geometries found throughout the class of multiheme proteins,^27, 83^ and preserved under hundreds of nanoseconds of MD simulations.^84^ These packing geometries are the parallel-displaced (“slip-stacked”, denoted “S”) and perpendicular (“T-stacked”, denoted “T”) motifs (Figure 2A).

**Figure 2.**
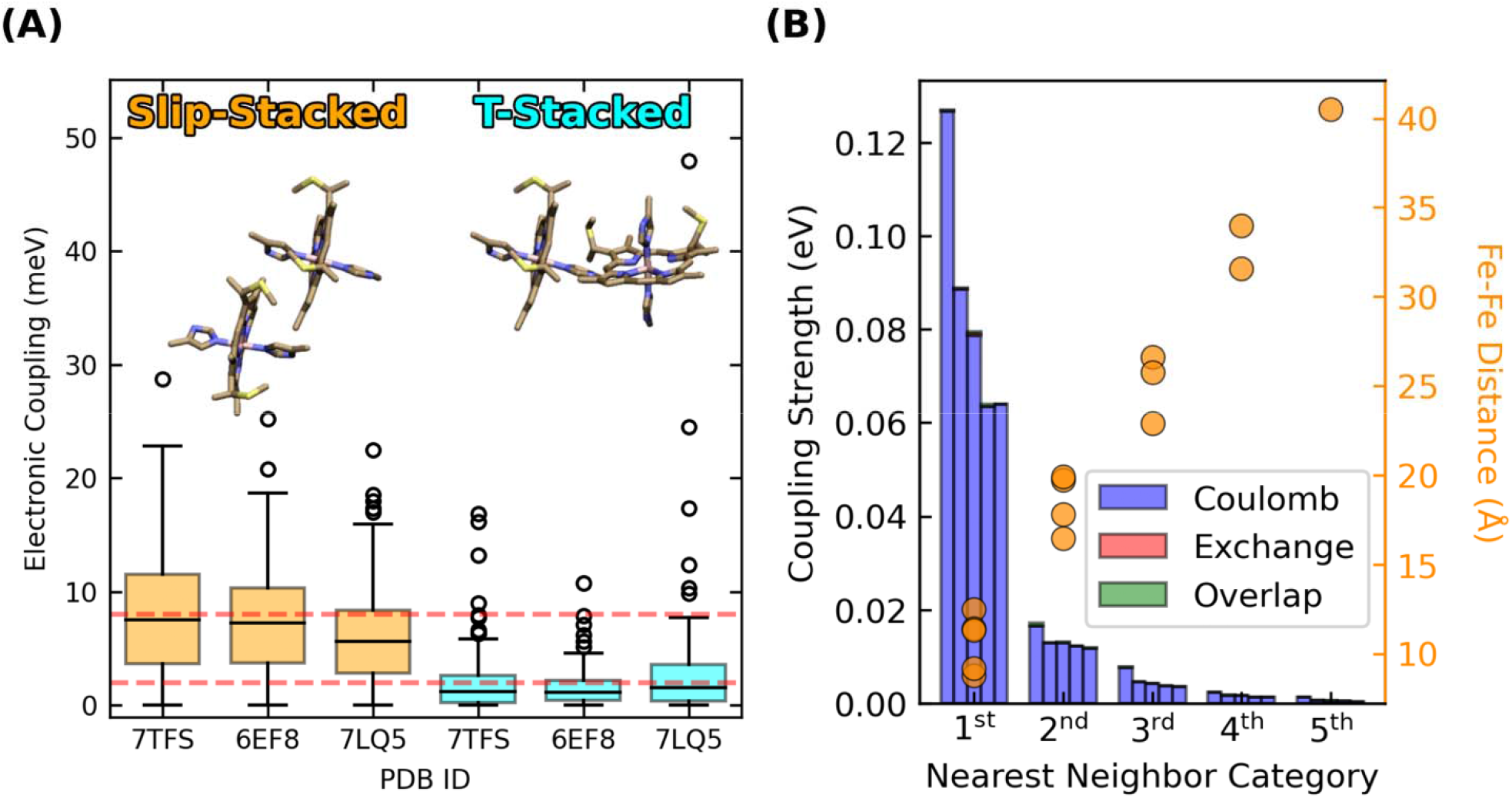
Slip- and T-stacked hemes in the multiheme architecture of cytochrome ‘nanowires’ have weak quantum mechanical electronic couplings but stronger classical excitonic interactions. **(A)** Electronic couplings in Omc- E, S, and Z. The data is reproduced from.^41^ **(B)** Excitonic couplings computed for the hemes in OmcS at the CIS/[LANL2DZ(Fe):6-31G(d)(H,C,N,S)] level of theory.

Four of the five CNs (OmcE, A3MW92, F2KMU8, and OmcS) exclusively have a (TS)_n_ pattern, whereas OmcZ has a (TSSTSTS)_n_ pattern along the main chain, as well as a heme that branches from the main chain.

Root-mean-squared electronic couplings 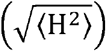 previously computed for Omc-E, S, and Z were 0.007–0.010 eV for slip-stacked and 0.002–0.006 eV for T-stacked geometries (Figure 2A).^41^ These values are in good agreement with other studies on OmcS^47, 53^ and multiheme cytochromes in general.^77^ Given the conserved geometries and coupling ranges, values of 0.008 and 0.002 eV for slip- and T-stacked heme pairs (red dashed lines in Figure 2A) are adopted to analyze the electron flux through all CNs in the present work.

#### 3.1.2. Geometric Constraints on Heme Couplings Make Electron Transfers Sub-Optimal

T-stacked hemes intrinsically have lower electronic couplings compared to slip-stacked hemes (Section 3.1.1). The consequence, according to prior experiments^85, 86^ and the computations of Section 3.5 (*vide infra*) is ∼10-fold slower electron transfer rates. If T-stacked hemes are a ‘speed-bump’, why are ∼50% of all heme pairs in CNs of this rate-throttling variety?

To transfer electrons, hemes must be placed within tunneling distance (≤14 Å). The closer the hemes are packed, the more readily endergonic steps can be tolerated and traversed.^32^ Close proximity improves wavefunction overlap for electron transfer, but even for hemes in van der Waals contact, electronic couplings are ≤0.01 eV (Figure 2A).

At the same time, an entropic penalty for organizing densely packed hemes in a protein interior must be offset. Thioether linkages between Cys residues and the hemes are perhaps needed, *inter alia*,^87^ to oppose entropically-driven dissociation;^88^ multi-heme proteins with more than three hemes exclusively have *c*-instead of *b*-type hemes,^89^ where the difference is the presence of the thioether linkages.

These linkages work in concert with axial ligands donated by other residues to the heme-Fe centers to place geometrical constraints on the packing geometries. Most notably, the slip-stacked geometry that is most favorable for heme-to-heme electron transfer cannot be physically accommodated for more than two consecutive heme pairs by the protein backbone to which the hemes are tethered.^8^ The geometrically acceptable possibilities include a continuous chain of T-stacked hemes,^8^ which is not observed, or the typical alternating pattern of parallel and perpendicularly-stacked hemes, as found in the CNs. Either way, the rate-throttling T-stacked motif is *unavoidable* in a cytochrome with more than three *c*-type hemes, and cytochromes with this many hemes only have the *c*-type variety.

Electron transfer through multi-heme cytochromes is physically constrained by the geometry of their construction. The implication is that CNs are *not* optimized (in an absolute sense) for long-range electron transfer. However, Section 3.6 will show the conductivity to be ‘good-enough’ for their proposed role in cellular respiration.

#### 3.1.3. Multi-Heme UV-vis Spectra Misinterpreted as Evidence for Electronic Band Structure

If electronic couplings in CNs, and multihemes more generally are weak, how can the appearance of a single Soret band in their UV-vis spectra be understood? Multihemes were proposed to have an electronic band structure to explain why the hemes do not have distinct absorption signatures.^80^ Weakly coupled hemes under physiological conditions, however, cannot support long-range delocalization. What, then, is the origin of the single Soret band, for example, in OmcS?^90^

The key quantity to consider is excitonic, as opposed to electronic coupling. Figure 2B shows the five largest heme-heme excitonic couplings between excited states for all possible pairs of hemes in a subunit of OmcS, grouped by their nearest-neighbor relationships (1^st^ through 5^th^).

Heme-heme excitonic couplings are almost entirely Coulombic in nature and exceed electronic couplings by ∼10-fold for the same pairs of adjacent (1^st^ nearest neighbors) hemes with Fe-Fe distances of 9–12 Å. Longer range interactions are comparable to thermal energy.

These observations follow from the distinct physical origins of electronic and excitonic couplings: The electronic couplings in Figure 2A require orbital overlap, decay exponentially with distance, and only involve the tails of the donor and acceptor wavefunctions. The excitonic couplings in Figure 2B do not require orbital overlap, involve interactions of the full transition dipole moments on the two hemes, and decay with a slower r^-3^ dependence. Thus, Figure 2 indicates that hemes in CNs are weakly electronically coupled because of poor wavefunction overlap in highly conserved packing geometries, but more strongly excitonically coupled because of Coulombic interactions at short distances.

Though excitonic couplings are stronger than electronic couplings, their presence is still not spectrally resolvable for the hexaheme homopolymer OmcS. The 422-nm Soret band for OmcS has a full-width-at-half-maximum that is ∼36 nm,^90^ only twice the full-width-at-half-maximum for a single heme in solution or bound to a globin.^91, 92^ This observation can be understood by considering the factors that can separate the Soret absorptions from multiple hemes.

Figure 3A shows that Gaussian-broadened spectra simulated within a Frankel exciton model are very nearly identical when none, only Coulombic, or all (Coulombic, exchange, and overlap) site-site interactions for the hemes in OmcS are included. Displacements along the two most populated normal coordinate deformations of the heme macrocycles in OmcS^41^ only cause an ∼8 nm variation in the Soret absorption maximum (Figure 3B). A similarly small spread is produced by the distinct binding-site electrostatics of the protein-water environment (Figures 3C). Even if these tuning effects are additive, the spread in absorption maxima of the individual hemes would be within the envelope of the overall Soret band.

**Figure 3.**
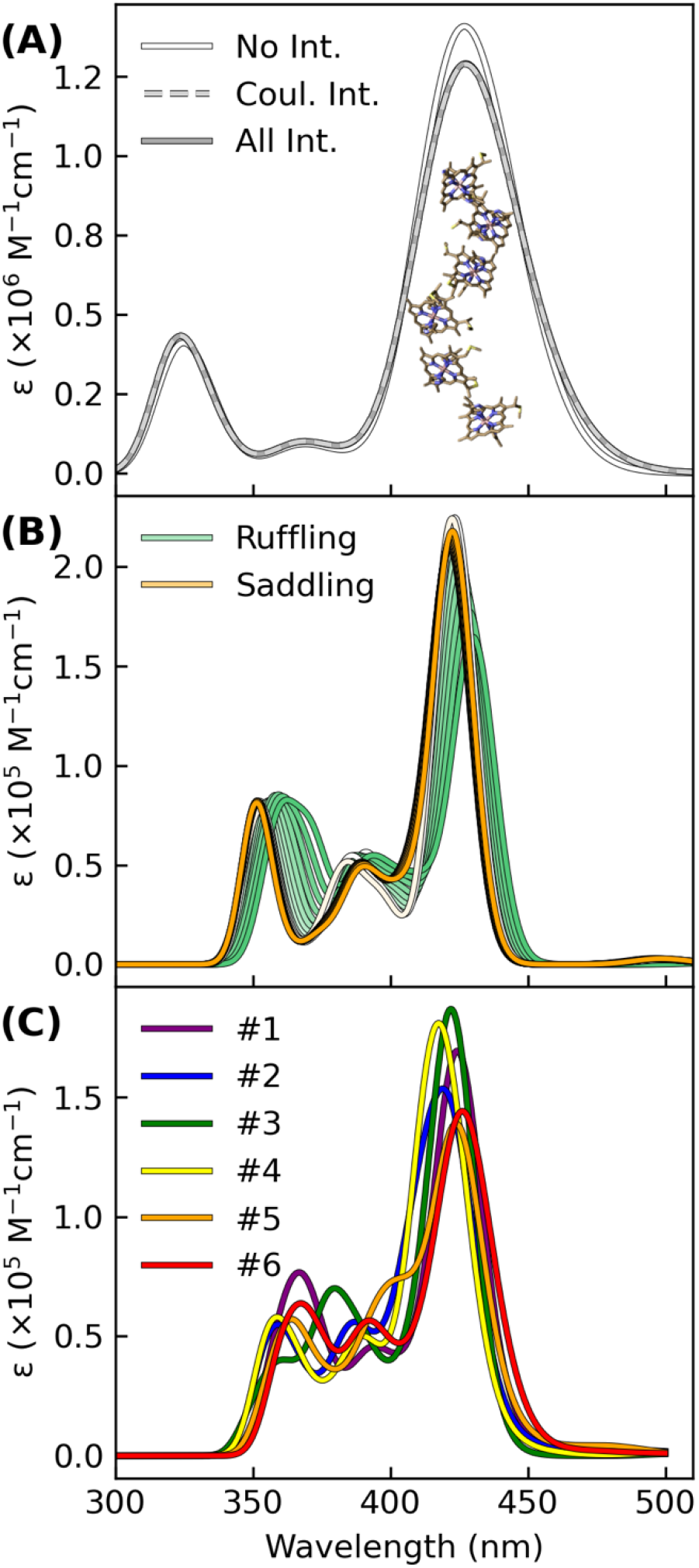
Excitonic, conformational, and electrostatic factors minimally distinguish the spectral signatures of each heme in OmcS, thereby giving a Soret band that is a superposition of nearly coincidental absorptions. **(A)** Spectra simulated for the shown hexaheme array with none, only Coulombic, or all (Coulombic, exchange, and overlap) site-site interactions. **(B)** Spectra simulated for the heme group systematically displaced by 0.1 increments from 0.0 to 1.0 Å along the ruffling and saddling normal coordinate deformations that were previously found to be most populated by the hemes in OmcS.^41^ (C) Thermally averaged spectra of each heme (#1–#6) within the central subunit of an OmcS trimer. **(A–C)** The spectra were simulated using either CIS (panel A) or TD-BLYP (panels B and C) with a mixed basis set [LANL2DZ(Fe):6-31G(d)(H,C,N,S)] and uniformly shifted to align the Soret maximum to the experimental 420 nm position. Gaussians were fit to each transition with a broadening factor of 0.05 eV to simulate the lineshapes.

Taken together, conformational, excitonic, and electrostatic interactions have a fairly minor influence on differentiating the absorption signatures of each heme in OmcS, and presumably other multiheme cytochromes. Instead of all hemes having a single absorption because of electronic delocalization, the separate hemes have nearly coincidental absorptions that are spectrally unresolved. The single Soret band for a multiheme is not inconsistent with weakly electronically coupled hemes.

### 3.2. Redox Potentials, Anti-Cooperativities, and Reaction Free Energies

Only homo, as opposed to heteropolymeric CNs have so far been discovered. Homopolymerization has the interesting consequence that the overall intra-filament driving force is zero: The rise and fall in free energy for an electron through a subunit must repeat when the electron gets to the next subunit.

The driving force between the redox half reactions connected by the CNs is expected to be ∼0.1 V.^93^ Even if the potential falls off linearly across the micron-long filament, the contribution to the energetics at the level of heme-to-heme electron transfers is less than thermal energy.^94^ Furthermore, because the electron transfers are between chemically identical bis-histidine-ligated *c*-type hemes, each reaction is a self-exchange, and there is formally no driving force.

The heme-to-heme driving forces are not rigorously zero, however, because the redox potential of each heme is differently affected by the heterogeneity of the protein-water environment.^41^ The redox potentials are also electrostatically influenced by the oxidation state of all other hemes in an effect known as redox anti-cooperativity:^95^ The oxidation of heme *i* is disfavored by the oxidation of heme *j* due to the electrostatic repulsion that would result between like charges. A positive cooperativity may result from redox-linked conformational transitions.^95, 96^

The following sub-sections delineate tradeoffs and physical constraints for driving forces that are encoded in the multiheme architecture of CNs; namely: (1) The dense packing of hemes causes electrostatic microenvironments to be shared and redox potential differences between adjacent hemes in van der Waals contact to be small; and (2) Dense packing also causes strong (∼0.1 eV) redox anti-cooperativities, but the linear topology of CNs makes the ΔG° landscape largely insensitive to these heme-heme interactions.

#### 3.2.1. Shared Electrostatic Microenvironments Limit Driving Forces in Multihemes

The potential of a redox transition in a multi-center redox protein can be expressed as the sum of site energies and site-site interaction energies, which are respectively represented by the diagonal and off-diagonal elements of an energy matrix.^54^ Figure 4 shows the energy matrix computed with BioDC for the tetra- (OmcE, A3MW92, and F2KMU8), hexa- (OmcS), and octaheme (OmcZ) CNs. The figure also compares the energy matrices for the two CryoEM models of OmcS (PDB 6EF8 and 6NEF) and OmcZ (PDB 7LQ5 and 8D9M), as well as the results for both of these proteins to a prior quantum mechanics/molecular mechanics (QM/MM) investigation^41^ that *assumed* no heme-heme interactions.

**Figure 4.**
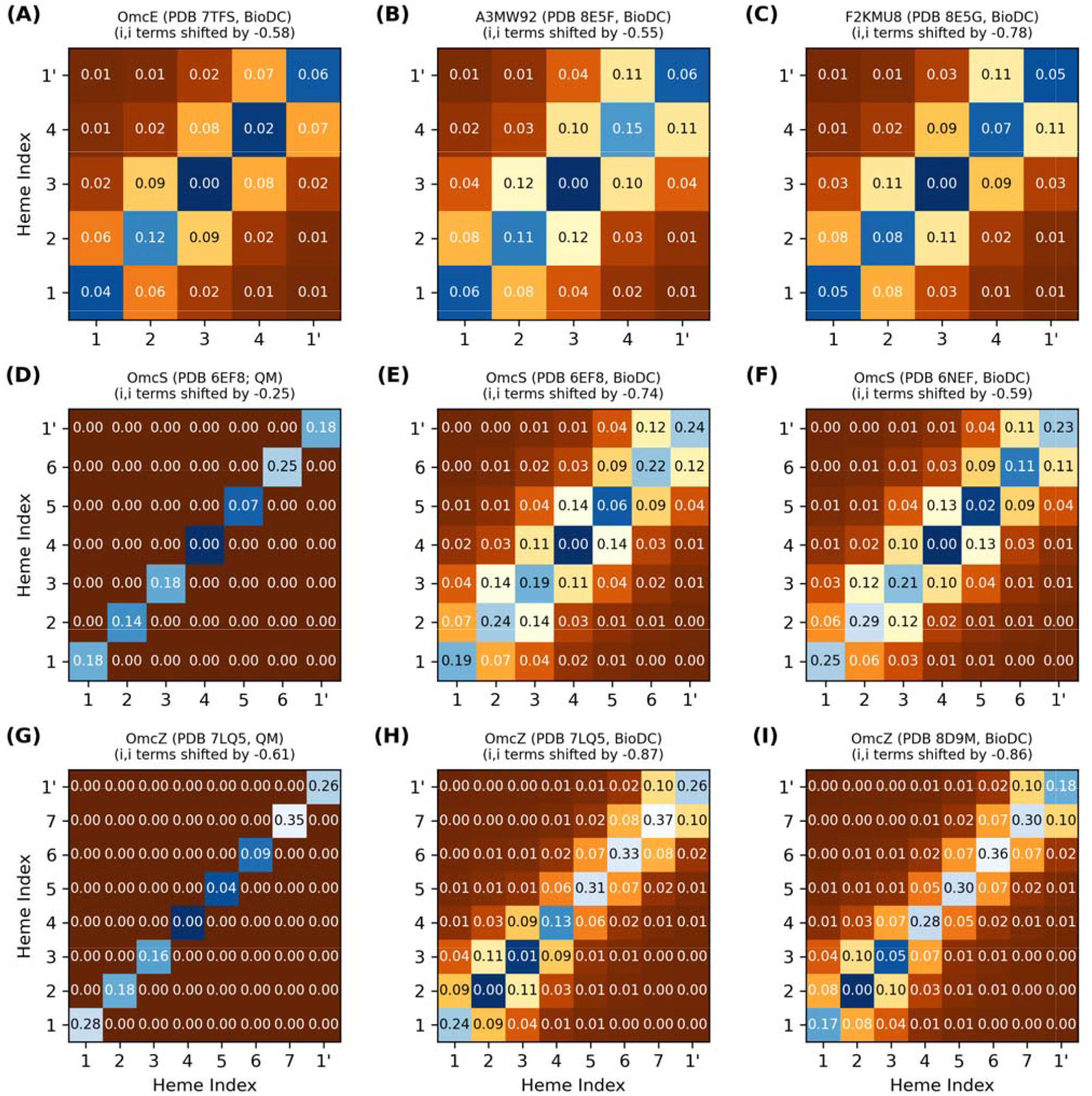
Thermodynamic characterization of relative heme oxidation energies (diagonal elements) and heme-heme interaction energies (off-diagonal elements) in the structurally characterized cytochrome ‘nanowires.’ All energy matrices were computed with BioDC, except panels **(D)** and **(G)** which are from a prior QM/MM investigation^41^ that *assumed* all off-diagonal elements were zero. For each matrix, the hemes are indexed in their linear sequence from 1 to N, and “1’” indicates the first heme of the next subunit. All energies are in eV.

The thermodynamic description of a multiheme protein, let along all structurally characterized CNs, is extremely demanding computationally with quantum mechanics^96^ and experimentally requires advanced nuclear magnetic resonance techniques that have yet to be extended to a system with as many redox centers as the CNs.^97^ Figure 4 therefore showcases the insights currently only available via the systematic series of calculations implemented in BioDC using the PBSA method.

The diagonal site energies in Figure 4 are expressed relative to the most negative site, because only the *differences* between sites matter for the energetics of electron transfer. More positive values on the diagonal indicate an electrostatic destabilization of the oxidized versus reduced state and should correlate with higher (more positive) redox potentials.

The relative site energies are predicted by both QM/MM and/or BioDC to be ≤0.15 eV for the tetrahemes, ≤0.25 eV for the hexaheme, and ≤0.37 eV for the octaheme. For several other multiheme cytochromes studied to date, ΔG° has also been found to be ≤|0.3| eV.^53, 64, 98^Site energy differences are kept relatively small because residues that electrostatically shift the redox potential of one heme also shift the redox potential of the adjacent heme in van der Waals contact to a similar extent and in the same direction.

#### 3.2.2. Adjacent Heme-Heme interactions are ≤0.12 eV

Close-packing, however, introduces heme-heme redox anti-cooperativities. Figure 4 shows that the off-diagonal terms for first-nearest neighbor hemes are predicted to be 0.05–0.14 eV for all CNs, whereas more distant interactions are comparable to thermal energy. Several lines of evidence support this finding.

1. BioDC previously described quantitatively the role of heme-heme interactions in water-soluble as well as membrane-embedded *de novo* designed heme protein maquettes.^99^
2. Prior Constant pH and Redox Molecular Dynamics (C(E,pH)MD) simulations^43^ found interaction strengths of ∼0.08 eV by noting the difference in potentials when hemes were titrated simultaneously versus independently with all other hemes held in the oxidized state.
3. Excitonic couplings between hemes computed at the QM/MM level (Figure 2B) are almost entirely Coulombic and agree quantitatively with the range of redox anti-cooperativities computed with classical electrostatics. These two calculations model distinct physical processes where the fate of the electron is different, but both create a positive hole that electrostatically interacts with the neighboring heme. The similarity of this electrostatic interaction is why excitonic couplings and redox anti-cooperativities are of the same magnitude. Oxidation, in a sense, is the limiting extent of a photo-excitation.
4. A long established Debye-Hückel shielded electrostatics (DHSE) model^95^ predicts similar, albeit smaller, interaction energies of 0.03–0.06 eV for the Fe-to-Fe distances found in the CNs. However, this model assumes an effective dielectric constant of 8.6, whereas the dielectric constants of the heme binding sites in the CNs are estimated by Eq. 3 to be 5.2–7.6 (Section 2.2); a previous study found dielectric constants of 3–7 for OmcS.^43^ A smaller dielectric constant means electrostatic interactions are less screened, and the interaction energies should therefore be larger, as found here.
5. A recent spectroelectrochemical experiment on OmcS was *interpreted* to mean heme-heme interactions are negligible,^100^ but for hemes in van der Waals contact, this interpretation is physically unreasonable and relies on a fundamental misunderstanding of statistical mechanics.

As the solution potential is swept in a spectroelectrochemical experiment, a multi-center redox protein passes through a series of macroscopic oxidation stages.^101^ Each stage comprises an ensemble of microstates that describe all possible ways of distributing reducing equivalents among the redox centers. The potentials at which these macroscopic transitions occur are observed by spectroelectrochemistry. But from those potentials, it is not possible to deduce anything about the statistical weights of the microstates that produced the bulk-averaged observable. The finding that a model of independent Nernstian centers fits the titration curve does not rule out the presence of heme-heme interactions any more than the equally good fit of a model with sequentially-coupled Nernstian centers confirms their presence.

Furthermore, the prior report^100^ used a 6-parameter fit of independent or sequentially-coupled redox reactions to account for the titration curve without any sensitivity analysis to guard against overfitting. A re-analysis of the titration curve for OmcS (Supplementary Figure 7) indicates that a model of only 3 hemes is sufficient to fit the curve according to either the Akaike or Bayesian information criterion, which balances model fit against model complexity, or even the plateauing of the root-mean-squared-error of the fit with the increasing number of hemes in the model.

At the low resolution of the smooth experimental titration curve, half (3/6) of the hemes in a subunit of OmcS appear to have potentials indistinguishable from the potentials of the other hemes, and the full range of potentials is only ∼0.2 V. The small differences and overall range are in good agreement with the prediction of relative site energies in Figure 4.

Worth noting in passing because the original authors did not: The spectroelectrochemical data definitively invalidates their prior models of why OmcS is more conductive upon cooling or chemical reduction. Malvankar and co-workers predicted a massive hysteresis between cathodic and anodic directions (a 0.4 V shift in midpoint potential), as well as potentials as low as -0.826 V versus standard hydrogen electrode (SHE).^47^ Neither prediction, published in 2022, was born out,^100^ even by earlier spectroelectrochemical data collected in 2020^48^ and 2021^49^ under the supervision of Malvankar.

#### 3.2.3. Linear Filament Topology Blunts the Effect of Redox Anti-Cooperativity on Driving Forces

The relative oxidation energies on the diagonal of each energy matrix in Figure 4 do *not* monotonically increase in the geometric order of hemes #1 to N in the CNs. This observation means that the preferred thermodynamic oxidation sequence is *different* from the oxidation order that must happen as electrons flow linearly along the heme chain. If the oxidation of each heme shifts the potential of the 1^st^ nearest neighbor by ∼+0.1 V because of anti-cooperative interactions, oxidizing the hemes in geometric versus thermodynamic order has a significant consequence (Figure 5).

**Figure 5.**
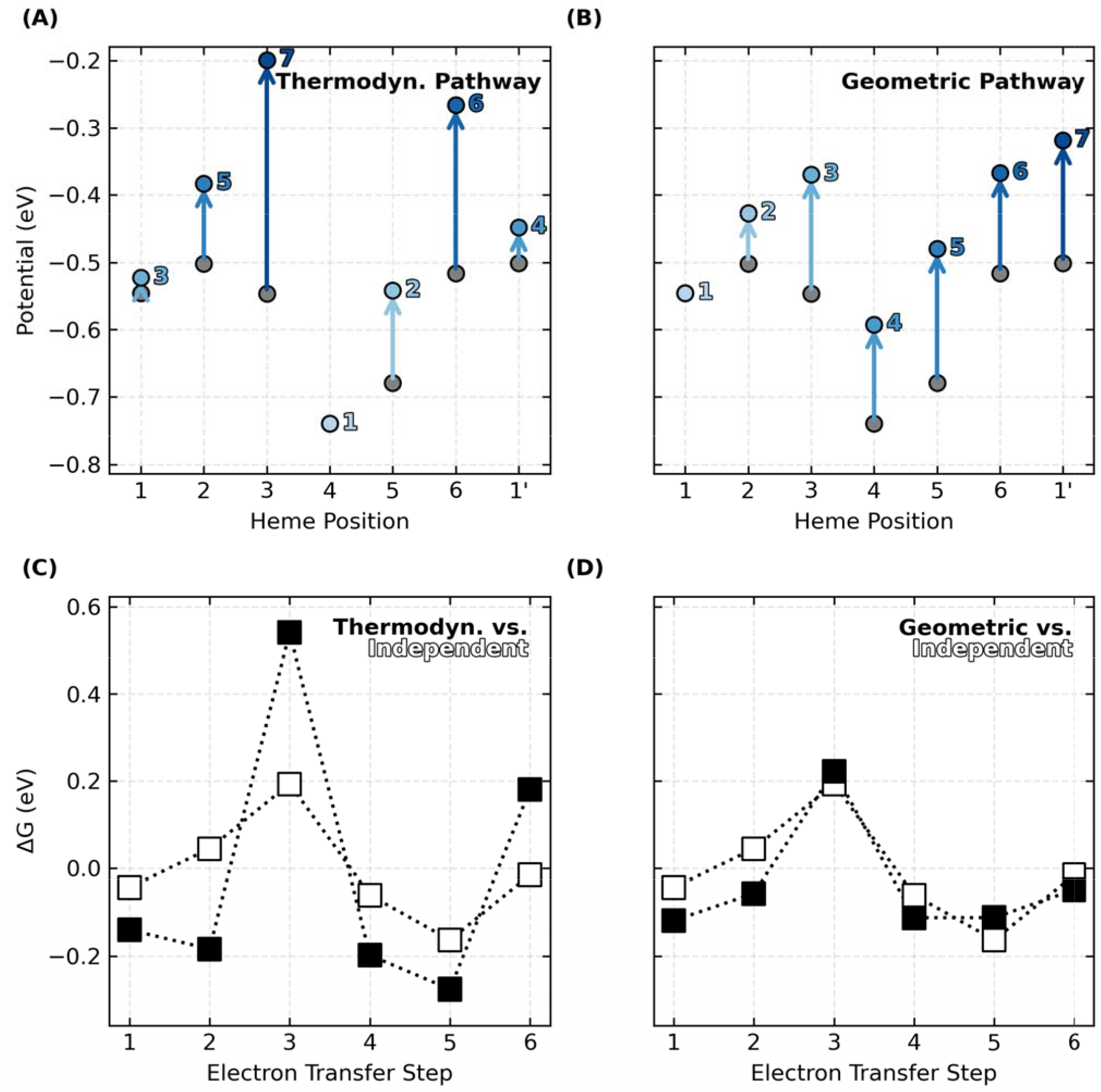
Oxidation of hemes in the linear sequence prescribed by the topology of cytochrome ‘nanowires’, as opposed to the thermodynamically preferred order, blunts the effect of redox anti-cooperativities on the landscape for electron transfer. **(A, B)** Shifts in oxidation energies due to redox anti-cooperativities as the hemes oxidize in order of their thermodynamic preferences or geometric arrangement. The x-axis follows the linear sequence of the filament whereas the light-to-dark blue annotations indicate the oxidation order. **(C, D)** Comparisons of the landscapes for anti-cooperatively interacting hemes that oxidize in thermodynamic or geometric order with the landscape for non-interacting hemes that oxidize independently. The data shown is for OmcS. Analogous figures for the other cytochrome ‘nanowires’ are Supplementary Figures 8–14.

The predicted thermodynamic order for heme oxidations in OmcS, for example, is #4, #5, #1, #1’, #2, #6, and #3 (Figure 5A), where the prime mark indicates the first heme of the next subunit. This ordering (except for the position of Heme #1) matches a prior QM/MM investigation.^41^

But the specific ordering is less important here than the fact that some hemes become oxidized *after* the flanking heme on either side has already been oxidized. The result is that heme-heme interactions compound, and the middle heme oxidizes at a much more positive potential. This is what happens for Heme #3 (oxidizing after Hemes #2 and #4) and Heme #6 (oxidizing after hemes #5 and #1’) in OmcS (Figure 5A). The consequence is that the peaks and valleys of the free energy landscape are accentuated relative to if the hemes oxidized independently with no heme-heme interactions (Figure 5C).

If, instead, hemes oxidize in order of geometric arrangement (Figure 5B), the potential of each and every heme shifts by a similar amount from the preceding oxidation of the heme immediately before it in the sequence. With every heme shifting by roughly the same amount as a cascade of oxidations moves through the linear topology of the CN, the potential differences, and free energy landscape are largely unaffected (Figure 5D).

The same conclusions are reached for all examined CNs (Supplementary Figures 8–14). Thus, by virtue of having a linear topology that constrains the hemes to oxidize in a geometric sequence different from thermodynamic preferences, potential shifts due to heme-heme interactions do not compound, and the free energy landscape for multi-step electron transfer is rendered largely robust to redox anti-cooperativities.

### 3.3. Reorganization Energy

Electron transfer reorganization energy is generally dissected into inner- (λ_in_) and outer- (λ_out_) sphere components. λ_in_ reflects oxidation-state dependent changes in internal coordinates that, for a heme group, contribute 0.05–0.08 eV to the activation barrier.^102, 103^ λ_out_ reflects the energetic penalty for rearranging the environment to accommodate altered charge distributions on the electron donor and acceptor.

To quantify λ_out_, various methods have been proposed that account for the influence of electronic polarizability in different ways and to different extents. Measurements on cytochrome *c* have served as a reference, but experimental values vary by as much as ∼0.5 V depending on conditions.^104, 105^ Different approaches have either reached the same level of agreement^58, 61, 66, 106, 107^ or been judged successful relative to different experimental values.^58, 108^ To clarify matters for CNs and hemoproteins more generally, different approaches for computing λ_out_ are compared for three of the CNs (Section 3.3.1) before one of the methods is used to compare the full set of CNs (Section 3.3.2). The results are summarized in Figure 6 and Supplementary Tables 3–5.

**Figure 6.**
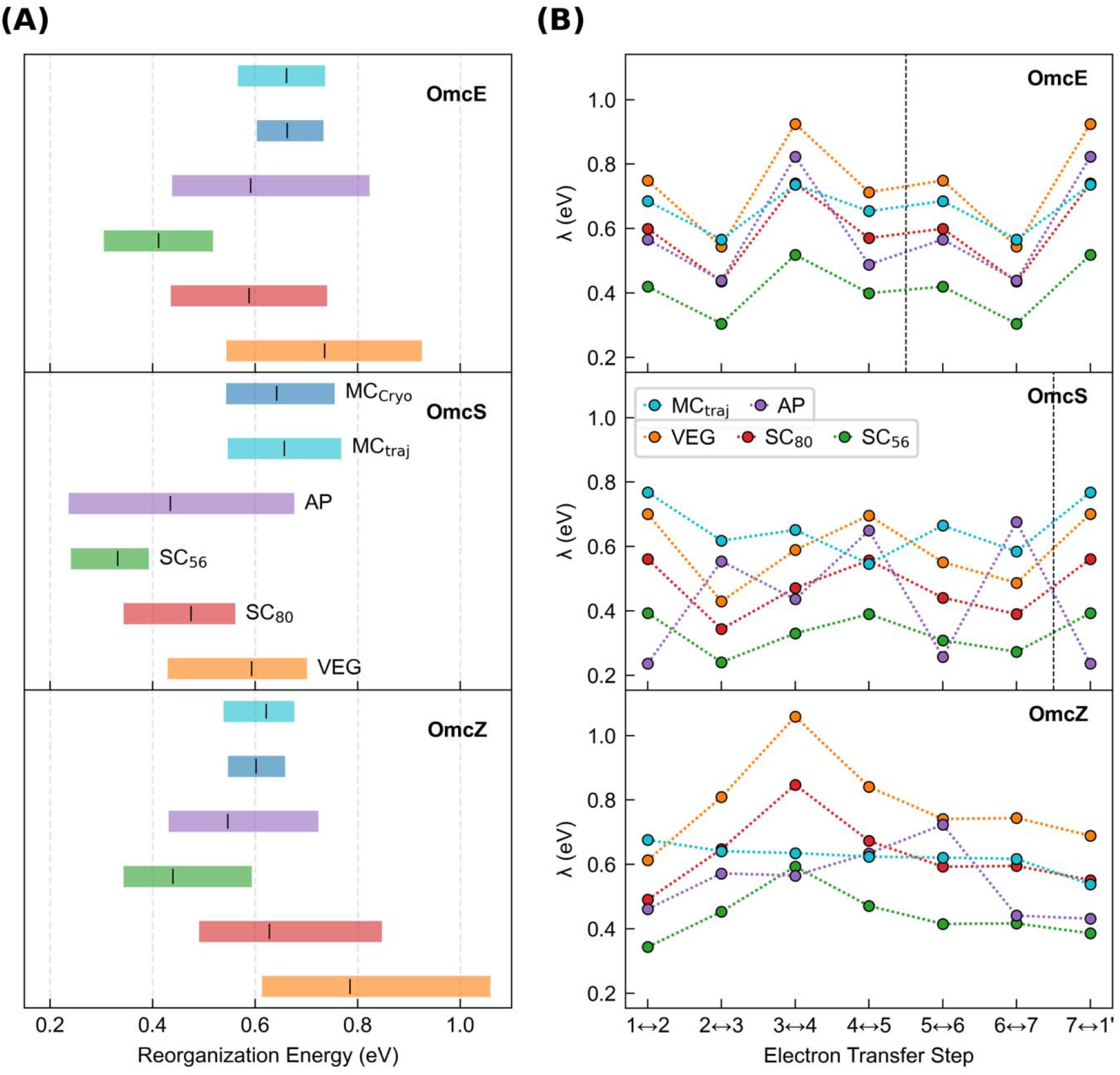
Outer-sphere reorganization energies in the cytochrome ‘nanowires’ is always tenths of an eV regardless of whether or how electronic polarizability is considered. **(A)** Range of for electron transfers through Omc-E, S, and Z according to different methods. **(B)** Variation in as a function of the position along the chain and the various computational methods. **(A–B)** MC_Cryo/traj_, VEG, SC_80_, SC_56_, and AP respectively indicate was estimated from an empirically parameterized form of the Marcus continuum equation applied to either the CryoEM geometries or molecular dynamics trajectories, Coulombic vertical energy gaps, those gaps scaled by factors of 0.80 or 0.56, or combined with the perturbatively-computed polarization energy for the active-site. Vertical das ed lines mark subunit boundaries to allow comparisons across filaments with fewer electron transfer steps per subunit than OmcZ. Data is repeated after the boundary because of the homopolymeric nature of the filaments.

The following discussion shows that all methods gave λ_out_ between 0.24 and 1.06 eV. Since these values are ≥20-fold larger than 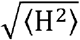 (Section 3.1), standard Marcus theory is the appropriate theoretical framework for describing electron transfer. The prior proposal of coherence-assisted transport predicated on unusually low (∼0.15 eV) reorganization energies in OmcS ^44^ can be put aside.

#### 3.3.1. Coulombic Vertical Energy Distribution Approach

λ_out_ was estimated from the distributions of vertical energy gaps (VEGs) in the reactant and product states for each heme-to-heme electron transfer in CNs previously submitted to MD simulations, namely, Omc-E, S, and Z. The distributions were reasonably Gaussian in every case (Supplementary Figures 1–3) and showed little sign of non-ergodic behavior: The ratio of the variance to Stokes shift reorganization energy was within ±0.2 of unity (Supplementary Table 3). λ_out_, taken as λ^rxn^ in Eq. 7, was (Figure 6, “VEG”) 0.544–0.925 (OmcE), 0.429–0.701 (OmcS), 0.613–1.06 (OmcZ) eV.

#### 3.3.2. Empirically Corrected Vertical Energy Gaps for Medium Polarizability

To account for electronic polarization of the environment in response to oxidation-state cycling at the active site, recommended scaling factors (SC) of 0.80 or 0.56 were applied to λ_out_ from the VEG approach.^109, 110^ The resulting ranges for Omc-E, S and Z ‘nanowires’ were 0.435–0.740, 0.343– 0.561, and 0.490–0.847 (Figure 6; SC_80_), or 0.305–0.518, 0.240–0.393, and 0.343–0.593 (Figure 6; SC_56_) eV.

#### 3.3.3. Perturbatively Corrected Vertical Energies for Active Site Polarizability

An alternative approach to correct for the polarizability of the heme active site was used in which the VEGs are expanded by perturbation theory to second order as a sum of Coulombic and polarization energies.^58, 107^ The polarization energies depend on the change of the polarizability tensor for the heme group between the oxidized and reduced states (Δ **α**) and the electric field (**E**) exerted on the heme by the protein-water environment.

##### 3.3.3.1. Heme Polarizability

Δ **α** was computed for His-His ligated hemes, both in vacuum and in the distinct protein-water binding sites of OmcS. Supplementary Tables 1 and 2 show that Δ **α** was minimally changed from its vacuum value by the protein-water environment and thermalization during MD, as well as replacement of the distal His ligand with Met. The analysis therefore focuses on the *in vacuo* results for the His-His ligated *c*-type heme found in all known CNs and makes comparisons to the literature on the His-Met ligated heme of cytochrome *c*.^61, 66^ The results are proposed to be general for both His-His and His-Met ligated *c*-type hemes in any hemoprotein.

DFT techniques have been found to accurately reproduce the (iso/aniso)-tropic polarizabilities of rigid to semi-rigid molecules,^111-113^ including a heme memetic.^114^ Using these methods and ensuring basis set convergence on the results, Figure 7 (Supplementary Table 1) shows that the isotropic polarizability (α_iso_) of the heme cofactor is 86 to 104 Å in either oxidation state regardless of the approximate density functional and basis set combination employed.

**Figure 7.**
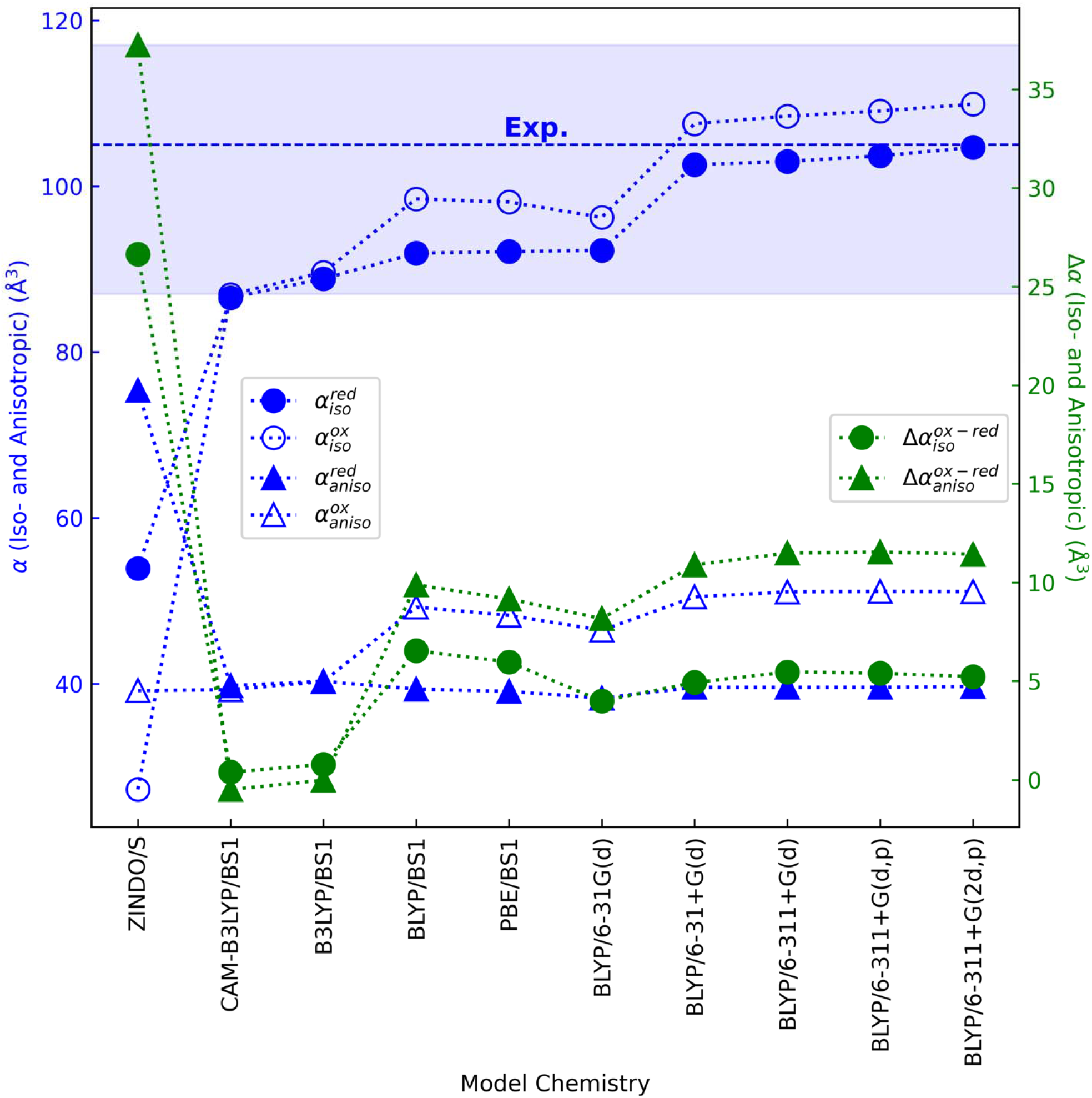
Validated and converged quantum chemistry models show that a bis-histidine ligated *c*-type heme is similarly polarizable in the reduced and oxidized states, far more so than previously predicted by the unvetted ZINDO/s method for this purpose.^58, 107^ The average (dashed line) and standard deviation (shaded region) of the experimental isotropic polarizability for the heme memetic tetraphenylporphyrin-iron(III) chloride are shown. ^114^ Neither the isotropic polarizability in the reduced state nor the anisotropic polarizability in either oxidation state was reported from the experimental work.

The range for α_iso_ fell within the experimental uncertainty for the heme memetic *meso*-tetraphenyl-iron(III) porphyrin,^114^ and exceeded the prior estimates of 54 and 27 Å^3^ in the reduced and oxidized states, respectively, from the Zerner’s Intermediate Neglect of Differential Overlap with Singles (ZINDO/S) method applied to a His-Met ligated *c*-type heme.^61, 66^

The change in isotropic polarizability between oxidation states (Δα_iso_) was found by DFT methods to be at most 6.5 Å^3^, which is also independent of environment and ligand set. The obtained Δα_iso_ was 4-fold smaller than that previously predicted by ZINDO/s, but it is in good agreement with the estimate obtained by analyzing the UV-vis spectra of cytochrome *c* in both redox states.^115^ Spectral analysis from 600 to 1.8 nm found no significant absorption intensity outside the visible range that would change this estimate.^116^

A similar story applies to the anisotropic polarizability (α_aniso_). DFT methods predict α_aniso_ to be ∼40 and ∼50 Å^3^ in the reduced and oxidized states, respectively, giving Δα_iso_ of ∼10 Å^3^. Prior ZINDO/s calculations on a His-Met ligated heme, by contrast, predicted 75 and 39 Å^3^ in the reduced and oxidized states, respectively, with a difference, again, four-fold larger than found by DFT methods.

Small changes in Δ**α** are consistent with small geometric changes (*e*.*g*., ∼0.01 Å and <1° in bond lengths and angles, respectively, at the BLYP/6-31G(d) level of theory) between the redox states and the low internal reorganization energy for the heme group. The discrepancy between ZINDO/s and DFT most likely reflects the inadequacy and unreliability of the former method for predicting polarizabilities. The use of ZINDO/s for this purpose requires summing contributions to the polarizability from vertical excitations computed up to 10 eV above the ground state.^61^ As the sum over excited states must grow to large (*e*.*g*., 100) numbers^61^ to converge the polarizability, increasingly larger errors accumulate when using a semi-empirical method parameterized to reproduce only the lowest-lying excited states. Furthermore, ZINDO/s, unlike DFT, has not been parameterized or benchmarked against experiment for computing polarizabilities.

The overall conclusion is that: Oxidation-state dependent changes in iso- and anisotropic polarizabilities for the heme cofactor are predicted by DFT to be far *smaller* (<15 Å^3^) than previously found with ZINDO/s semi-empirical theory, regardless of different axial ligands, molecular environments, or chosen DFT model chemistries.

##### 3.3.3.2. Protein-Water Electric Fields

The protein matrix for all structurally characterized CNs displays a formal net negative charge, where “formal” refers to the assumption of standard pK_a_s for titratable residues. On a per-subunit basis of the homopolymeric filaments, the formal net charges are -3*e* (A3MW92 and OmcS), -6*e* (OmcE), -8*e* (OmcZ), and -11*e* (F2KMU8).

Despite the almost quadrupling of per-subunit formal net charge, the heme-Fe centers in an interior subunit of all CNs experience a similar but wide range in electric field magnitudes from 0.07 to 0.7 V/Å (Supplementary Table 6). Across the binding sites in a given protein, the field magnitudes (|**E**|) are 0.34±0.13/0.37±0.10 (OmcZ), 0.36±0.20 (A3MW92), 0.42±0.22/0.36±0.22 (OmcS), 0.43±0.15 (OmcE), and 0.52±0.12 (F2KMU8), where the “/” indicates results for the two CryoEM structures of OmcS (PDB 6EF8 and 6NEF) and OmcZ (PDB 7LQ5 and 8D9M).

These field magnitudes were calculated (as done previously for heme systems^117^) directly on the CryoEM structures after geometry optimization, and therefore do not include conformational dynamics or solvation. For the three filaments simulated by MD in an explicit aqueous solvent with sufficient counterions for overall charge neutrality (Omc-E, S, and Z), the solvent typically suppressed |**E**| exerted by the protein matrix. Figure 8 shows the dependence of |**E**| in OmcZ on the cutoff distance for including the solvent; analogous figures for Omc-E and S are shown in Supplementary Figures S15–S16.

**Figure 8.**
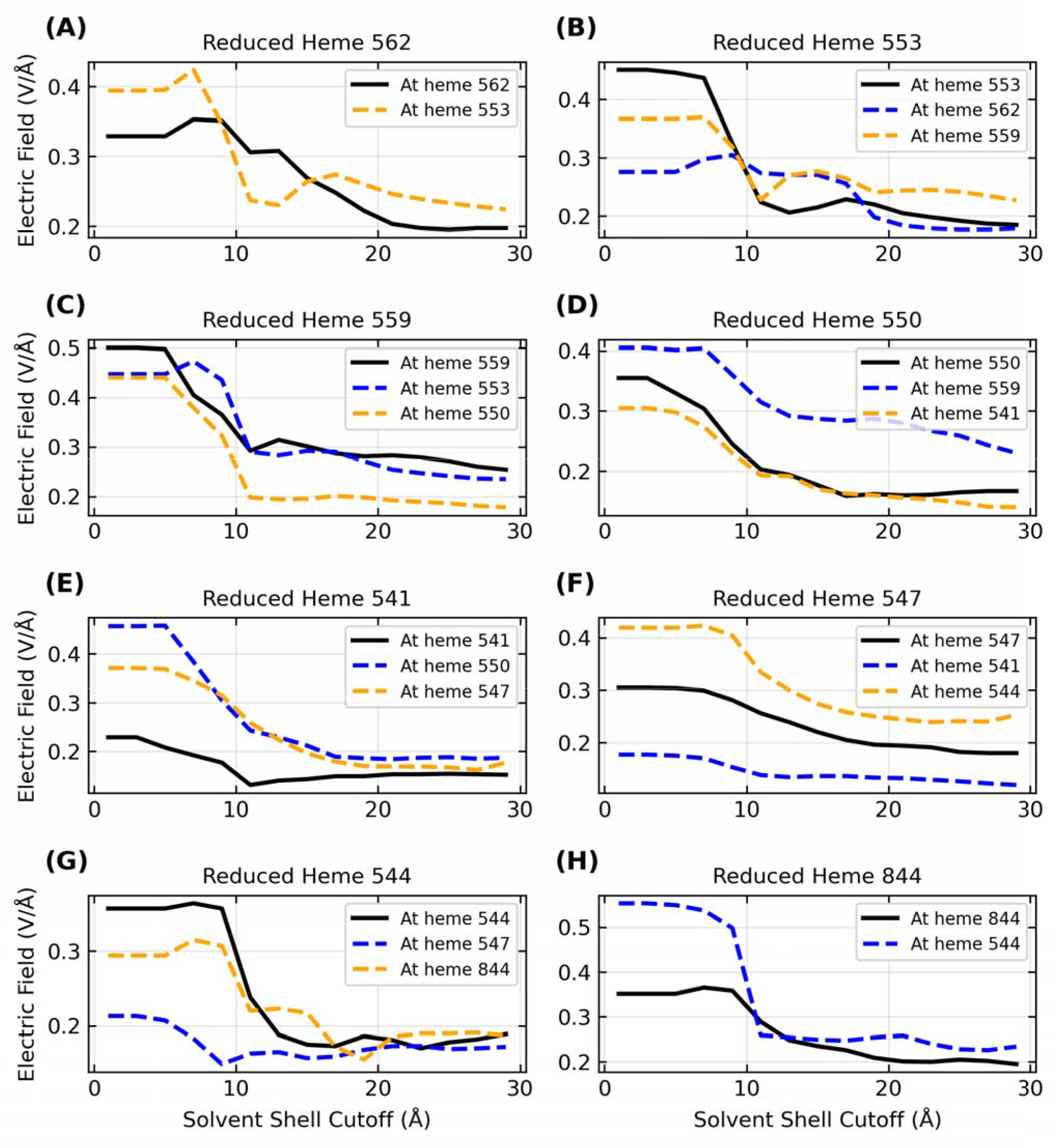
Inclusion of the explicit aqueous electrolyte with an increasing solvent cutoff causes a sizable dampening in the strength of the electric field exerted on the heme-Fe centers of OmcZ for the reduced donor (black curve) and the oxidized acceptor (blue/orange curve) heme on either side of it in the chain. The panels in alphabetical order track the movement of an electron through the filament where the sequence of electron donors by ID is 562 → 553 → 559 → 550 → 541 → 547 → 544 → 844.

The range of |**E**| shrank and shifted to lower values for each protein as more of the solvent was included. For the reduced state of each heme in OmcZ (black curves in Figure 8), for example, the range of |**E**| changed from 0.23–0.50 V/Å at a 1.0 Å cutoff to 0.15–0.25 V/Å at a 29 Å cutoff for the solvent (Supplementary Table 7). The conclusion is that the hydration state of CNs significantly affects the magnitude of the electric field experienced at the heme-Fe centers.

Since the solvent dampens the electric field exerted by the protein, and the total field experienced by the electron influences the activation barrier for the redox reaction, it follows that physiological redox conduction through a protein is sensitive to the hydration state. This fact is well known,^118^ and a prior study proposed that changes in the solvent microenvironment around OmcS contributes to its anti-Arrhenius conductivity behavior.^43^ In articulating the fact, though, it is clear that all electrical characterizations to date on air-dried CNs are biologically irrelevant, even apart from the known structural distortions that likely occur upon dehydration.^8^

But why, then, are reported conductivities the same for hydrated and dehydrated samples of CNs?^119^ The simplest hypothesis is that electrons are not conducted by a biological redox cascade in either state because of the chosen experimental technique. That is, the residence time of the electrons may be so short that there is insufficient time for the solvent to screen the changing electrostatic potential along the protein. Whether the solvent is absent or frozen on the timescale of electron transfer, it cannot participate in the process. If the solvent does not participate, the process is abiological electron transport, not biological electron transfer.^120^

In support of this conclusion, the same electrical measurements currently attributed to filamentous cytochromes^26, 119^ were argued to be incompatible with cytochromes before the CryoEM structures were solved.^121, 122^ Given that CryoEM has shown the filaments to be cytochromes,^26, 28, 83^ spectroelectrochemistry has confirmed their redox activity,^100^ and ground-state biological electron transfer is well-known to be redox-mediated, electrical measurements that have always been so inconsistent with redox chemistry that they required invoking a “new paradigm” of metallic-like conductivity^45, 46^ have simply never been biologically relevant.

Returning to the discussion of electric fields in the context of electronic polarizability and reorganization energy, Figure 9 shows a key result: The electric field in any of the CNs is similar but not identical in magnitude (as well as direction) at the heme-Fe centers of the donor (filled circles) and acceptor (unfilled circles) at each electron transfer step. Differences in the field along the heme chain can couple to the equal and oppositely signed Δ**α** for oxidation of the donor and reduction of the acceptor to produce net polarization energy and thereby change λ_out_.

**Figure 9.**
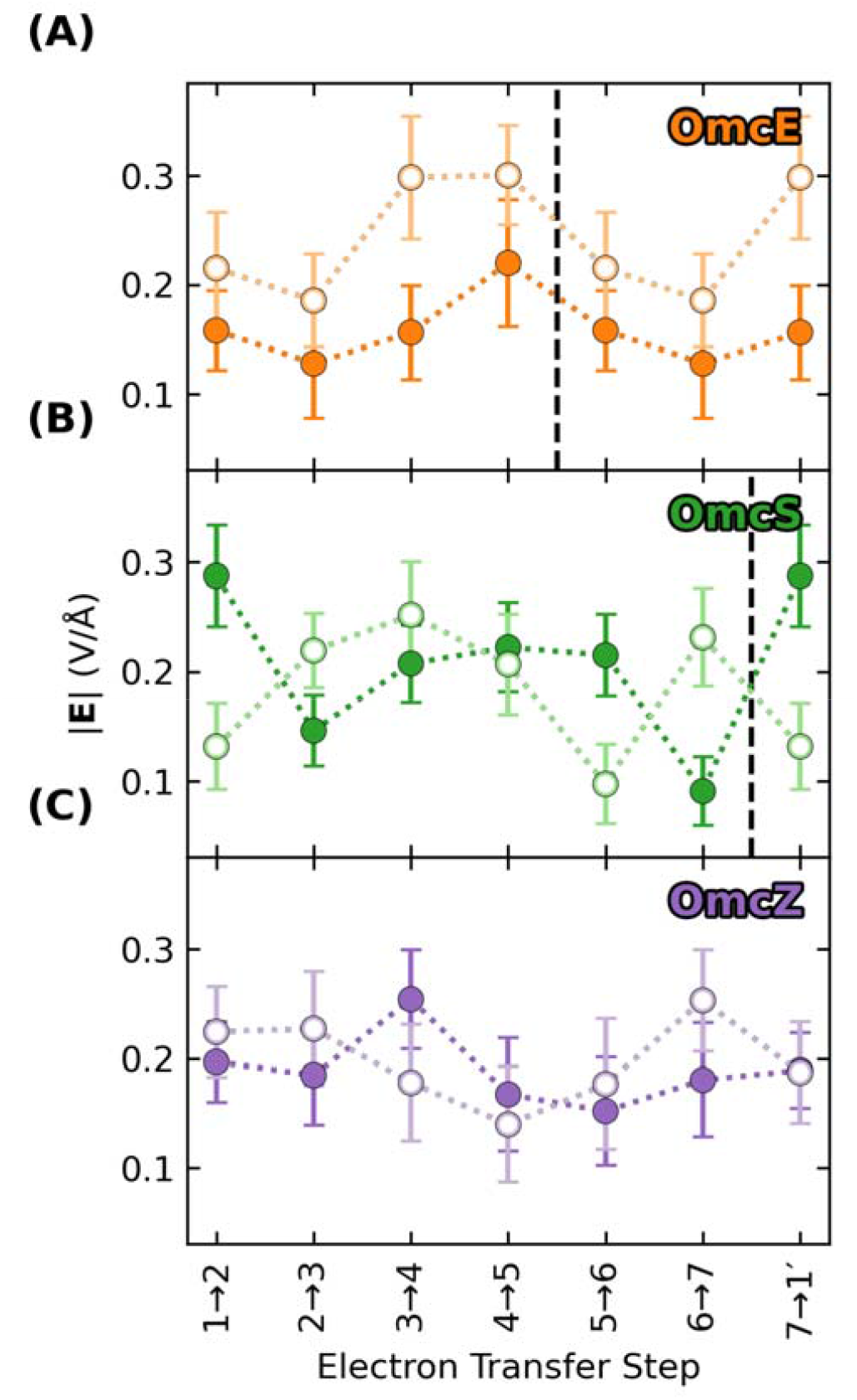
The electric field exerted on the donor (filled) and acceptor (unfilled) for a given electron transfer step is not identical, which means that the oppositely signed polarization energies of for the two redox partners will not perfectly cancel, and the net result will contribute to the reorganization energy. Vertical dashed lines mark subunit boundaries to allow comparisons across filaments with fewer electron transfer steps per subunit than OmcZ. Data is repeated after the boundary because of the homopolymeric nature of the filaments.

##### 3.3.3.3. Active Site Polarized λ_out_

Generally speaking, the polarization energy computed by Eq. 10 for Omc-E, S, and Z was negative (stabilizing) for the donor and positive (destabilizing) for the acceptor. Figure 10A shows the most common case: The donor was stabilized more than the acceptor was destabilized in the reactant state, and the effect was to decrease the magnitude of the total (Coulombic + donor polarization + acceptor polarization) VEG for the forward electron transfer. Conversely, the acceptor was destabilized more than the donor was stabilized in the product state, which decreased the magnitude of the total VEG for the reverse reaction. The result was a lowering of λ_out_. This active-site-polarization effect (shown in Figure 6 as “AP”) lowered λ_out_ by 0.11–0.23 (OmcE), 0.05–0.47 (OmcS), and 0.02–0.49 (OmcZ) eV (Supplementary Table 3).

**Figure 10.**
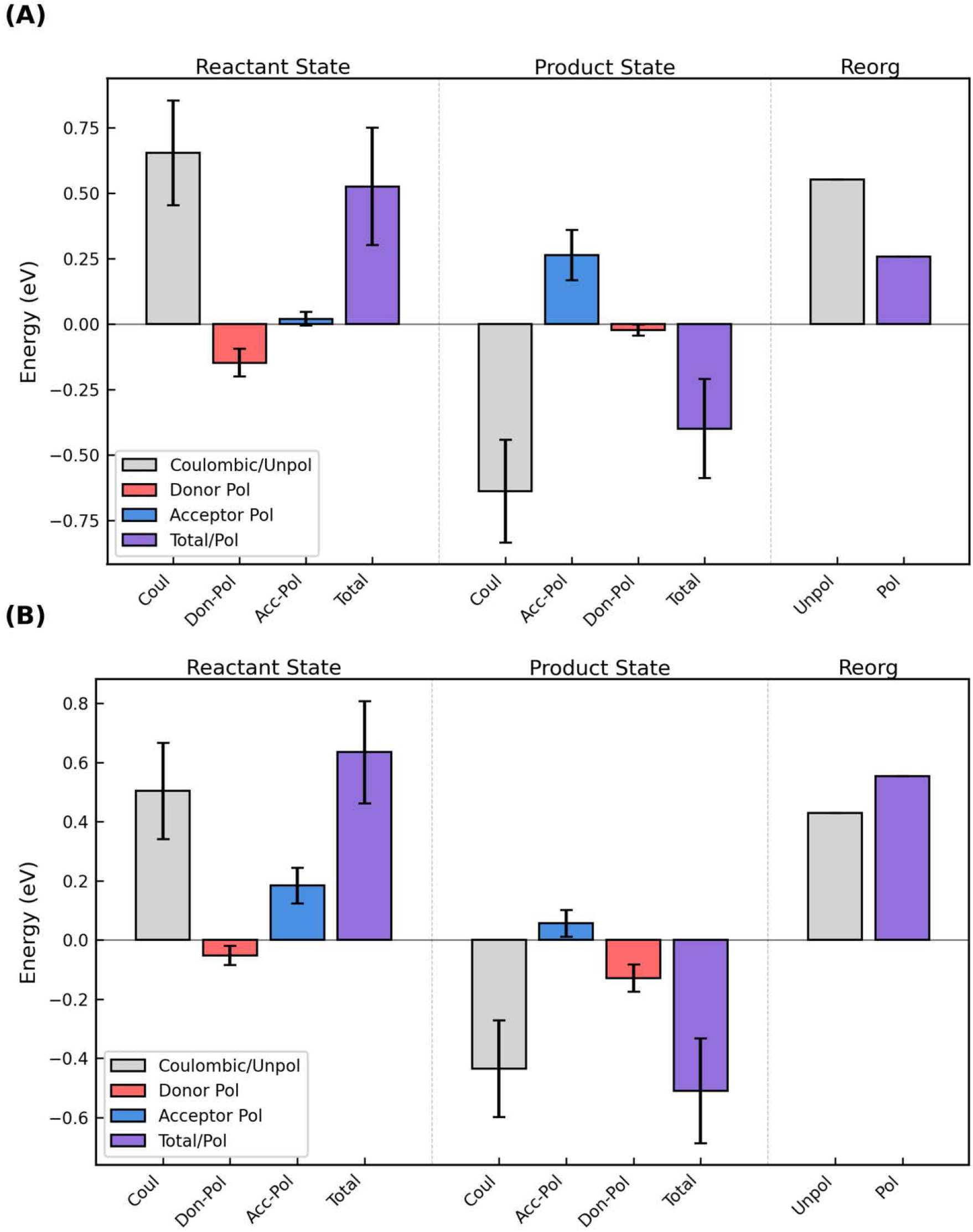
The oppositely signed polarization energies for oxidation of the donor and reduction of the acceptor in the reactant and product states can **(A)** decrease or **(B)** increase the total vertical energy gap and respectively cause a decrease/increase in the outer-sphere reorganization energy. The figure shows the Coulombic energy (gray), the separate polarization energies of the donor (red) and acceptor (blue), and the sum of those three terms (purple) for the reactant and product states, as well as the unpolarized (gray) and polarized (purple) reorganization energies for the electron transfer. The figure shows the results for **(A)** Heme #5 → #6 and **(B)** Heme #2 → #3 in OmcS; analogous figures for all electron transfer steps in Omc- E, S, and Z are shown as Supplementary Figures 17-33.

For two electron transfer steps, both in OmcS (Heme 2→3 and 6→1’, where the prime indicates the heme from the next subunit), the destabilization of the acceptor in the reactant state and the stabilization of the donor in the product state exceed, respectively, the stabilization of the donor in the reactant state and the destabilization of the acceptor in the product state (Figure 10B). The result was an increase in the VEGs for the forward and reverse reactions, and a larger λ_out_.

Figures analogous to Figure 10 are shown for each electron transfer step in Omc-E, S, and Z as Supplementary Figures 17–33.

λ_out_ by this AP approach was 0.438–0.823 (OmcE), 0.236–0.676 (OmcS), 0.431–0.723 (OmcZ) eV. Of note, the ergodicity factor changed with the inclusion of electronic polarizability from 0.9–1.2 for Omc-E, S, and Z to 1.3–1.5 (OmcE), 1.0–2.3 OmcS, and 1.2–1.7 (OmcZ) (Supplementary Table 3). The increased deviation from unity indicates a modest non-ergodicity that has been seen before, for example, for cytochrome *c*.^61^

##### 3.3.4. Empirically Parameterized Marcus Continuum Approach

A much simpler approach to estimate λ_out_ is to use a form of the Marcus continuum expression (“MC” approach) in which the static dielectric constant (ϵ_S_)was previously parameterized as a linear function of the total (donor+acceptor) SASA. ^98^ The approach is motivated by the facts that the solvent usually constitutes ∼50% of λ_out_, ^102^ and for hemoproteins, λ_out_ is controlled by SASA. ^123^

This approach (Eqs. 3 and 4 in Methods) only depends on structural parameters, which were measured from the CryoEM structures of the known CNs (Supplementary Figure 34A and Supplementary Table 4), as well as hundreds of nanoseconds of previously published MD simulations for three of these structures (Supplementary Figure 34B and Supplementary Table 5).

All adjacent heme pairs cluster around Fe-to-Fe distances of ∼9 and ∼11 Å, which are respectively characteristic of slip- and T-stacked packing geometries. The donor+acceptor SASA of adjacent heme macrocycles (not including propionic acid substituents) was typically <78 Å^2^. ϵ_S_ of the heme binding sites ranged from 5.2–7.6, or in more chemically intuitive terms, from being slightly more polar than chloroform (ϵ_S_ = 4.8 at 25°C) to as polar as tetrahydrofuran (ϵ_S_ = 7.6 at 25°C) solution.

The similar Fe-to-Fe spacings and macrocycle SASAs constrained λ_out_ in all CNs to the narrow range 0.543 to 0.770 eV (Figure 6, “MC”). Thermal averaging caused relatively minor changes in Fe-to-Fe distances, donor+acceptor SASA, and λ_out_.

#### 3.3.5. Summary of the Effect of Polarizability on λ_**out**_

λ_out_ decreased by 0.105–0.225 eV (11–31%) for OmcE using the AP versus VEG method. For OmcZ, λ_out_ was reduced by 0.018–0.494 eV (2–46%). OmcS gave a more complicated picture: λ_out_ decreased by 0.047–0.463 eV (7–67%), except for two electron transfers where it increased 0.125– 189 eV (29–39%). The conclusion is that the recommended scaling factors of 0.56–0.80 are generally in the appropriate range, but their universality does not hold.

For its simplicity and sufficient accuracy, the MC approach was mostly used for the comparative study of the structurally characterized CNs in the rest of this work. The uncertainty in activation energies and rates introduced by this choice is discussed below.

### 3.4 Intra-Filament Electron Transfer Activation Energy

The activation energy (E_a_) in standard Marcus theory is defined as (Eq. 14):

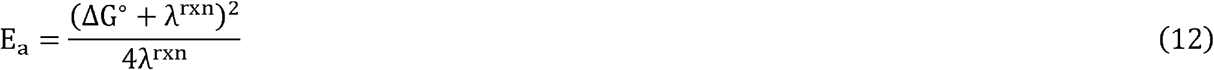

Using ΔG°s for the geometric oxidation sequence imposed by the linear topology of the CNs, E_a_ is *always* ≤0.3 eV (Figure 11A; Supplementary Table 8). This result is insensitive to the method used to compute (Figure 11B; Supplementary Table 9). Relative to the MC approach for λ_out_, the absolute average deviation in E_a_ caused by the other methods are 0.027 (SC_80_), 0.033 (VEG), 0.037 (AP) and 0.058 (SC_56_). Differences of this magnitude would change the computed rates in the next section by a factor of 10, but the conclusions would be unaffected.

**Figure 11.**
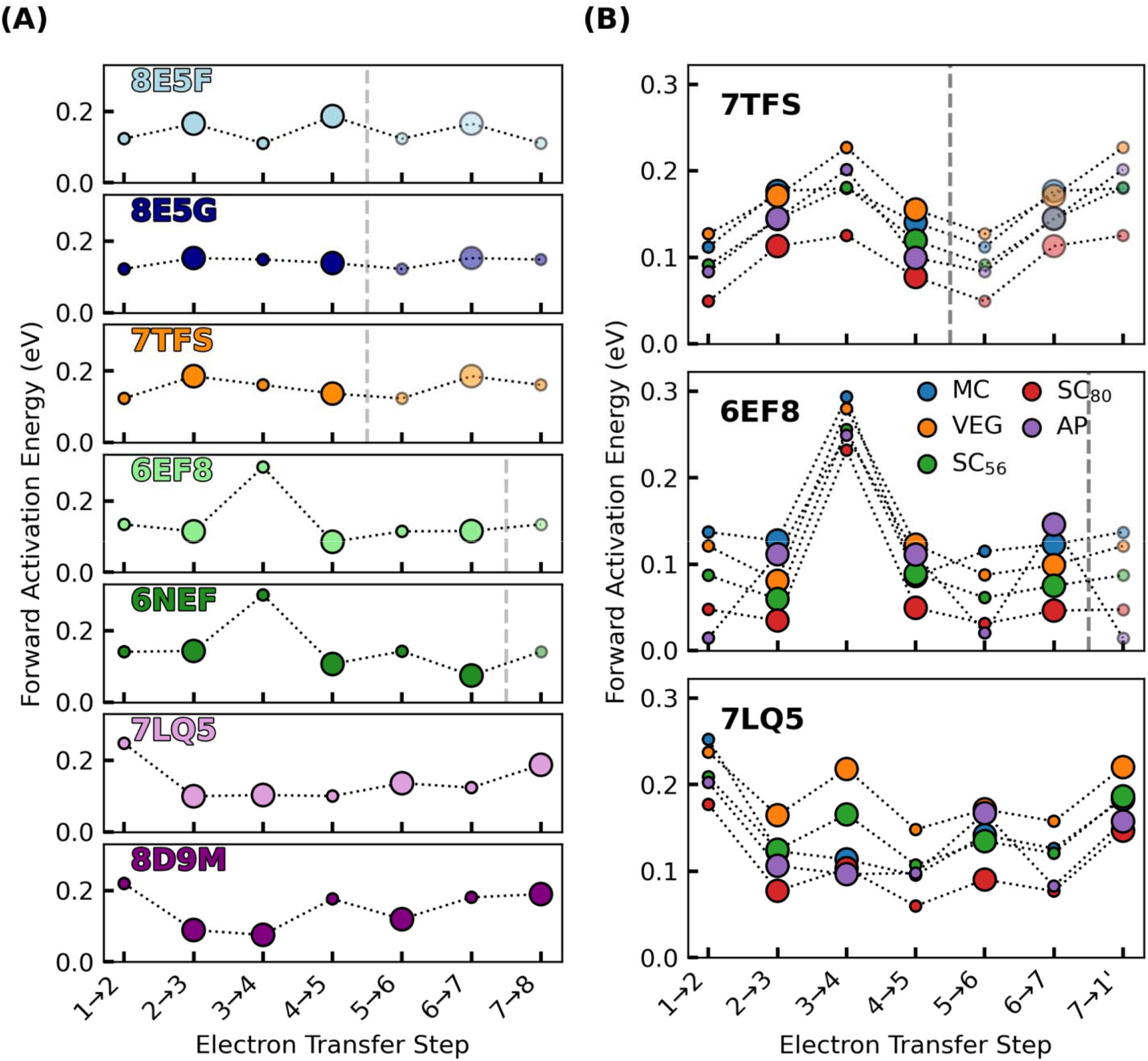
Activation energy for electron transfer in the known cytochrome ‘nanowires’ is always ≤0.3 eV, regardless of if or how electronic polarization is included in the estimate of outer-sphere reorganization energy. Panel **(A)** shows the cytochrome ‘nanowires’ by PDB accession code: 8E5F, 8E5G, 7TFS, 6EF8/6NEF, 7LQ5/8DoM = A3MW92, F2KMU8, OmcE, OmcS, and OmcZ. Panel **(B)** shows the variation in activation energy for three of these filaments for which the outer-sphere reorganization energy was computed by various methods discussed in the main text. The size of the markers in both panels is proportional to the electronic coupling. Activation energies for the forward reactions are shown; Supplementary Figure 35 shows the activation energies for the backward reactions. All data is presented in Supplementary Tables 8 and 9.

**Figure 12.**
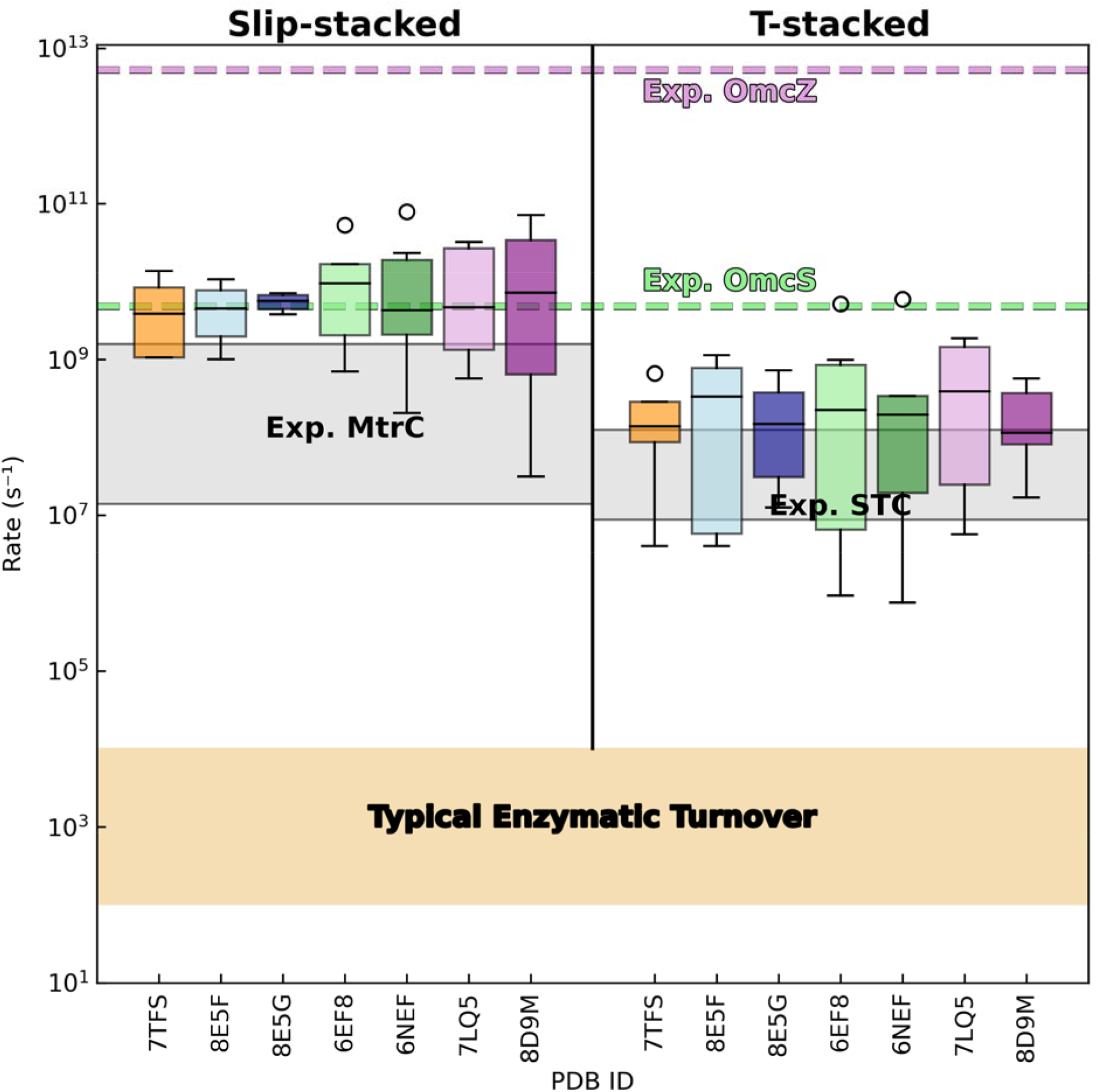
Electron transfer rates in cytochrome ‘nanowires’ are not metabolically limiting, comparable to rates in other multihemes, and orders of magnitude slower than implied by the reported conductivities. Distributions of predicted (forward and backward) rates between slip- and T-stacked heme pairs are shown for the known cytochrome ‘nanowires.’ The gray shaded regions show experimental rates reported for photosensitized variants of the metal reducing cytochrome (MtrC)^85^ and small tetraheme cytochrome (STC)^86^ from *Shewanella oneidensis*. The beige-shaded region covers the range of typical rates for enzymatic turnover.^138^ The dashed green and purple lines indicate the electron transfer rate that would be needed for every intra-wire heme-to-heme electron transfer to reproduce the previously reported conductivities of single Omc-S and Z filaments; see the main text for the derivation of these effective hopping rates.

With the MC approach for λ_out_, E_a_ is 0.110–0.255 for A3MW92, F2KMU8, and OmcE, 0.075–0.300 eV for OmcS, and 0.076–0.248 eV for OmcZ. Thus, it appears that the range for E_a_ is conserved among CNs.

### 3.5. Intra-Filament Electron Transfer Rates

Given the energetic parameters discussed in the preceding sections, the computed electron transfer rates within slip- and T-stacked heme pairs across all CNs cover (as expected)^47^ large and overlapping ranges (Figure 11; Supplementary Table 8): 3.1 ×10^7^ − 7.9 × 10^10^ s^-1^, and 7.5 ×10^5^ − 5.9 × 10^9^ s^-1^, respectively. However, the corresponding median rates across CNs reside in much narrower ranges of 3.8 ×10^9^ − 9.5 × 10^9^ and 3.4 ×10^8^ − 3.9 × 10^108^ s^-1^. This result suggests: (1) Typical rates within T-stacked pairs are at least an order of magnitude slower than within slip-stacked pairs; (2) The rates for specific heme packing geometries are roughly transferrable between proteins; and (3) The various CNs support similar electron fluxes (see Section 3.6). The following subsections put the computed rates into context with experimental expectations (Section 3.5.1) and the reported conductivities of the CNs (Section 3.5.2).

#### 3.5.1. Comparison to Experimental Expectations

The minimum, maximum, and median electron transfer rates predicted for the T-stacked geometry in all CNs are typically within a factor of ∼10 of the corresponding experimental rates for the same heme packing geometry in the small tetraheme cytochrome (STC).^86^ Predicted minimum, maximum, and median rates for slip-stacked hemes are typically within a factor of 60 (but in one case as high as 272) compared to rates measured in the metal reducing cytochrome (MtrC).^85^ As observed for the T-versus slip-stacked heme pairs in STC and MtrC, electron transfer rates were 10-fold smaller in the former versus latter heme packing geometry for the CNs.

The one-to-two order of magnitude agreement with experimental expectations should be weighed with the following considerations:

(1) The actual electron transfer rates in CNs are unknown. The comparison *assumes* that the rates are transferrable from different proteins for a given heme packing geometry. The similarity in the distributions of computed rates for different CNs in Figure 11 makes this assumption seem reasonable.

(2) The overestimation of slip-stacked rates relative to experimental expectations is inconsequential in the context of multi-step electron transfer: The slowest step in a multi-step reaction is rate-limiting, and ∼50% of the heme pairs in CNs are of the rate-throttling T-stacked variety. For this packing geometry, the predicted rates are in much closer agreement with experimental expectations.

(3) An order-of-magnitude uncertainty is introduced by the different methods for computing λ_out_.

(4) A ≤14-fold discrepancy between theory and experiment was reported for an analysis on the same protein using state-of-the-art MD and DFT methods.^86^ A far more approximate but expedient protocol was implemented in the BioDC program and used to facilitate the comprehensive study of all structurally-characterized CNs.

Inspired by the work of Blumberger and co-workers,^53^ the BioDC program^124^ respectively estimates the reaction free energy, reorganization free energy, and RMS coupling from classical (PBSA) electrostatic calculations (Section 3.2), the empirically parameterized form of the Marcus continuum equation (Section 3.3.4), and the tilt angle between adjacent hemes (Section 3.1). These methods were applied only to the CryoEM geometries, although the independently solved structures for both Omc-S and Z gave similar results throughout this work.

Given only one or two geometries for each protein, the approximate methods used to compute the energetics of electron transfer, the exponential dependence of the rate on these energetics, the accepted order-of-magnitude error using a much more sophisticated protocol, and the fact that the actual rates in CNs are unknown, the obtained level of agreement with experimental expectations seems reasonable. It is also sufficient for the conclusions of this work.

#### 3.5.2. Comparison to Reported Electrical Conductivities

As already noted, electron transfer rates in CNs have not been measured yet. However, an *effective* rate can be derived from the reported conductances(G)^47,119^ via the Einstein-Smoluchowski relationship and standard electrical definitions (Eq 13).

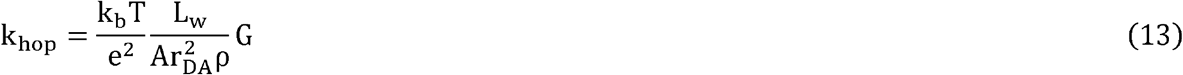

The first term collects the Boltzmann constant (k_b_), temperature (T), and elementary charge (e). The second term comprises physical properties of the CN; namely, the length (L_w_) and cross-sectional area (A) of the conductor, the average charge-hopping distance (r_DA_), and the charge density of the filament (ρ).

L_w_ is taken as 3×10^−5^cm because this is a typical electrode spacing used in characterization studies. ^47^ The heme chain at the core of each CN is approximately cylindrical, and the radius of the heme group is 7.5 ×10^−8^ cm;^44^ A is therefore π*r*_2_ =1.8 ×10^−14^ cm. The minimum edge-to-edge separation is the appropriate distance metric for heme-to-heme electron transfer, ^125^ and for each CN r_DA_ ≈ 5 Å. More specifically, for the only two CNs (Omc-S and Z) electrically characterized so far,^47, 119^ r_DA_ is 4.86 (PDB 6EF8) – 5.01 (PDB 6NEF) ×10^−8^ cm for OmcS, and 4.40 (PDB 7LQ5) – 4.59 (PDB 8D9M) ×10^−8^ cm for OmcZ. The shortest average distance for each protein is used below.

ρ is given by the number of reducing equivalence per unit volume. Each heme can only be occupied by one reducing equivalent at a time, and the maximal conductivity is realized when 50% of the hemes are reduced, because in that state, there is an equal proportion of charge donors and acceptors. In the fully reduced (oxidized) state, there would only be electron donors (acceptors), and no electron transfer would be possible. This physical picture is one reason why the reported observation of 6-fold higher conductivity in the fully reduced versus fully oxidized state of OmcS^47^ does not make physical sense for biological, thermally activated, multi-step redox conduction, and thereby suggests these electrical characterizations are biologically irrelevant.

If ρ is taken for maximal conductivity, there are 3 or 4 reducing equivalence in a subunit of Omc-S and Z, respectively. The length of a subunit in these homopolymers is 4×10^−7^ and 5×10^−7^ cm, respectively, and A is the same because both have a conductive heme chain. Thus, ρ is 3.6–3.9 × 10^−20^ charges/cm^3^.

A similar charge density was reported for filaments from *Geobacter sulfurreducens* when they were argued to be composed of pili instead of cytochromes.^126^ It has been argued that this data should be re-interpreted *as if* it was measured on the now-known cytochrome OmcS,^26^ but the comparison is still flawed: The charge density was measured by subjecting the protein to a 10-V bias, and thereby accessed electronic states completely forbidden to biology.

With the discussed physical parameters, the reported conductances of 1.1 ×10^−10^ and 4.9 ×10^−8^ S (Siemens) for Omc-S and Z respectively give effective hopping rates by Eq. 13 of 3.3 ×10^−10^ and 1.4 ×10^−13^ s^-1^. These rates are physically unreasonable for the multiheme architecture of CNs as described in the following paragraphs.

The effective k_hop_ computed by Eq. 13 assumes that uncorrelated electron transfers occur between sites of localization along a uniform chain. Operation of linear electron transfer chains, like those found in CNs are well described without site correlations, even in the presence of site-site interactions.^127^ But CNs are not uniform since every-other step is ∼10-fold slower than the preceding step. The effective rate for diffusion over alternatingly slow (T-stacked) and fast (slip-stacked) heme-to-heme rates cannot exceed the slowest rate. And yet, the k_hop_ s implied by the experimental conductivities are 10^2^ and 10^5^-fold larger than the fastest known rate (1.25 ×10^−8^ s ^−1^) for the rate-limiting T-stacked packing geometry.^86^

The effective k_hop_ for OmcZ is even more implausible since it is at the coupling-maximized ‘speed limit’ of 10^13^ s^-1^ proposed by Gray and Winkler^128^ for a van der Waals metal-to-metal contact distance of 3 Å. Electronic couplings in OmcZ are weak (Section 3.1) and the metal-to-metal distance is 9–11 Å.^28, 83^

These results complement the conclusion in Section 3.3.3.2 that the electrical measurements are biologically irrelevant. Some of the excessive rates may be due to the abiological nature of the electrical contacts with the protein,^8, 94^ Unlike molecular redox partners in biology, the electrodes are not of molecular dimensions, and the observed current scales with the number of contacts. For example, the atomic force microscopy (AFM) tip used in characterization studies^47, 119^ spanned ∼10 subunits of the filaments.

It has also been noted that drop-casting, air-drying, and crushing proteins under an AFM tip on a denaturing bare gold electrode has little relationship with biological molecular recognition and redox chemistry.^8, 51^ The electrical measurements, by virtue of the experimental techniques, are biologically irrelevant.

A key insight of the present work is to put aside these abiological measurements to instead ask if the computed conductivities of CNs suffice for their proposed role in cellular respiration.

### 3.6 Intra-Filament Electron Flux

CNs are proposed to discharge the metabolic flux of electrons from a microorganism. Logical questions are then: What is the metabolic flux and do the conductivities of the CNs permit a bioenergetically reasonable number of them to serve this function? An upper bound for the single-filament conductivity needed for cellular respiration is derived in Sections 3.6.1–3.6.2, and the computed conductivities are compared to it in Sections 3.6.3–3.6.4. The metabolic discussion focuses on *Geobacter sulfurreducens* since this model organism produces 3/5 of the known CNs.

#### 3.6.1 Metabolic Requirements versus Solid State Conductivities

A *G. sulfurreducens* cell respires at a rate of ∼10^6^ e^-^/s regardless of whether the terminal electron acceptor is an electrode,^1, 7^ a metal salt,^2, 6^ or a mineral.^3^ Independent of the identity of the cation to be reduced (Ag^+^, Fe^3+^, Co^3+^, V^5+^, Cr^6+^, and Mn^7+^), the different redox potentials of the metal salts (∼0.21 V for AgCl versus ∼0.8 V for AgNO_3_ on the normal hydrogen electrode scale),^6^ or the downregulation/deletion of some cytochromes in the microbe, the rate of Fe^2+^-heme oxidation by the metal salts always equaled the rate of Fe^3+^-heme reduction by the respiratory machinery; what changed were the proportions of Fe^2+/3+^-hemes in the microbe. This result suggested that the electron efflux reflected the intrinsic respiratory rate that the cell must maintain for ATP synthesis. A similar rate of ∼10^5^ e^-^/s/cell was found for the bacterium *Shewanella oneidensis*.^4, 5^

Note that the respiratory rate is independent of whether CNs were specifically expressed in the experiments. How electrons exit the cell is a different question from how many electrons need to exit the cell.

It is not true—in fact, just the opposite—that cellular respiration is more efficient if/when CNs are involved. When a microorganism must switch from using an intra-to extracellular electron acceptor, protons from the oxidation of organic matter are left behind to accumulate in the cytoplasm and reduce the proton motive force for ATP synthesis.^93, 129^ Respiration, in fact, would entirely shutdown if there was no mechanism to remove some of the protons.^130^ Also, no energy is realized for the microbe through EET:^93^ The electrons are de-energized ‘spent-fuel’ at the end of the electron transport chain in the inner membrane that simply needs to be discarded to make way for respiration to continue. Whether or not CNs help to discharge respiratory electrons when intra-cellular acceptors are not available, they do *not* power life.

*G. sulfurreducens* realizes the metabolic flux of ∼10^6^ e^-^/s at a low potential difference between the intra- and extracellular redox half-reactions. Voltametric experiments indicated that *G. sulfurreducens* reaches its maximum respiratory rate on anodes posed at -0.1 V versus SHE and does not take advantage of the additional potential energy available at higher potentials.^131^

At an applied bias of -0.1 V vs. SHE, an electron flux of 6.2 ×10^−6^ s ^−1^ e^-^/s, or 1.0 pA gives, by Ohm’s law a conductance of 9.9 ×10^−12^ S. This value is the total conductance the ‘circuits’ must have that connect the cell to extracellular electron acceptors. If a *single* filament with the dimensions and charge density of Omc- S or Z (Section 3.5.2) were to discharge the *entire* metabolic current, its conductivity would need to be 1.7 ×10^−2^ S/cm. This is certainly an overestimate because (1) most studies indicate *G. sulfurreducens* generates ∼10-fold less current than the assumed 1 pA,^1, 2, 6, 7^ and (2) *G. sulfurreducens* has alternate routes (e.g., porin-cytochrome complexes) for expelling electrons.^132^ Only a fraction of the metabolic current would need to be discharged by a CN. And yet, as a single-filament conductivity, it is 10^2^–10^5^-fold *smaller* than the conductivities reported for single Omc-S and Z filaments. Why would a bacterium ever need such highly conductive filaments?

A natural thought may be for communal growth. However, biofilms on electrodes serving as infinite electron sinks are likely laboratory artifacts with no relevance to the physiology of *G. sulfurreducens* in its normal habitat. If a natural electron sink, Fe(III) oxyhydroxide, occupies 50% of the space around a *G. sulfurreducens* cell, bioenergetic calculations^133^ showed that the cell would need to reduce all available Fe(III) within a radius of 2–4 μm just to *approach* generating enough ATP for single-cell doubling. There is simply not enough oxidized mineral in a geographical area for *G. sulfurreducens* to support a multi-layer, microns-thick biofilm.

Living as a solitary microbe on a grain of sand,^133^ a *G. sulfurreducens* cell would have no use for the conductivities reported for its filaments. There are physical limits to the rate of intracellular acetate oxidation. For a single cell to make use of the conductivities reported for Omc-S and Z, the cell would respectively need to consume 9 ×10^−8^ and 4 ×10^−9^ molecules of acetate each second. This calculation is based on the fact that each acetate molecule yields 8 electrons, and ∼95% of those are used for ATP synthesis.^134^ Meanwhile, the known respiratory rate of ∼10^6^ e^-^/s only requires ∼10^5^ molecules of acetate per second.

Since the reported conductivities vastly exceed the metabolic needs of a single *G. sulfurreducens* cell, and the cell primarily lives in a solitary state in nature, we take the estimate of the needed conductivity from the respiration rate and consider the solid-state measurements to be biologically irrelevant.

#### 3.6.2. An Estimate of Maximal Single-Filament Conductivity for Cellular Respiration

It is not known how many filaments a *G. sulfurreducens* cell expresses, but it is certainly more than one. A study observed >20 filaments thought to be pili^135^ but now argued to be OmcS.^26^ A bioenergetic analysis previously suggested that 100 filaments/cell would be reasonable.^133^ If 100 filaments are parallel, carry the entire metabolic flux of electrons, and have the same physical of properties of Omc-S or Z, the filaments would need on average a charge diffusion constant by Eq. 14 7.5 ×10^−8^ cm^2^/s.

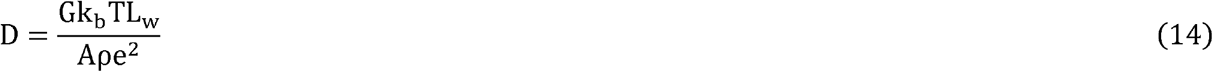

Can this diffusion constant be achieved within the above-mentioned energetic bounds for 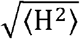(T-stacked: 0.002–0.006 eV; slip-stacked: 0.007–0.010 eV), ΔG° (±0.3 eV), and λ^rxn^ (0.24–1.06 eV)?

Monte Carlo simulations answered this question in the affirmative. Sets of 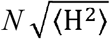, *N*ΔG° s, and *N* λ^rxn^ s were generated, where *N* is the number of electron transfer steps. Parameter sets were accepted if the associated diffusion constant was within 10% of the target value. For a CN with the TSTSTS unit cell of OmcS or the TSSTSTS unit cell of OmcZ (excluding the heme that branches from the main conductive chain), 1.0 ×10^6^ unique sets were found after 6.4–7.5 ×10^7^ sets were attempted, respectively.

Figure 13 summarizes the statistics for the compatible ΔG°s, λ^rxn^ s, and resulting E_a_ s from the Monte Carlo simulations, as well as the parameter sets that had the largest and smallest diffusion constants within 10% of the target value. Movies of representative parameter sets are included in the Supplementary Material.

**Figure 13.**
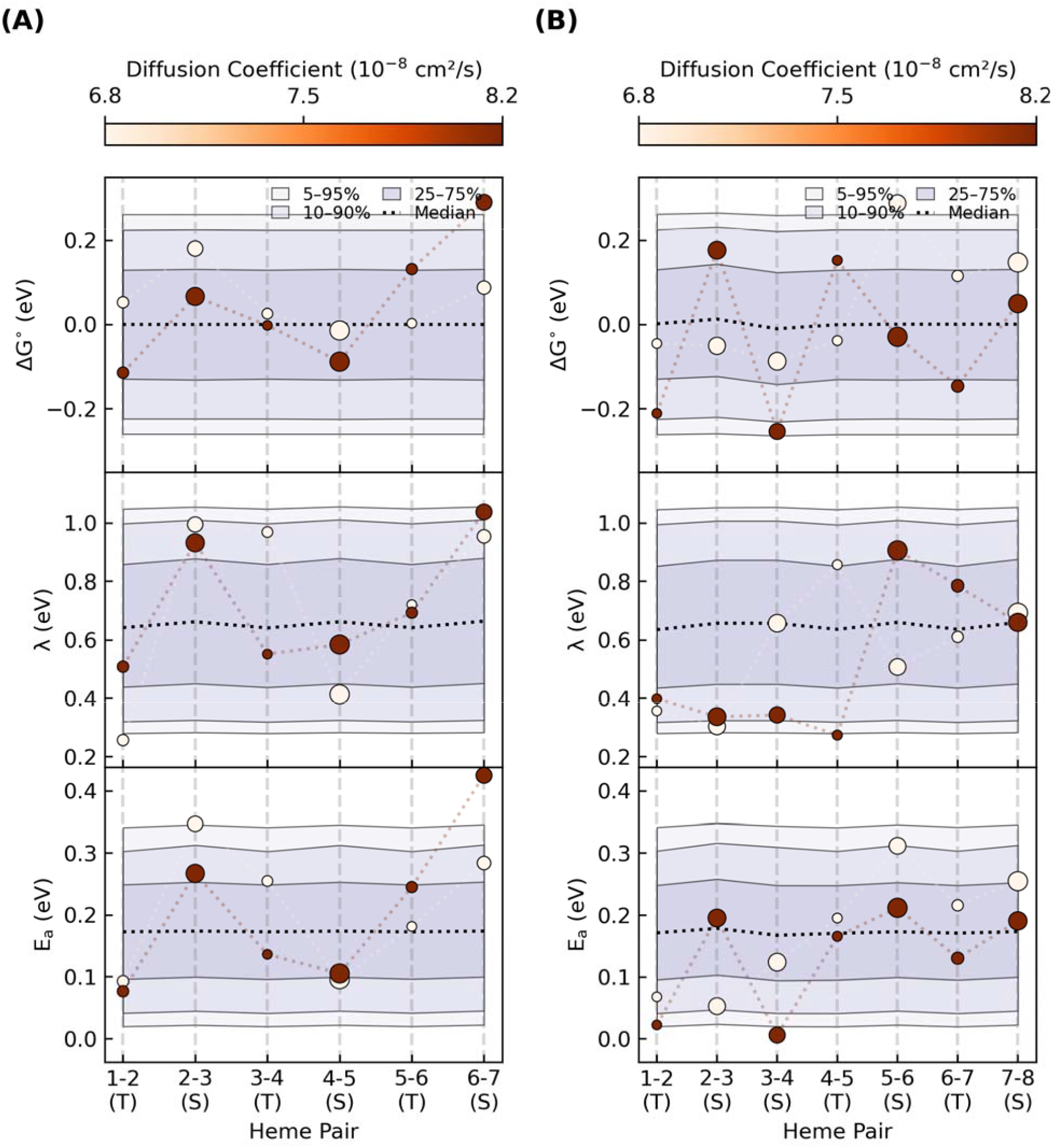
An enormous combination of energetic parameters can yield the maximum single-filament charge diffusion constant needed for cellular respiration. Each panel summarize the percentile ranges for one-million parameter sets that gave diffusion constants within 10% of cm^2^/s. The markers indicate the sets that gave the highest and lowest diffusion constants, with the size of the markers proportional to the electronic coupling for that step. Panels **(A)** and **(B)** show the repeat patterns for heme packing in Omc-S and Z, respectively.

#### 3.6.3. Cytochrome ‘Nanowires’ Suffice for Cellular Respiration

To make comparisons with the metabolically-required maximal diffusion constant for expelling electrons in the previous section, the set of k_et_s for moving an electron through a unit cell of each homopolymeric CN was used to evaluate the analytical Derrida formula^63^ for diffusive charge hopping along the periodic (pseudo)-one-dimensional heme chains The obtained diffusion constants and the predicted number of filaments/cell given those values are shown in Figure 14.

**Figure 14.**
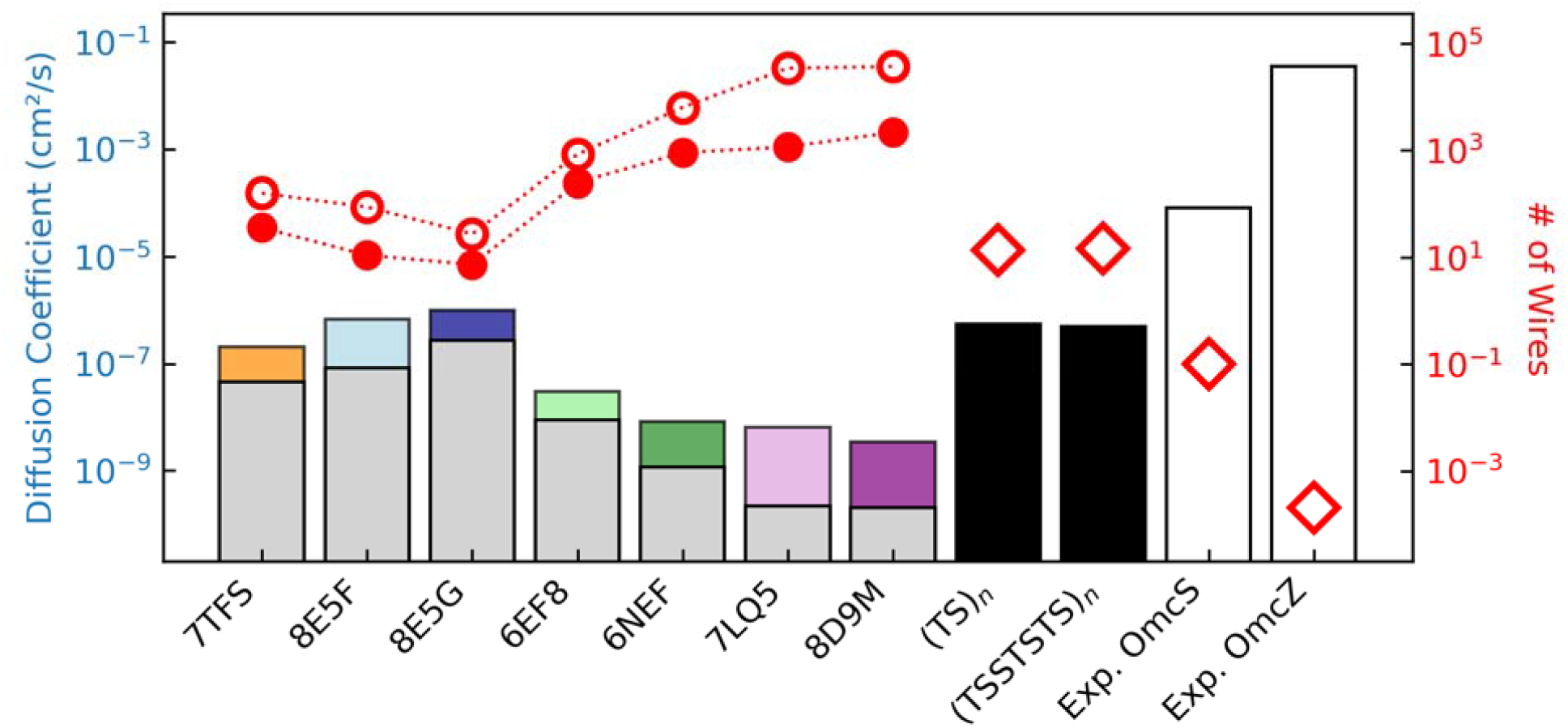
Charge diffusion within structurally characterized cytochrome ‘nanowires’ (colored ba s) is sub-optimal compared to heme chains with the same packing geometries but the fastest known electron transfer rates at every step (black bars), which in turn are unphysically dwarfed by the reported conductivities for Omc-S and Z (white bars). The colored versus gray bars show the influence of heme-heme interactions on the free energy landscapes. The red filled/unfilled circles show the number of filaments/cell needed given the computed diffusion constants with/without considering redox anti-cooperativities, and the estimate of the total (cellular) diffusion constant of cm^2^/s needed for parallel filaments. The unfilled diamonds indicate the number of filaments/cell given the diffusion constants obtained for the idealized heme chains or from the reported conductivities.

Diffusion constants (colored bars) range from 0.8 –2.7 ×10^−7^cm^2^/s for the tetrahemes A3MW92, F2KMU8, and OmcE; 1.1 – 8.9 ×10^−8^ cm^2^/s for OmcS depending on the CryoEM structure; and 2.0 – 2.2 ×10^−10^ cm^2^/s for OmcZ depending on the CryoEM structure. These diffusion constants are typically no more than a factor of 8 larger than when redox cooperativities are neglected (gray bars) in the geometrically imposed linear sequence of heme oxidations. The exception is OmcZ for which the increase is 17-(PDB 8D9M) or 30-fold (PDB 7LQ5). Differences between the colored and gray bars reflect the influence of redox anti-cooperativities on the ΔG° landscapes, as discussed in Section 3.2.3.

If the entire respiratory flux of electrons was discharged through parallel CNs having these diffusion constants, tens to hundreds of the tetra- or hexahemes, or thousands of the octaheme (red markers in Figure 14; filled/unfilled circles = with/without redox anti-cooperativities) would be needed. Since the cell may discharge less current than assumed here, the filaments would only need to discharge a fraction of it, and tens to hundreds of filaments are expected to be expressed, the prediction of the *maximal* number of filaments seems reasonable.

#### 3.6.4. Cytochrome ‘Nanowires’ are not Optimized for Electrical Conductivity

That the conductivity of CNs is biologically sufficient does not mean it is optimal. We already saw that there are physical constraints on the energetics of electron transfer by virtue of the multiheme architecture. Within those constraints, the question is: How ‘good’ are the CNs for redox conduction compared to idealized heme chains with the same sequence of slip- and T-stacked packing motifs, but the fastest known experimental rates (k_et,slip–stackced_ = 1.56 × 10^9^ and k_et,T–stackced_ = 1.25 × 10^8^s^-1^)?^85, 86^

The tetra-, hexa- and octahemes have diffusion constants that are respectively one, two, and three orders of magnitude lower than the idealized heme chains (Figure 14, black bars). Those idealized heme chains with maximal conductivities based on 1–10 ns electron transfers would, in turn, have diffusion constants that are two to four orders of magnitude lower than suggested from the reported conductivities for Omc-S and Z (Figure 14, white bars).

To give a more physically intuitive perspective, Figure 15 shows the length dependent currents that the various CNs could support at a 0.1 V bias with the computed diffusion constants (colored lines) via Eq. 15.^94^ The black open circles and squares in the figure represent the diffusive current through the idealized (TS)_n_ and (TSSTSTS)_n_ chains.

**Figure 15.**
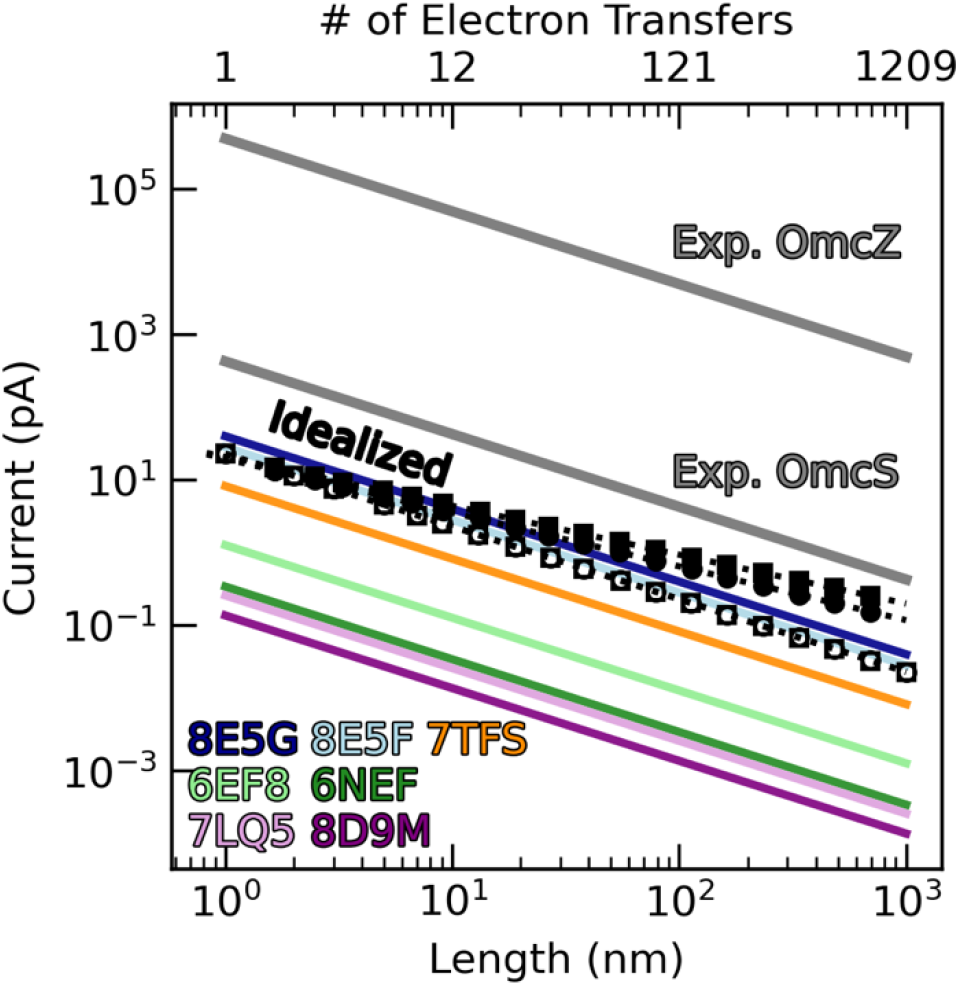
Cytochrome ‘nanowires’ are sub-picoamp conductors. The black unfilled circles and squares show the diffusive current at 0.1 V through the heme chains of Omc-S and Z, but with the fastest experimentally known heme-to-heme electron transfer rates at each step. The black filled circles and squares are the protein-limited steady-state currents for the same idealized Omc-S and Z chains. The sub-optimal diffusive currents at 0.1 V through structurally characterized cytochrome ‘nanowires’ are shown with colored lines. The unphysically large-for-a-cytochrome reported currents at 0.1 V through Omc-S and Z are shown in gray.

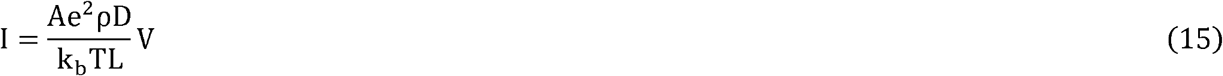

The closed circles and squares in Figure 15 represent the maximal protein-limited current computed using a multi-particle steady-state approach. Because the approach is computationally demanding, an equation for a hopping process (Eq. 16)^136^ was fit (R^2^ = 0.87 − 0.92) to computed fluxes for chains up to 18 steps and then extrapolated for longer filaments.

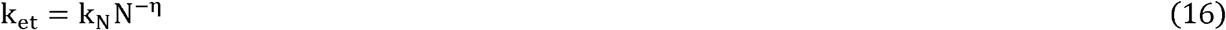

In Eq. 16, k_N_ is the hopping rate and N is the number of hopping steps. k_N_ was 1.4 − 1.5 ×10^8^ s^-1^, nicely reflecting the kinetic constraint of the T-stacked pairs present in either chain. η was 0.67–0.74 for the chains.

Both the diffusive and protein-limited kinetic models agree that a micron-long CN emanating from a microbial cell has a sub-picoamp ampacity. A micron-long chain of either the (TS)_n_ or (TSSTSTS)_n_ heme packing pattern and the fastest known T- and slip-stacked rates at every step can carry a diffusive current at 0.1 V of ∼0.02 pA, or a protein-limited current of ∼0.2 pA. Most of the actual CNs carry less currents.

There is no structural basis for the report that OmcZ transports 3.0 ×10^4^ pA at 0.1 V.^119^ This discrepancy may reflect the fact that the measurement was done on a sample with structural characteristics that are now known to disagree with the CryoEM geometry ^51^, perhaps suggesting the presence of impurities or experimental artifacts.

Taken together, the known CNs are *sufficiently* conductive for cellular respiration, but not *maximally* conductive for a multiheme cytochrome. From a physiological perspective, there is no need to be more conductive, because enzymatic turnover during organic matter oxidation in the cytoplasm and the ferrying of electrons across the periplasm to the CNs by a relay team of proteins likely places the rate-limitation on cellular respiration compared to intra-filament electron transfers.

With the conserved heme chain in CNs being sufficiently conductive regardless of the protein packaging, the latter can be customized for habitat specific interactions without having to re-invent the electron transfer mechanism. CNs are structurally diversified but functionally conserved because evolution has favored robustness over tunability in their design.^41^

## 4. Conclusion

To transport electrons over long distances, nature connects catalytic centers by chains of cofactors. The chain of heme cofactors in a CN must be densely packed to realize weak (≤0.01 eV) electronic couplings for electron transfer. But this dense packing (1) limits the accessible driving forces between adjacent hemes (≤|0.3| eV) by placing them in shared electrostatic microenvironments, (2) introduces strong (≤0.1 eV) redox anti-cooperativities that would accentuate the free energy landscape if it were not for the linear topology of the heme chain, and (3) incurs an entropic penalty that must be offset by tethering the hemes to the polypeptide backbone via thioether linkages.

These linkages physically require T-stacking of hemes for chains longer than three hemes, and in fact, almost every other heme pair in the CNs is of this geometry. T-stacking imposes ∼10-folder smaller electronic couplings and slower electron transfer rates than slip-stacking. As the slowest step in the cascade of redox reactions through CNs, the T-stacked pairs put a structurally-encoded limit on the electron efflux.

Prior experiments and the herein presented computations on all structurally characterized CNs indicate electron transfers between T-stacked hemes is constrained to the nanosecond timescale. Whether the electron flux is in a diffusive or protein-limited steady-state regime, a micron-long CN has an ampacity of ∼0.02–0.2 pA.

This result implies tens to thousands of filaments would be needed to carry the *entire* respiratory current of the microorganism known to produce three of the so-far characterized five CNs. Since the cell likely produces less current than the 1 pA assumed here, and the CNs would only need to carry a fraction of the total current because there are other routes for expelling electrons, the obtained micro-to-milli-Siemen/cm conductivities physically imposed by their multiheme architectures are sufficient for cellular respiration.

But these conductivities are not optimal compared to idealized heme chains with the fastest known rates for each step. There is simply no need for more speed since the intracellular organic matter oxidation and ferrying of electrons across the periplasm to the CNs are likely the metabolic bottlenecks.

This conclusion puts in stark contrast the unphysical nature of the reported milli-Siemens-to-Siemens/cm conductivities for CNs. These yet-to-be replicated measurements appear structurally implausible, biologically irrelevant, and theoretically inexplicable for a multiheme in physiological conditions.

Over the past decade these electrical measurements were first attributed to *e*-pili and then ascribed to cytochromes. Either way, the comment of Tender and co-workers seems appropriate, “To date [2015], the only results inconsistent with redox conduction [through cytochromes] occurring in *G. sulfurreducens* biofilms are those reported by Malvankar *et al*., which have been addressed elsewhere [36] and which have not been corroborated by others.”^29^

