## Supplementary Material for "Cytochrome ‘Nanowires’ are Physically Limited to Sub-Picoamp Currents that Suffice for Cellular Respiration"

### 1. Supplementary Figures

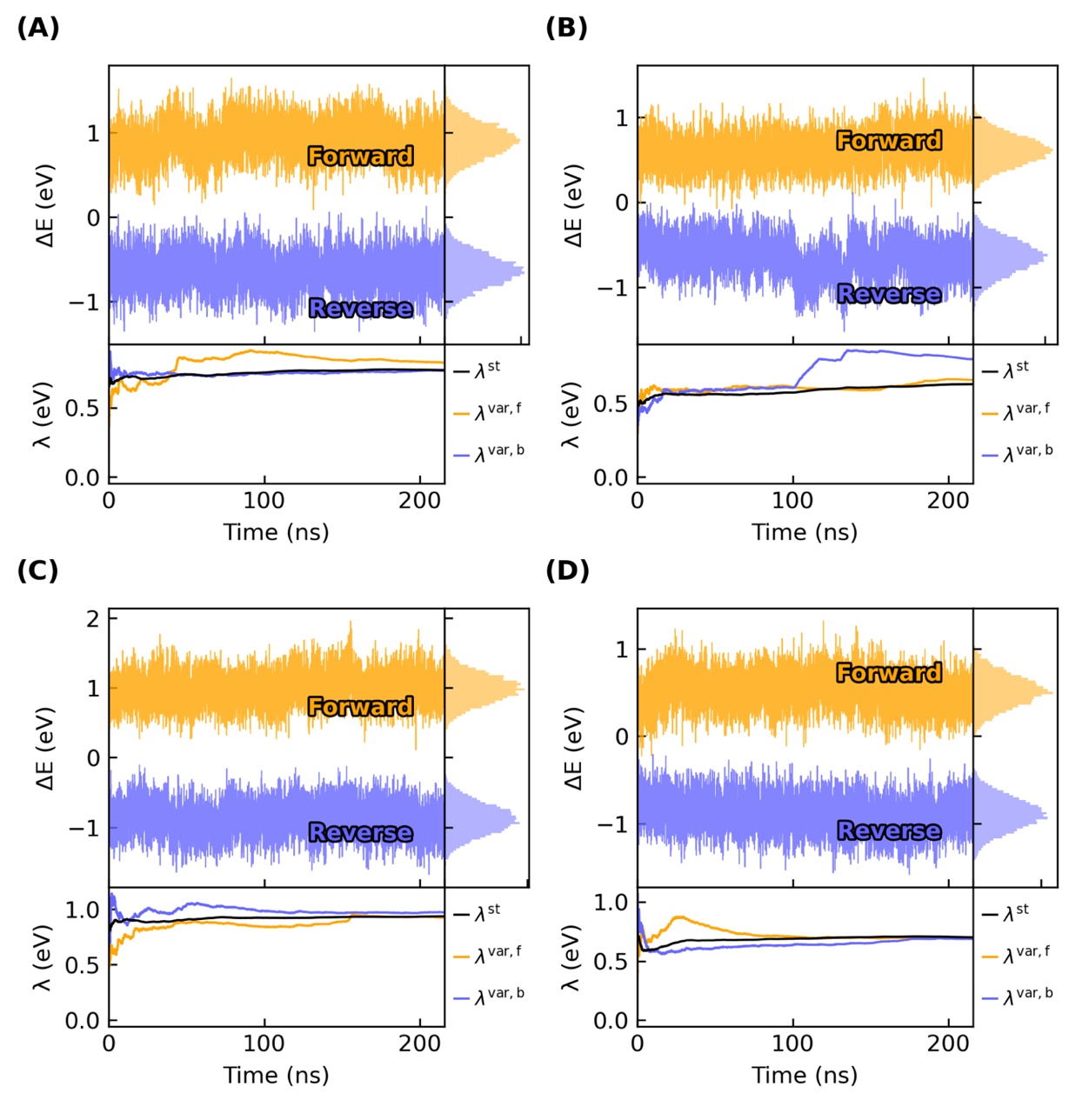

**Supplementary Figure 1.** Fluctuations and convergence of the vertical energy gap for the reaction R,O → O,R [R = reduced, O = oxidized] between Hemes (A) #1 and #2, (B) #2 and #3, (C) #3 and #4, and (D) #4 to 1’ (first heme of the next subunit). The simulations were previously described in.^1^ The convergence plots show the accumulated average and variances over the trajectories.

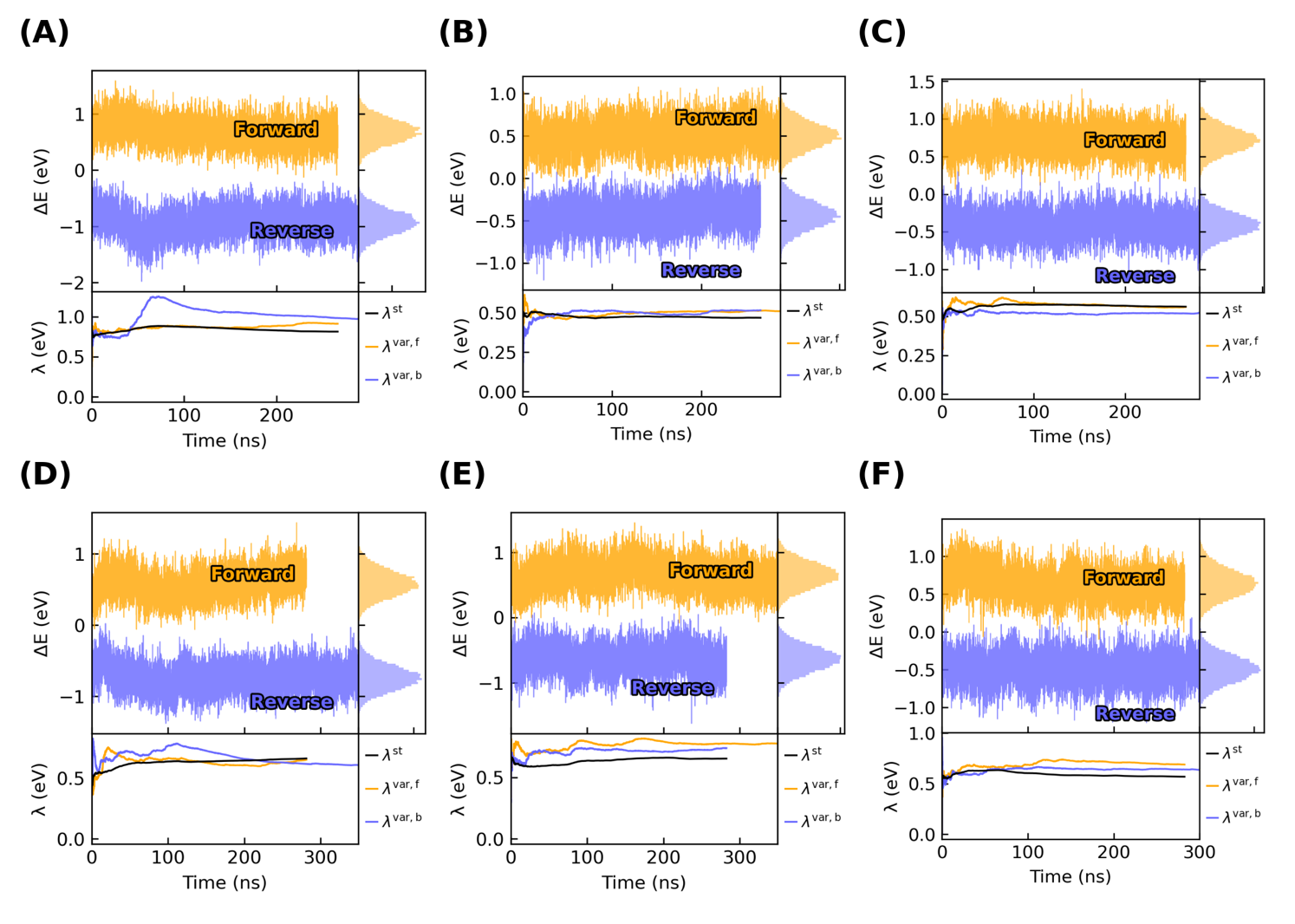

**Supplementary Figure 2.** Fluctuations and convergence of the vertical energy gap for the reaction R,O → O,R [R = reduced, O = oxidized] between Hemes (A) #1 and #2, (B) #2 and #3, (C) #3 and #4, (D) #4 and #5, (E) #5 to #6, and (F) #6 to 1’ (first heme of the next subunit). The simulations were previously described in ^1^. The convergence plots show the accumulated average and variances over the trajectories.

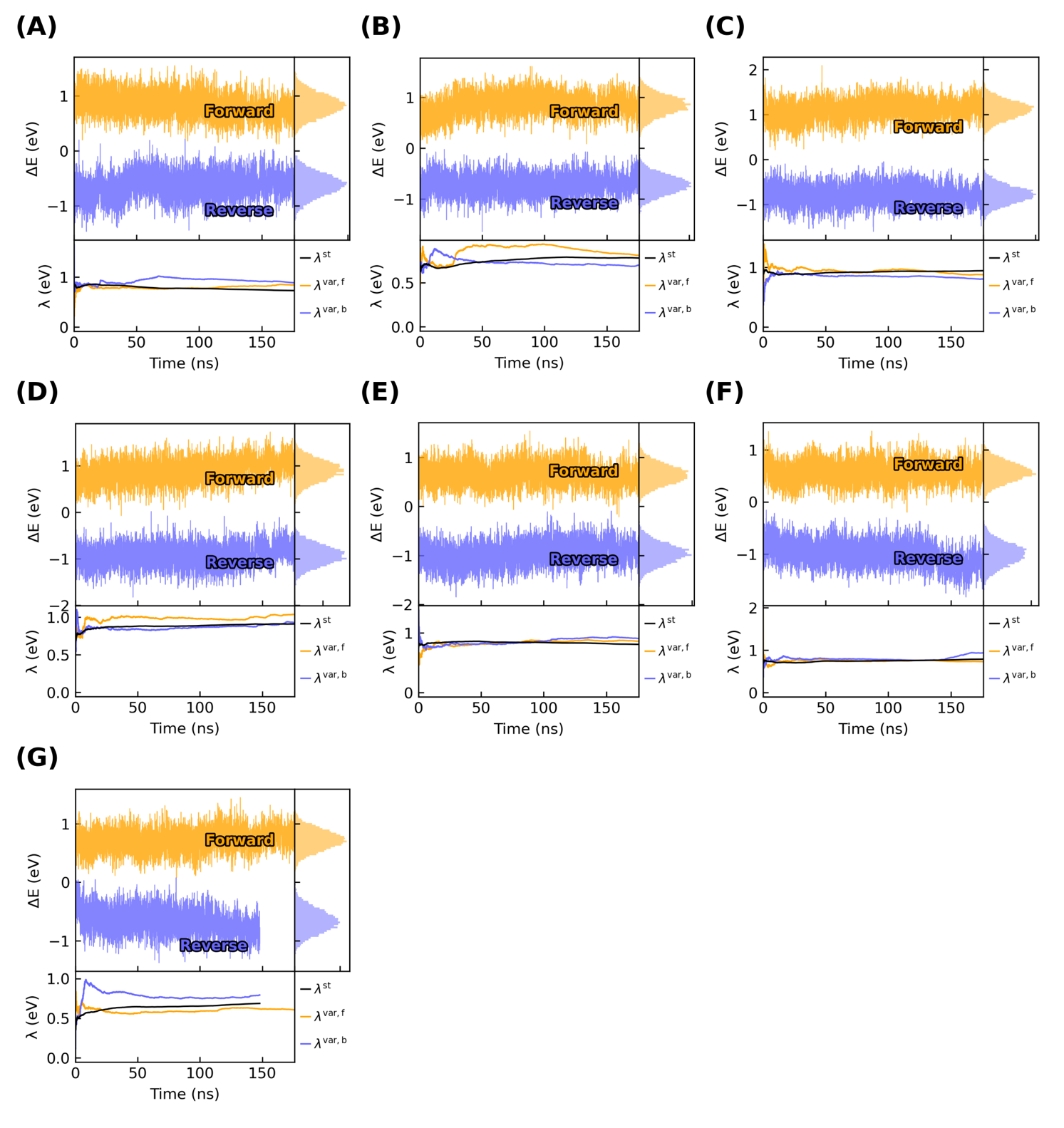

**Supplementary Figure 3.** Fluctuations and convergence of the vertical energy gap for the reaction R,O → O,R [R = reduced, O = oxidized] between Hemes (A) #1 and #2, (B) #2 and #3, (C) #3 and #4, (D) #4 and #5, (E) #5 to #6, (F) #6 to 7, and (G) 7 to 1’ (first heme of the next subunit). The simulations were previously described in ^1^. The convergence plots show the accumulated average and variances over the trajectories.

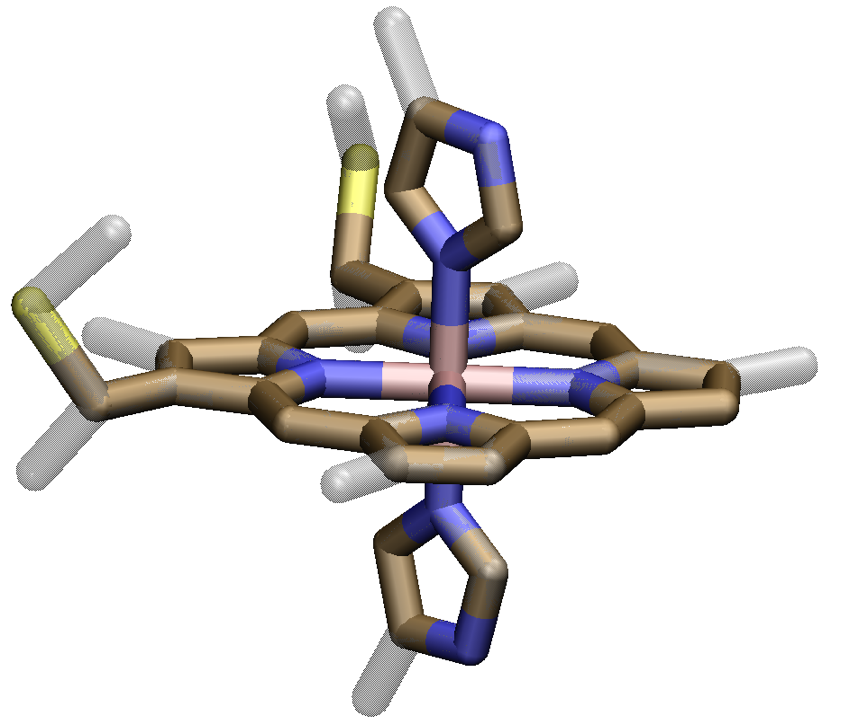

**Supplementary Figure 4.** Model of the heme cofactor used for optical spectra simulations. The atoms with a ‘ghostly’ appearance are methyl groups that were removed for the most truncated model. Structures with and without these methyl groups were optimized with the B3LYP hybrid density functional and mixed basis set (LANL2DZ for Fe and 6-31G(d) for H, C, N and S). No imaginary frequencies were found in a harmonic vibrational analysis. Hydrogen atoms are not shown for clarity.

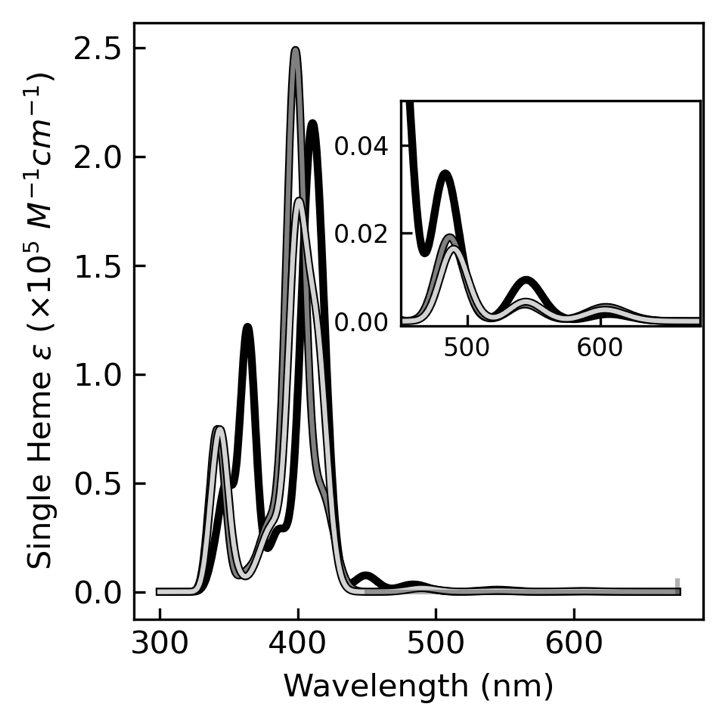

**Supplementary Figure 5.** UV-vis spectra of the fully optimized heme models with (black) and without (light gray) methyl substituents. The dark gray spectrum is for the unmethylated heme model with only the hydrogens replacing the methyl groups re-optimized after truncation. The difference between the black and dark gray spectra shows the direct electronic effect of the methyl substituents (a ~10 nm blue-shift of the Soret band). The difference between the dark gray and light gray spectra reflects geometrical changes elsewhere in the molecule due to the hydrogen-for-methyl substitution. The spectra were computed in the framework of time dependent density functional theory using the BLYP function and a mixed (LANL2DZ for Fe and 6-31G(d) for H, C, N, and S) basis set. The spectra were generated by fitting Gaussians to the simulated wavelengths with a broadening factor of 0.07 eV (1330 cm^-1^).

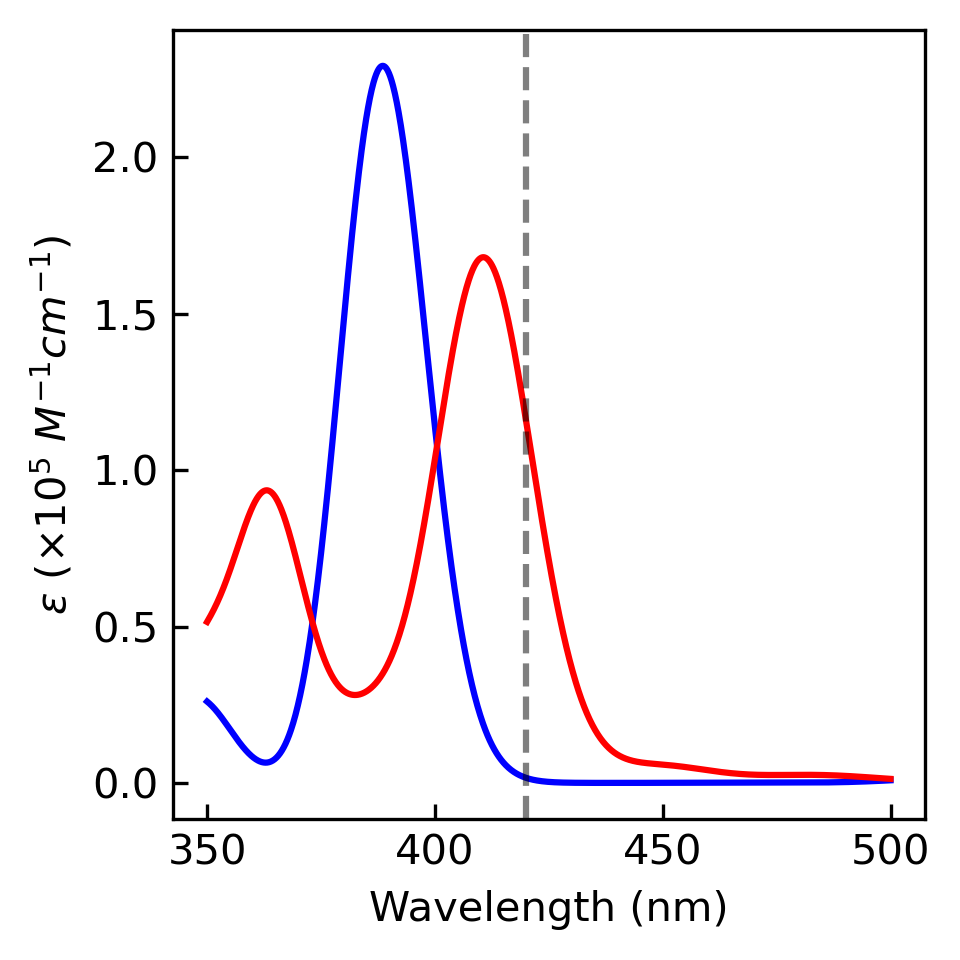

**Supplementary Figure 6.** UV-vis spectra of the fully optimized heme model with methyl substituents using the B3LYP (blue) or BLYP (red) density functional with the same basis set (LANL2DZ for Fe, 6-31G(d) for H, C, N, and S) for the time-dependent density functional theory calculation. The same optimized geometry (B3LYP/(LANL2DZ for Fe, 6-31G(d) for H, C, N, S) was used for both TD-DFT calculations. The dotted vertical line at 420 nm is the expected Soret absorption maximum for a reduced bis-histidine-ligated *c*-type heme.

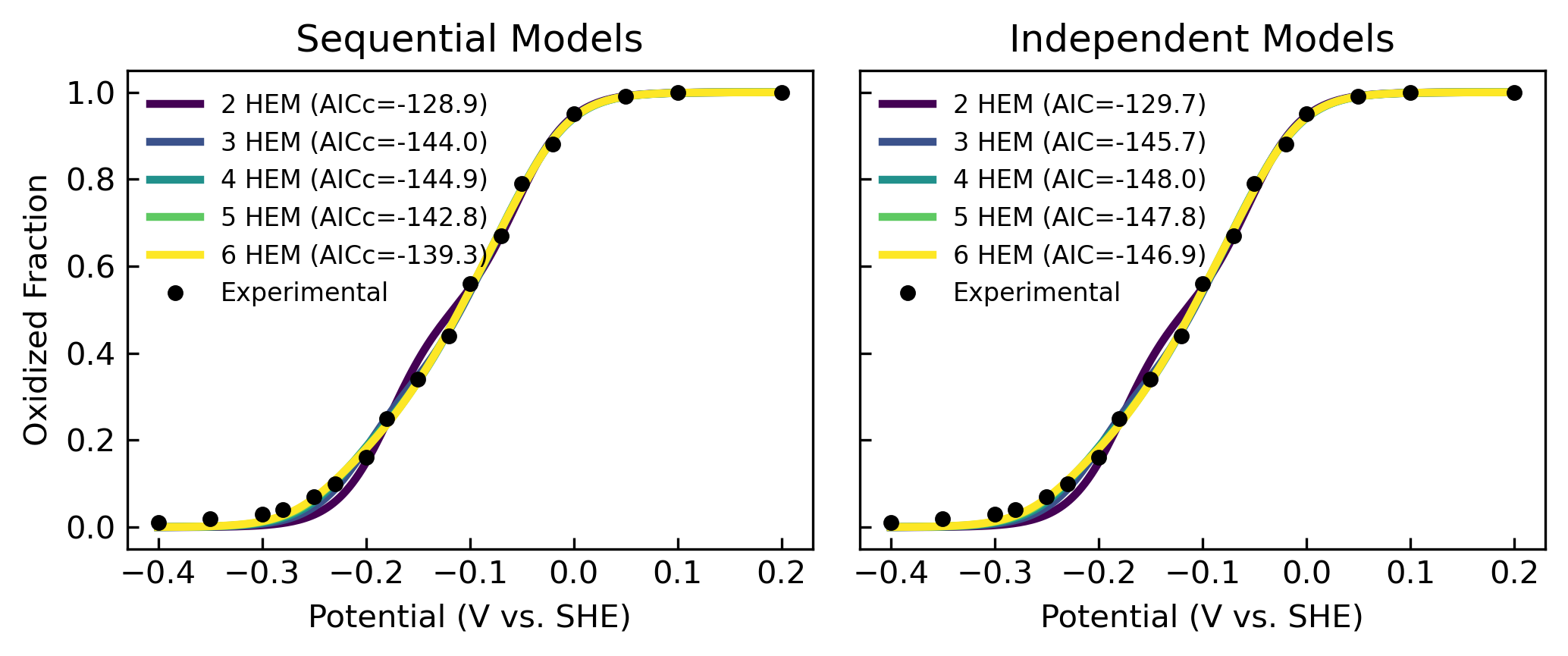

**Supplementary Figure 7.** The spectroelectrochemical data of ^2^ is fit with models of independently or sequentially coupled redox reactions that each have increasing number of hemes . The redox potential of each heme is a fitting parameter. The corrected (for small data sets) Akaike information criterion (AICc) is used to judge the balance between model fit and model complexity. The model with the most negative AICc value with the fewest fitting parameters is the best model.

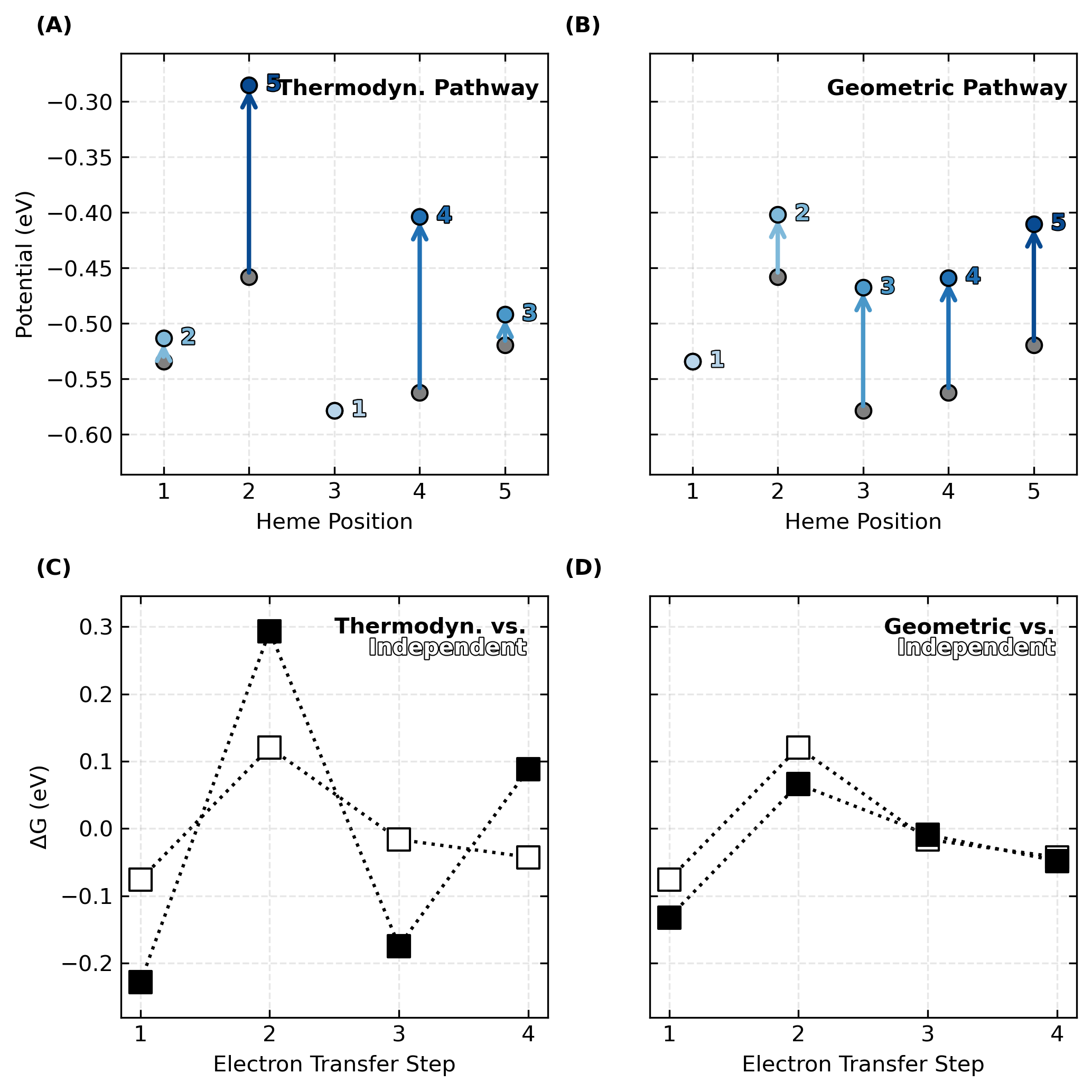

**Supplementary Figure 8.** Shifts in microscopic redox potentials and heme-to-heme reaction free energies in OmcE due to redox anti-cooperativity. Panels (A) and (B) show the redox potentials without (gray) and with (blue) heme-heme interactions. The hemes are either oxidized (A) in order of thermodynamic preference, or (B) the geometrical sequence imposed by the topology of the cytochrome ‘nanowire.’ Panels (C) and (D) show how the shifts in (A) and (B) respectively change the free energy landscape for multi-step electron transfer through the filament.

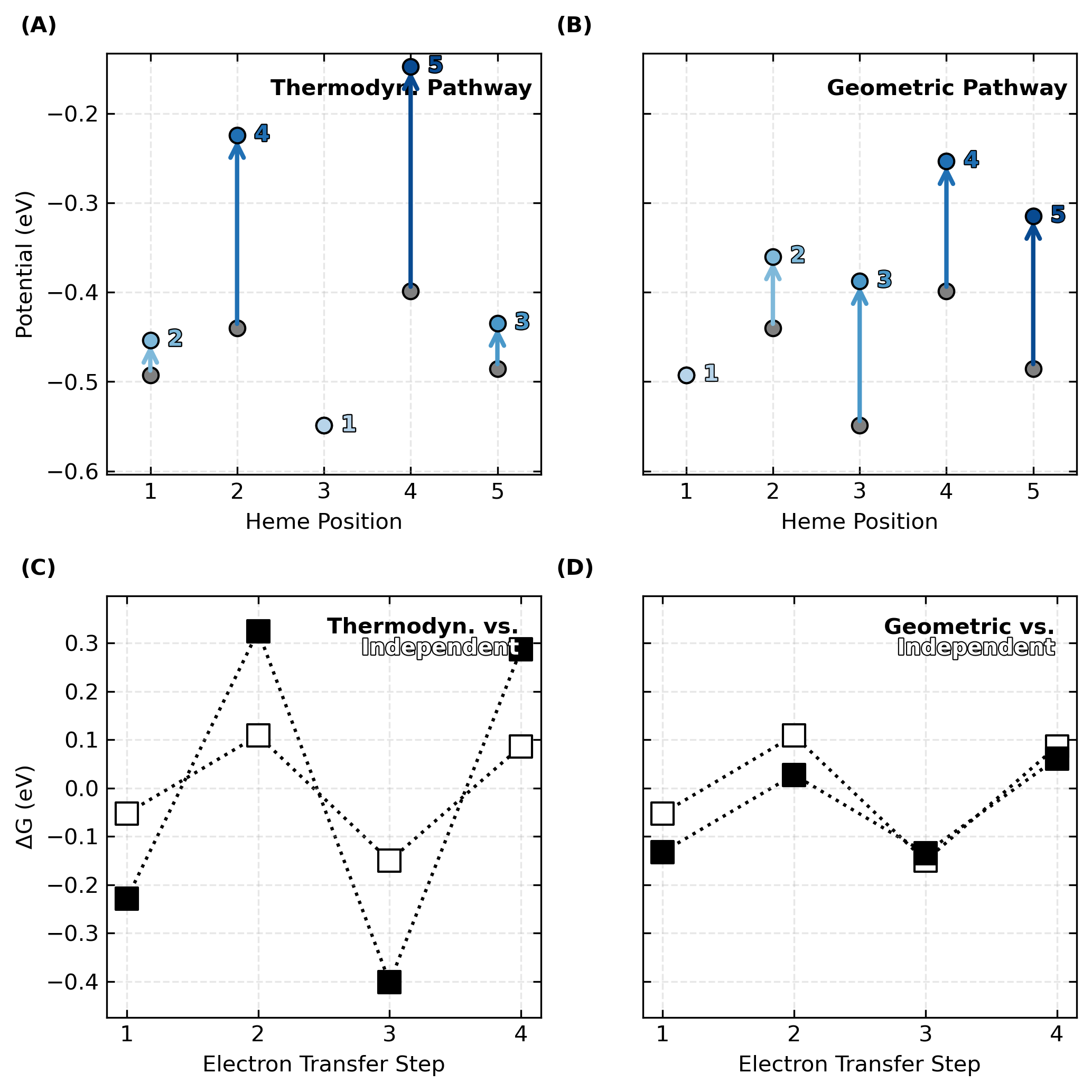

**Supplementary Figure 9. S**hifts in microscopic redox potentials and heme-to-heme reaction free energies in A3MW92 due to redox anti-cooperativity. Panels (A) and (B) show the redox potentials without (gray) and with (blue) heme-heme interactions. The hemes are either oxidized (A) in order of thermodynamic preference, or (B) the geometrical sequence imposed by the topology of the cytochrome ‘nanowire.’ Panels (C) and (D) show how the shifts in (A) and (B) respectively change the free energy landscape for multi-step electron transfer through the filament.

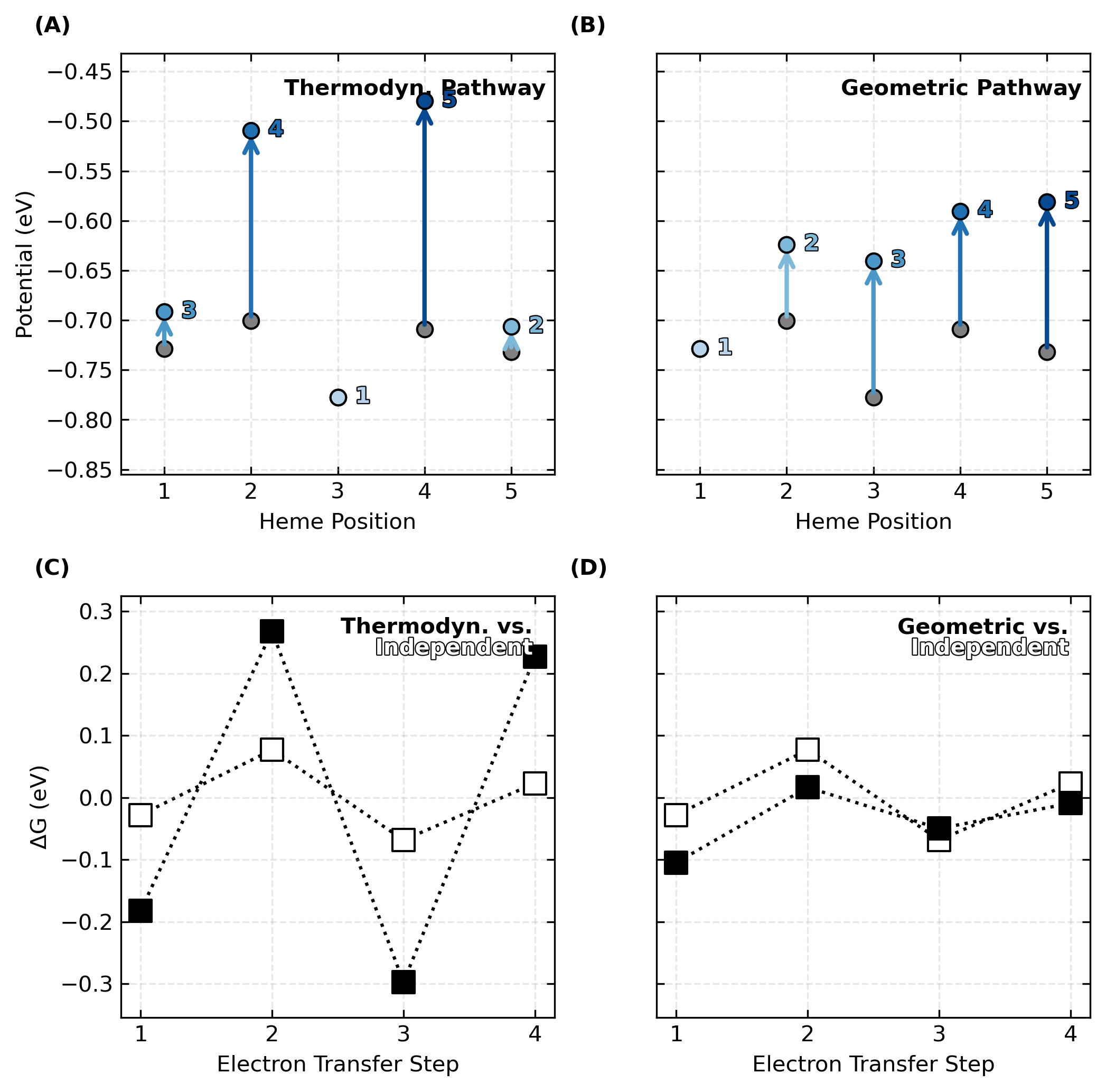

**Supplementary Figure 10.** Shifts in microscopic redox potentials and heme-to-heme reaction free energies in F2KMU8 due to redox anti-cooperativity. Panels (A) and (B) show the redox potentials without (gray) and with (blue) heme-heme interactions. The hemes are either oxidized (A) in order of thermodynamic preference, or (B) the geometrical sequence imposed by the topology of the cytochrome ‘nanowire.’ Panels (C) and (D) show how the shifts in (A) and (B) respectively change the free energy landscape for multi-step electron transfer through the filament.

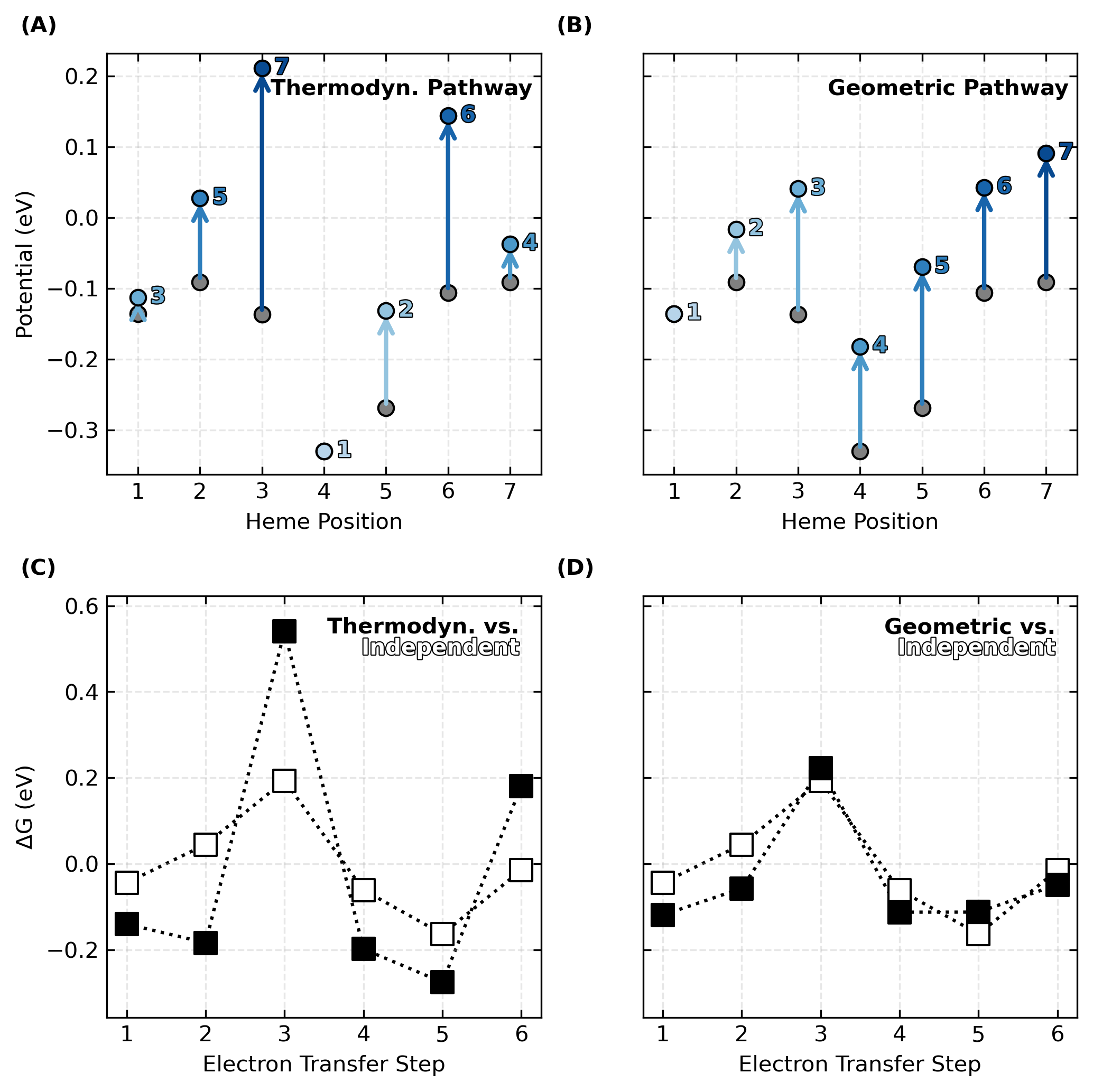

**Supplementary Figure 11.** Shifts in microscopic redox potentials and heme-to-heme reaction free energies in OmcS (PDB 6EF8) due to redox anti-cooperativity. Panels (A) and (B) show the redox potentials without (gray) and with (blue) heme-heme interactions. The hemes are either oxidized (A) in order of thermodynamic preference, or (B) the geometrical sequence imposed by the topology of the cytochrome ‘nanowire.’ Panels (C) and (D) show how the shifts in (A) and (B) respectively change the free energy landscape for multi-step electron transfer through the filament.

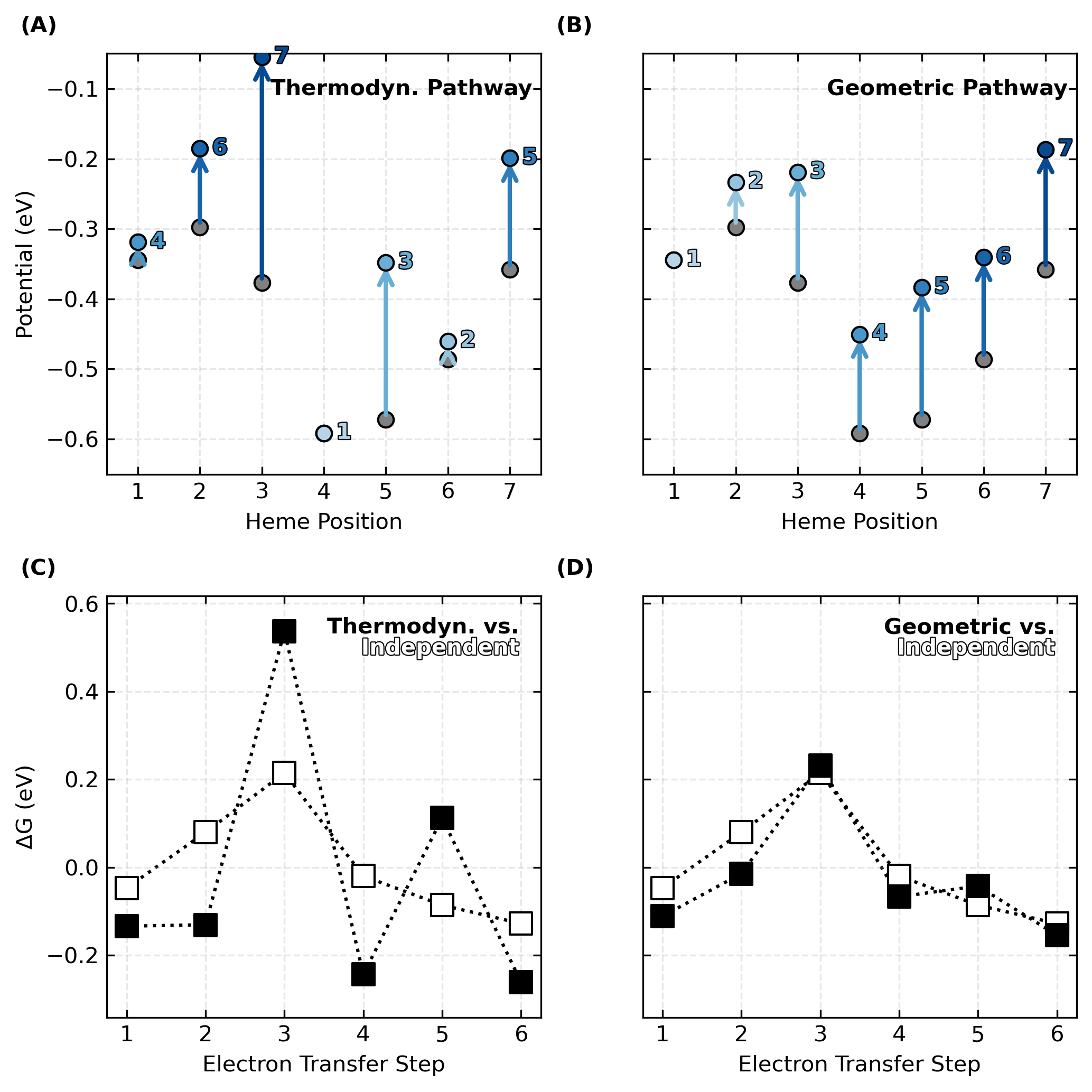

**Supplementary Figure 12.** Shifts in microscopic redox potentials and heme-to-heme reaction free energies in OmcS (PDB 6NEF) due to redox anti-cooperativity. Panels (A) and (B) show the redox potentials without (gray) and with (blue) heme-heme interactions. The hemes are either oxidized (A) in order of thermodynamic preference, or (B) the geometrical sequence imposed by the topology of the cytochrome ‘nanowire.’ Panels (C) and (D) show how the shifts in (A) and (B) respectively change the free energy landscape for multi-step electron transfer through the filament.

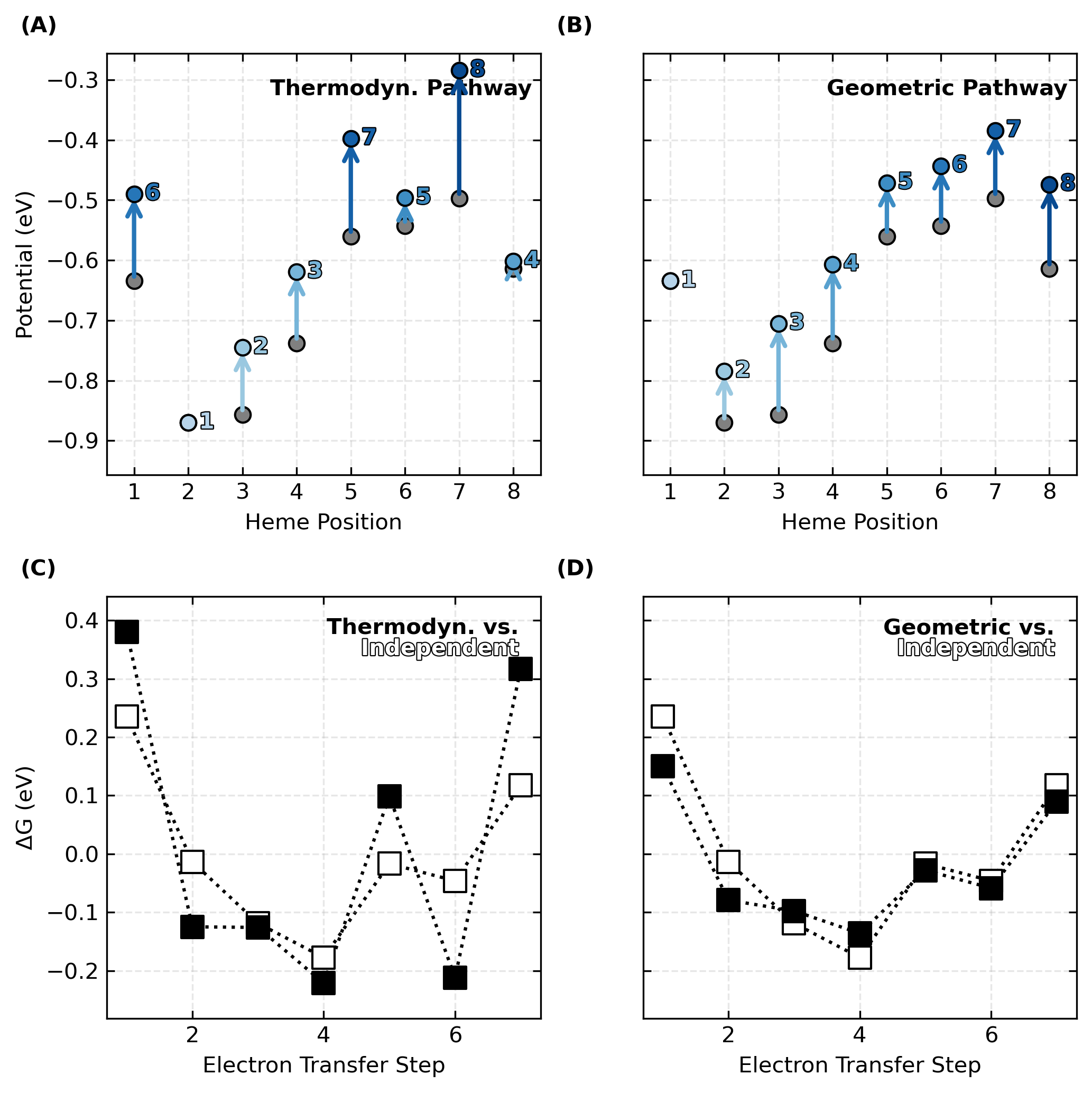

**Supplementary Figure 13.** Shifts in microscopic redox potentials and heme-to-heme reaction free energies in OmcZ (PDB 7LQ5) due to redox anti-cooperativity. Panels (A) and (B) show the redox potentials without (gray) and with (blue) heme-heme interactions. The hemes are either oxidized (A) in order of thermodynamic preference, or (B) the geometrical sequence imposed by the topology of the cytochrome ‘nanowire.’ Panels (C) and (D) show how the shifts in (A) and (B) respectively change the free energy landscape for multi-step electron transfer through the filament.

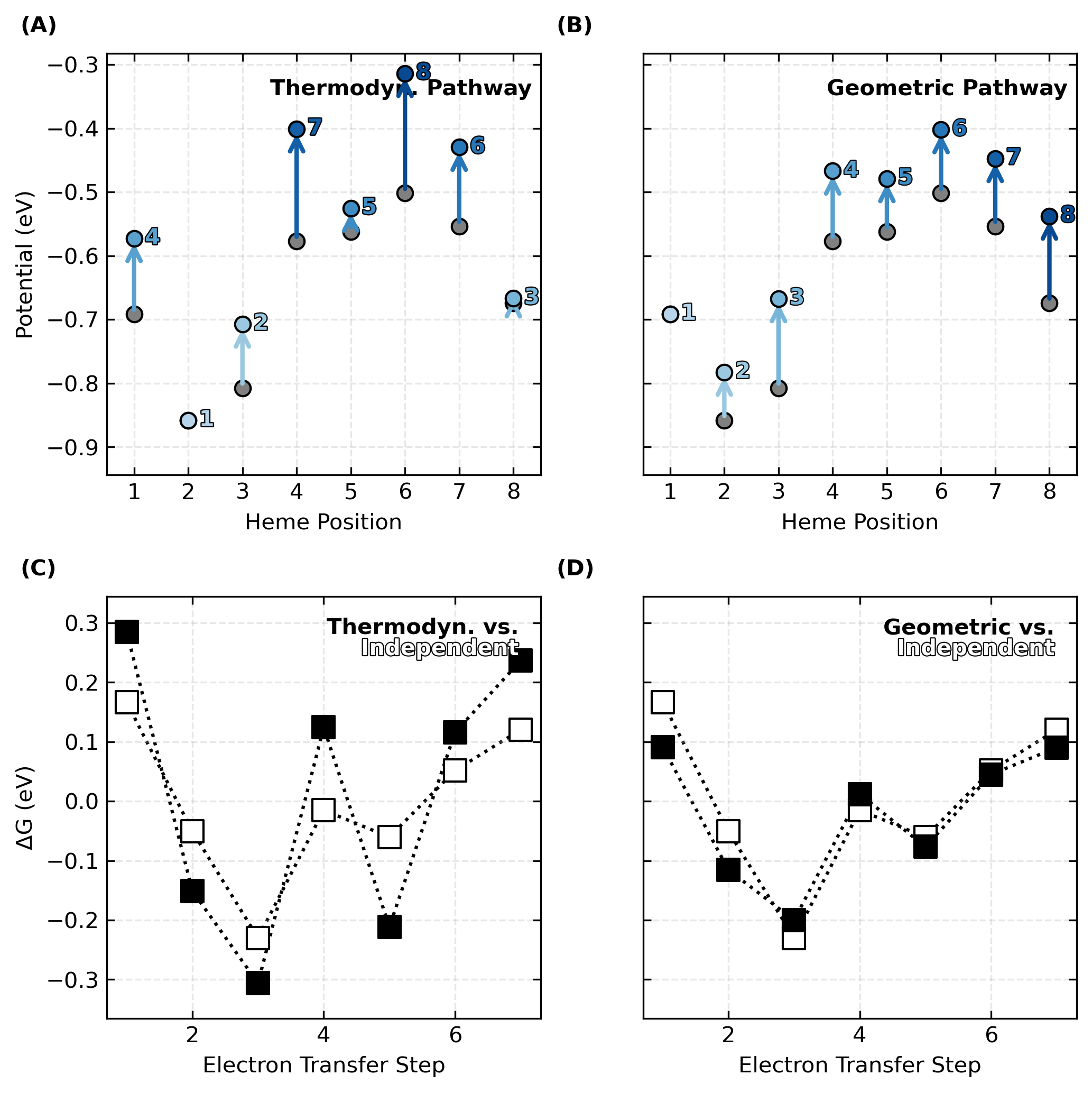

**Supplementary Figure 14.** Shifts in microscopic redox potentials and heme-to-heme reaction free energies in OmcZ (PDB 8D9M) due to redox anti-cooperativity. Panels (A) and (B) show the redox potentials without (gray) and with (blue) heme-heme interactions. The hemes are either oxidized (A) in order of thermodynamic preference, or (B) the geometrical sequence imposed by the topology of the cytochrome ‘nanowire.’ Panels (C) and (D) show how the shifts in (A) and (B) respectively change the free energy landscape for multi-step electron transfer through the filament.

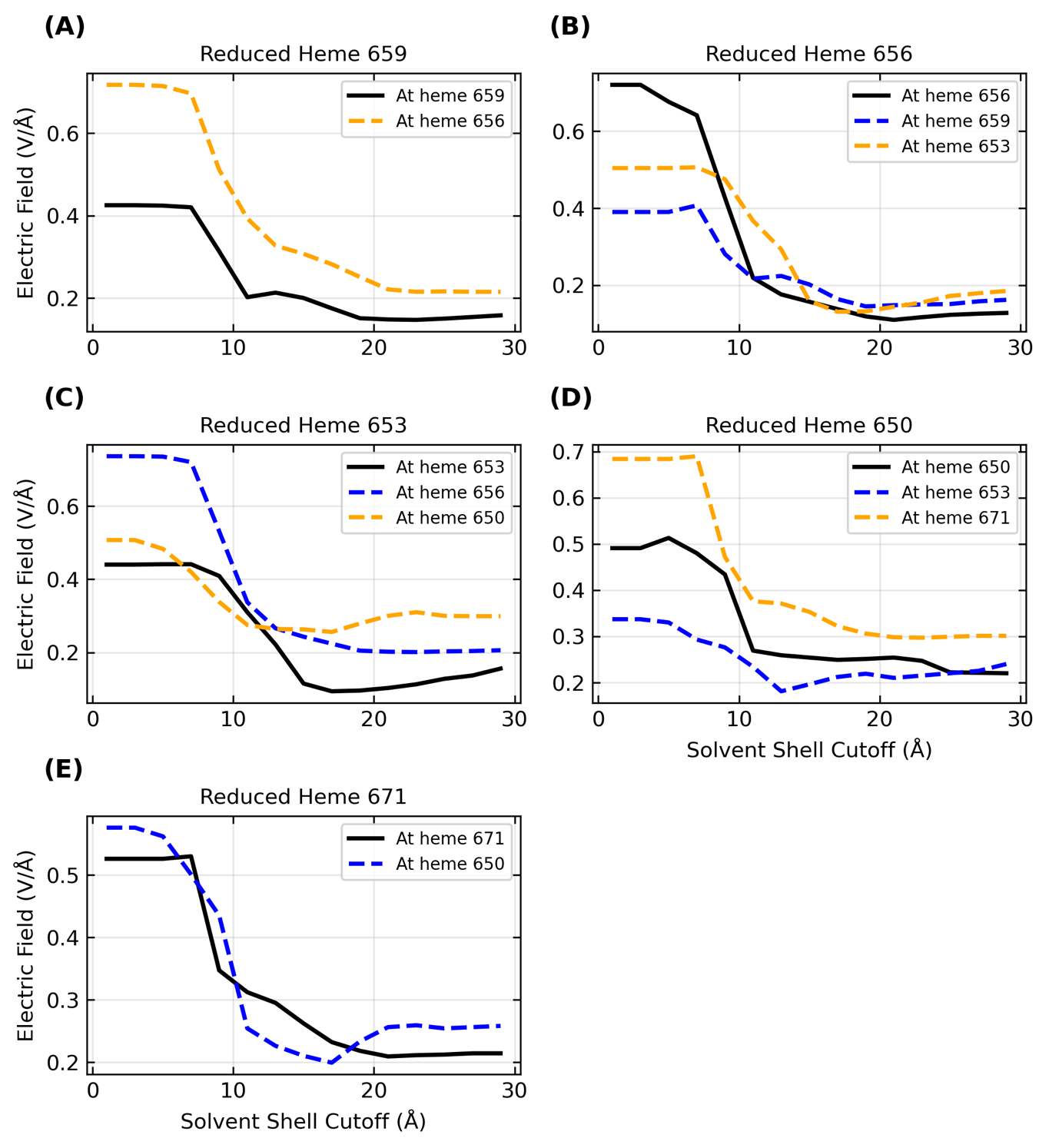

**Supplementary Figure 15.** Dependence of the protein-water electric field at the heme-Fe centers of OmcE as a function of the cutoff distance for including an explicit aqueous solvent with sufficient sodium counterions for overall charge neutrality. The hemes are numbered in the figure based on their residue IDs in the structure used for previously published molecular dynamics simulations.

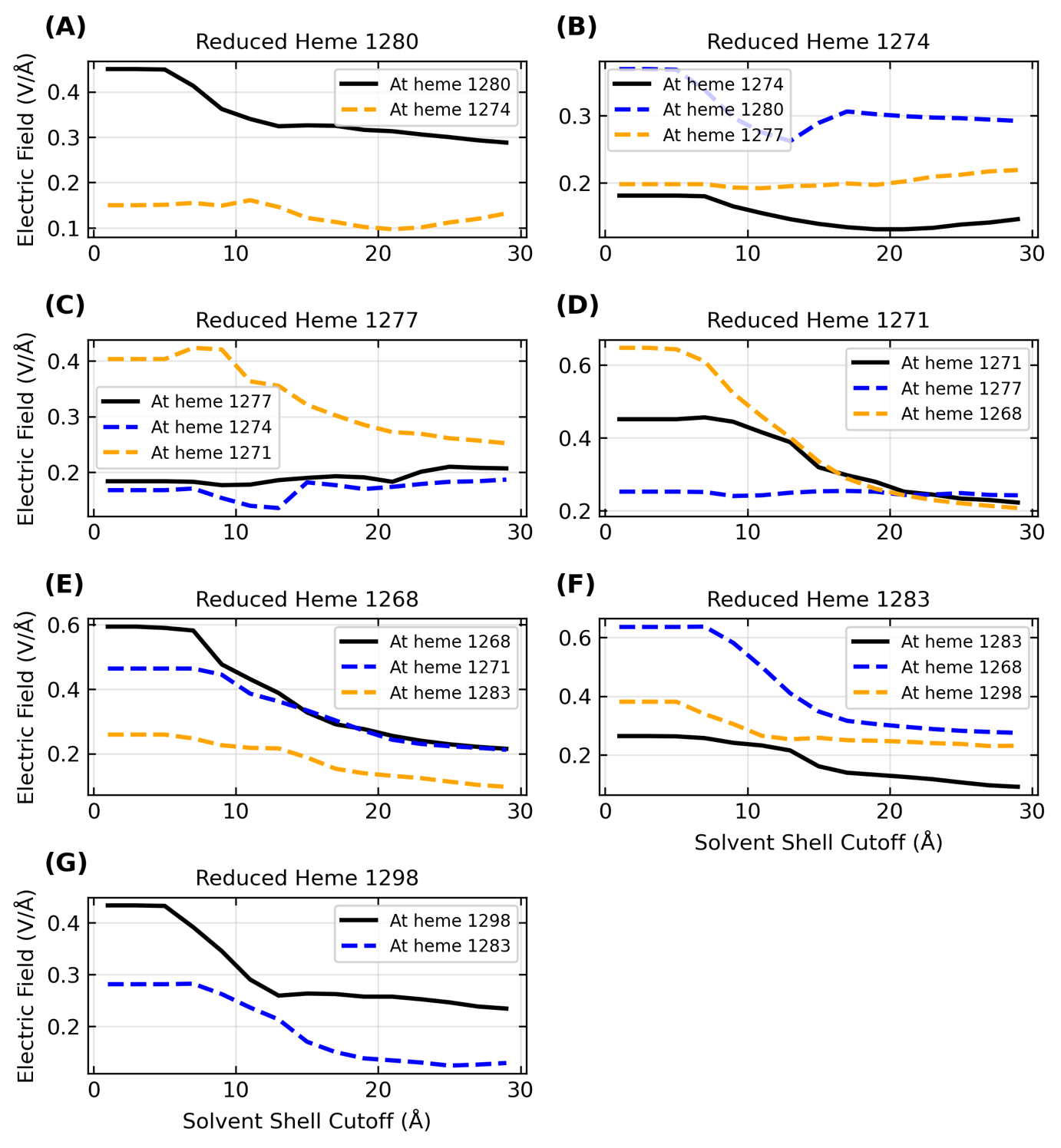
**Supplementary Figure 16.** Dependence of the protein-water electric field at the heme-Fe centers of OmcS as a function of the cutoff distance for including an explicit aqueous solvent with sufficient sodium counterions for overall charge neutrality. The hemes are numbered in the figure based on their residue IDs in the structure used for previously published molecular dynamics simulations.

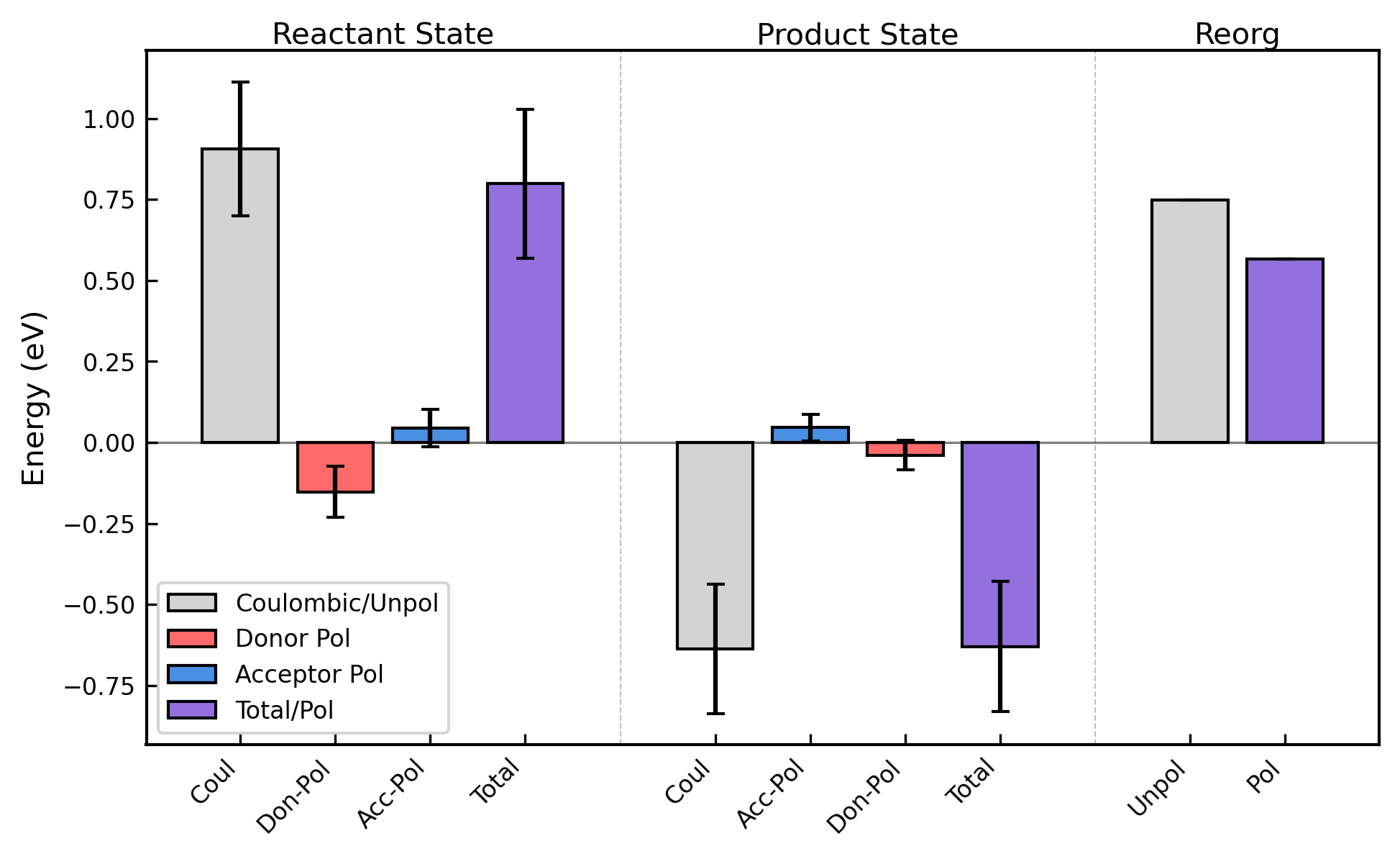

**Supplementary Figure 17.** Contributions to the vertical energy gap for the Heme 1 → 2 (forward) or 2 → 1 (reverse) electron transfer in OmcE from Coulombic interactions, as well as donor (Don) and acceptor (Acc) polarization (Pol). The effect of considering the polarization contributions on the reorganization energy is also shown.

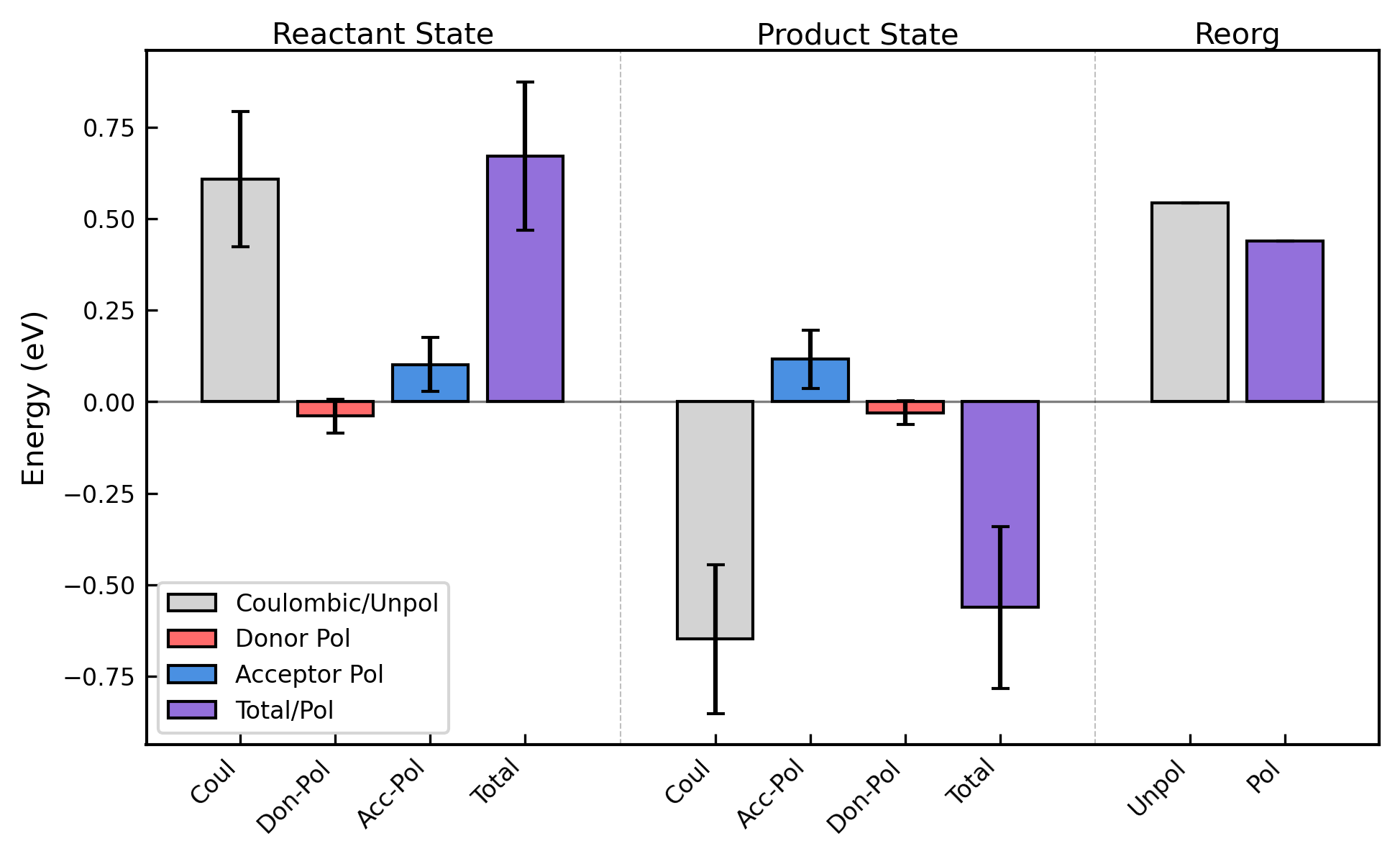

**Supplementary Figure 18.** Contributions to the vertical energy gap for the Heme 2 → 3 (forward) or 3 → 2 (reverse) electron transfer in OmcE from Coulombic interactions, as well as donor (Don) and acceptor (Acc) polarization (Pol). The effect of considering the polarization contributions on the reorganization energy is also shown.

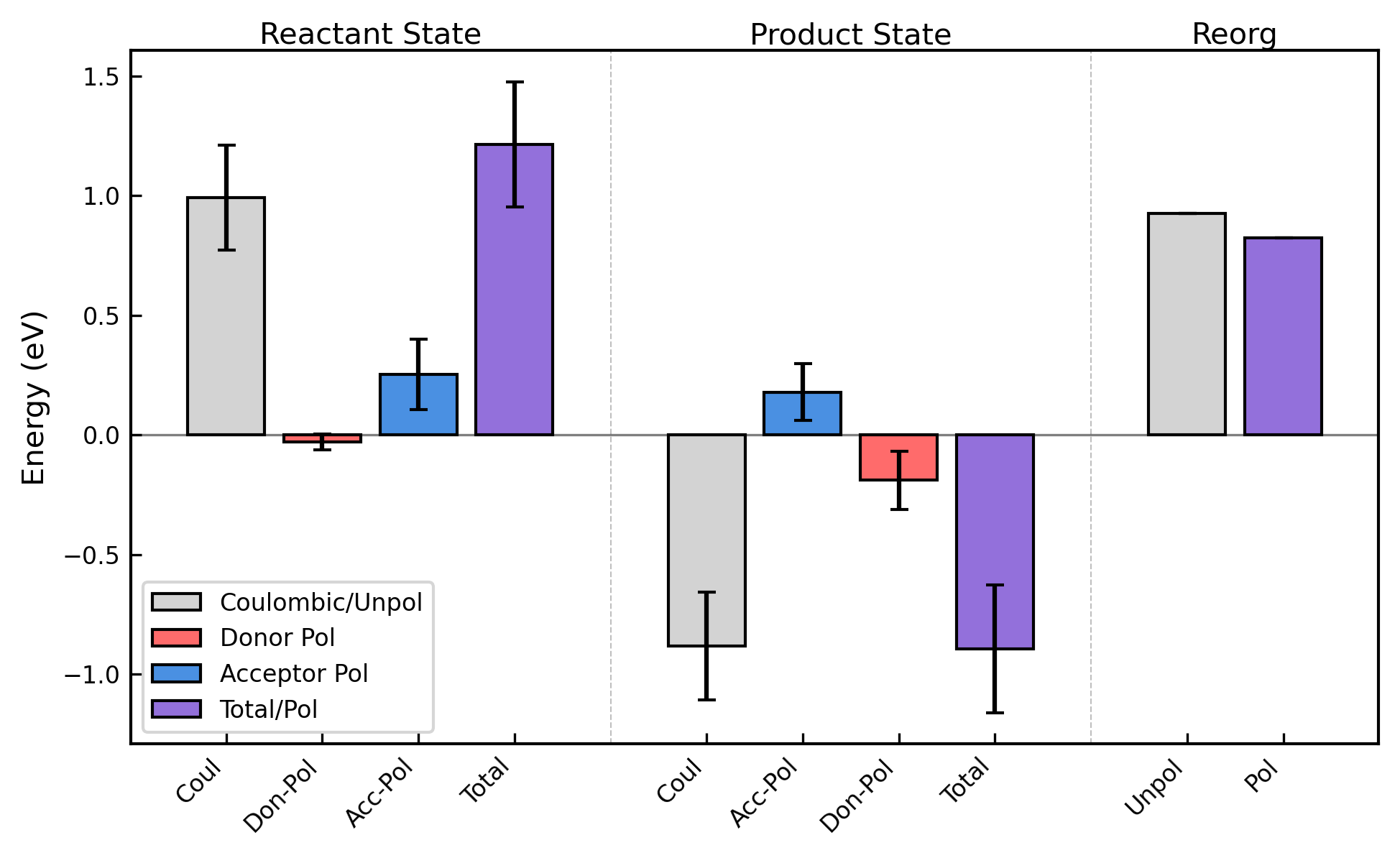

**Supplementary Figure 19.** Contributions to the vertical energy gap for the Heme 3 → 4 (forward) or 4 → 3 (reverse) electron transfer in OmcE from Coulombic interactions, as well as donor (Don) and acceptor (Acc) polarization (Pol). The effect of considering the polarization contributions on the reorganization energy is also shown.

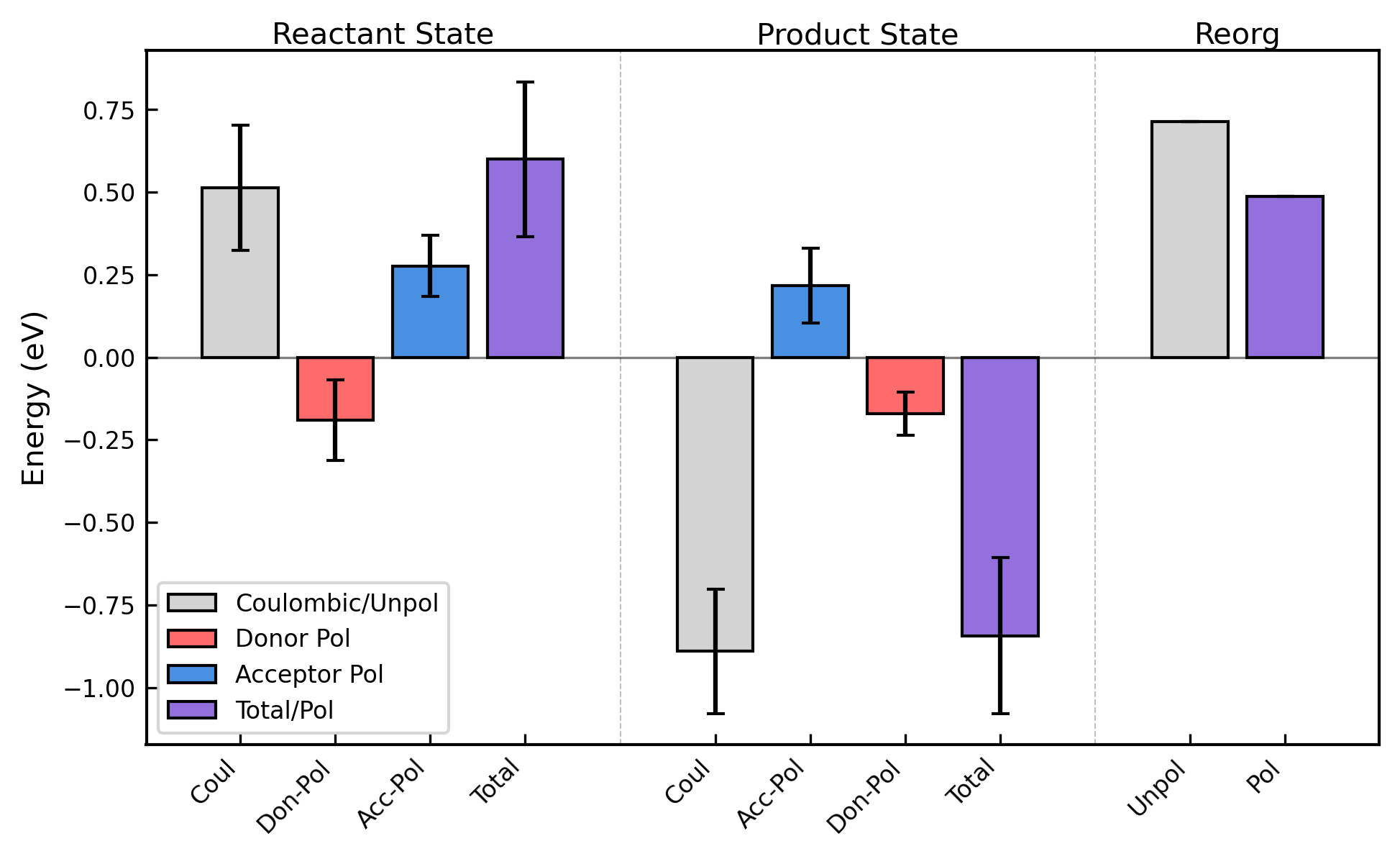

**Supplementary Figure 20.** Contributions to the vertical energy gap for the Heme 4 → 1’ (forward) or 1’ → 4 (reverse) electron transfer in OmcE from Coulombic interactions, as well as donor (Don) and acceptor (Acc) polarization (Pol). The effect of considering the polarization contributions on the reorganization energy is also shown.

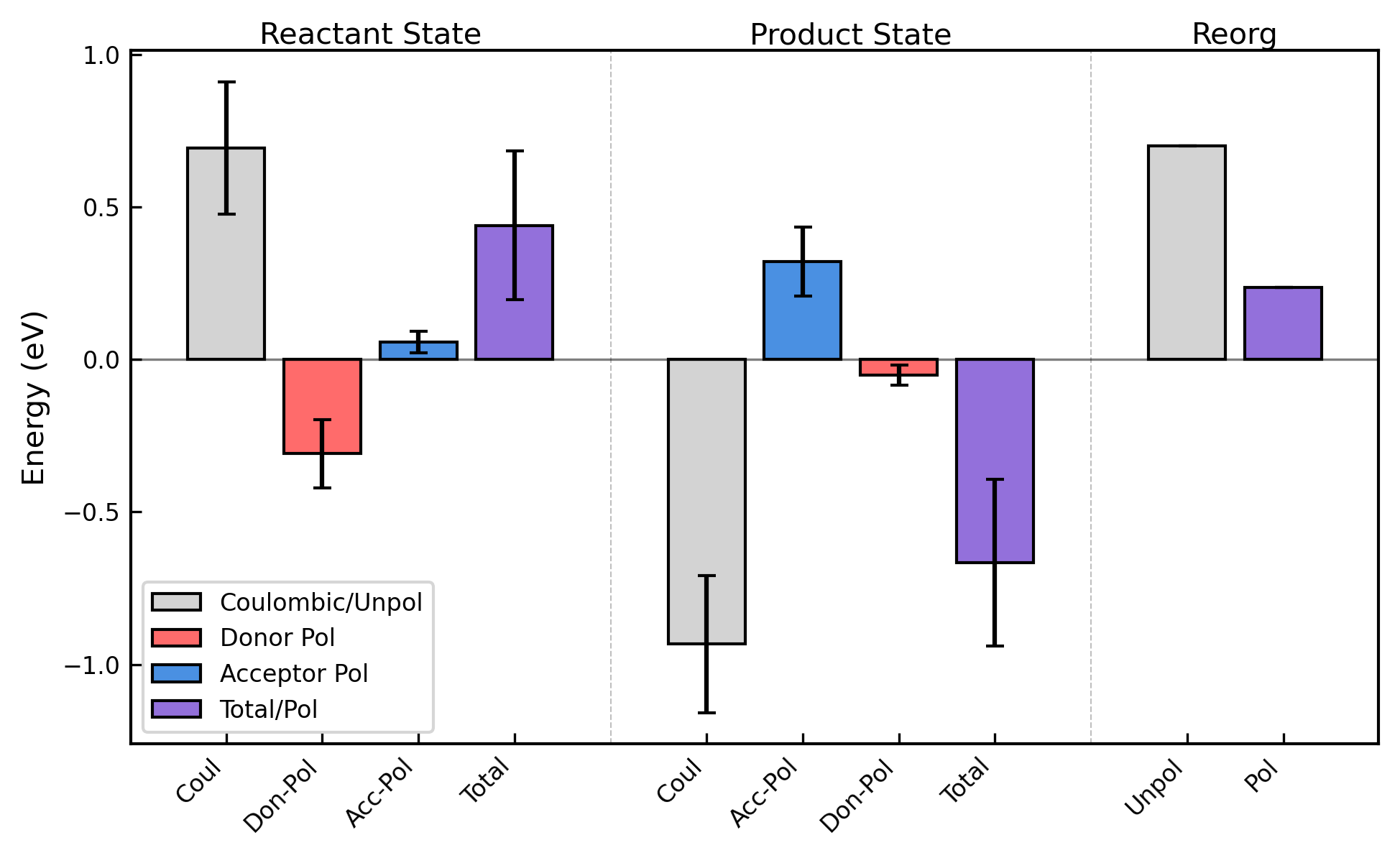

**Supplementary Figure 21.** Contributions to the vertical energy gap for the Heme 1 → 2 (forward) or 2 → 1 (reverse) electron transfer in OmcS from Coulombic interactions, as well as donor (Don) and acceptor (Acc) polarization (Pol). The effect of considering the polarization contributions on the reorganization energy is also shown.

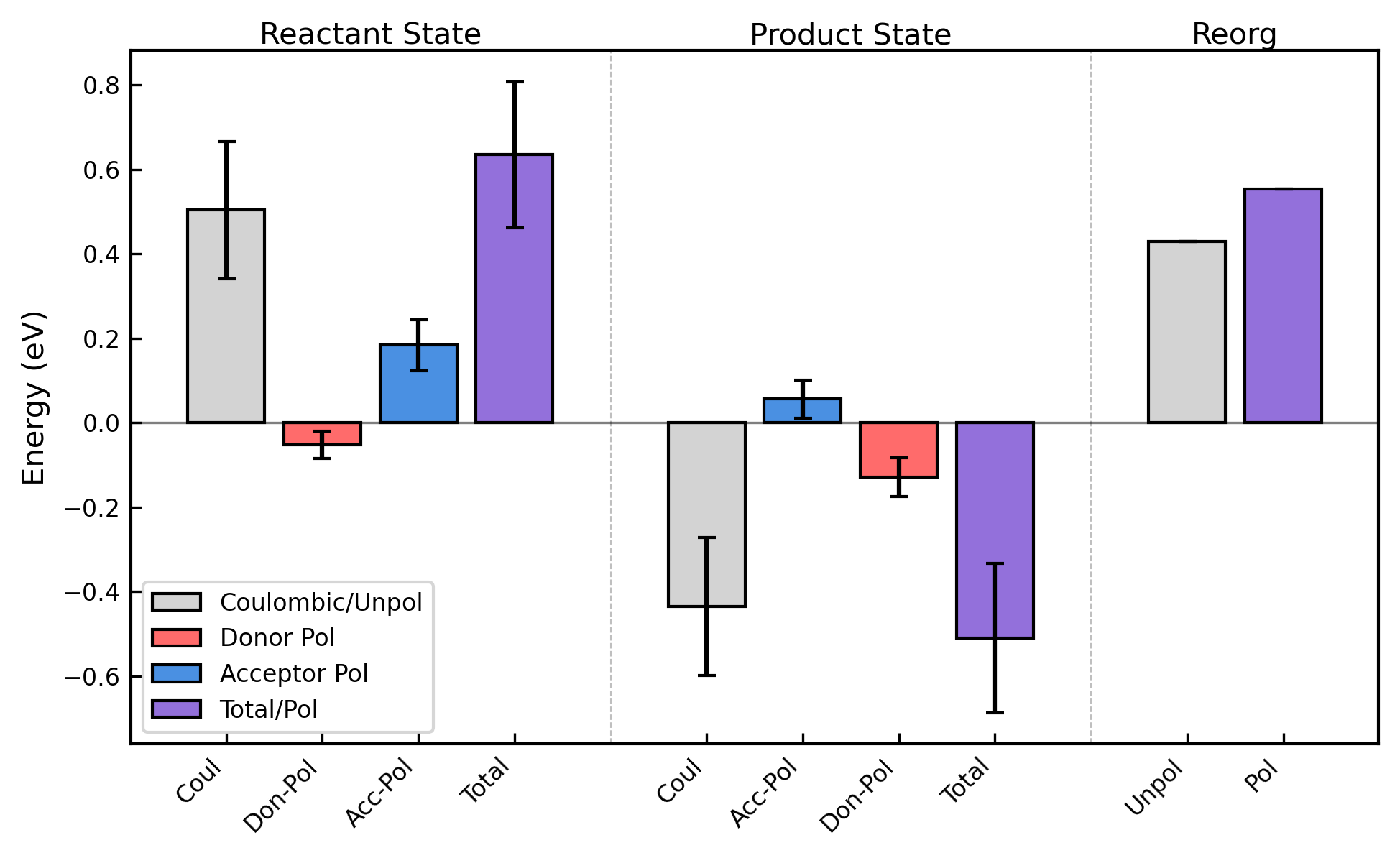

**Supplementary Figure 22.** Contributions to the vertical energy gap for the Heme 2 → 3 (forward) or 3 → 2 (reverse) electron transfer in OmcS from Coulombic interactions, as well as donor (Don) and acceptor (Acc) polarization (Pol). The effect of considering the polarization contributions on the reorganization energy is also shown.

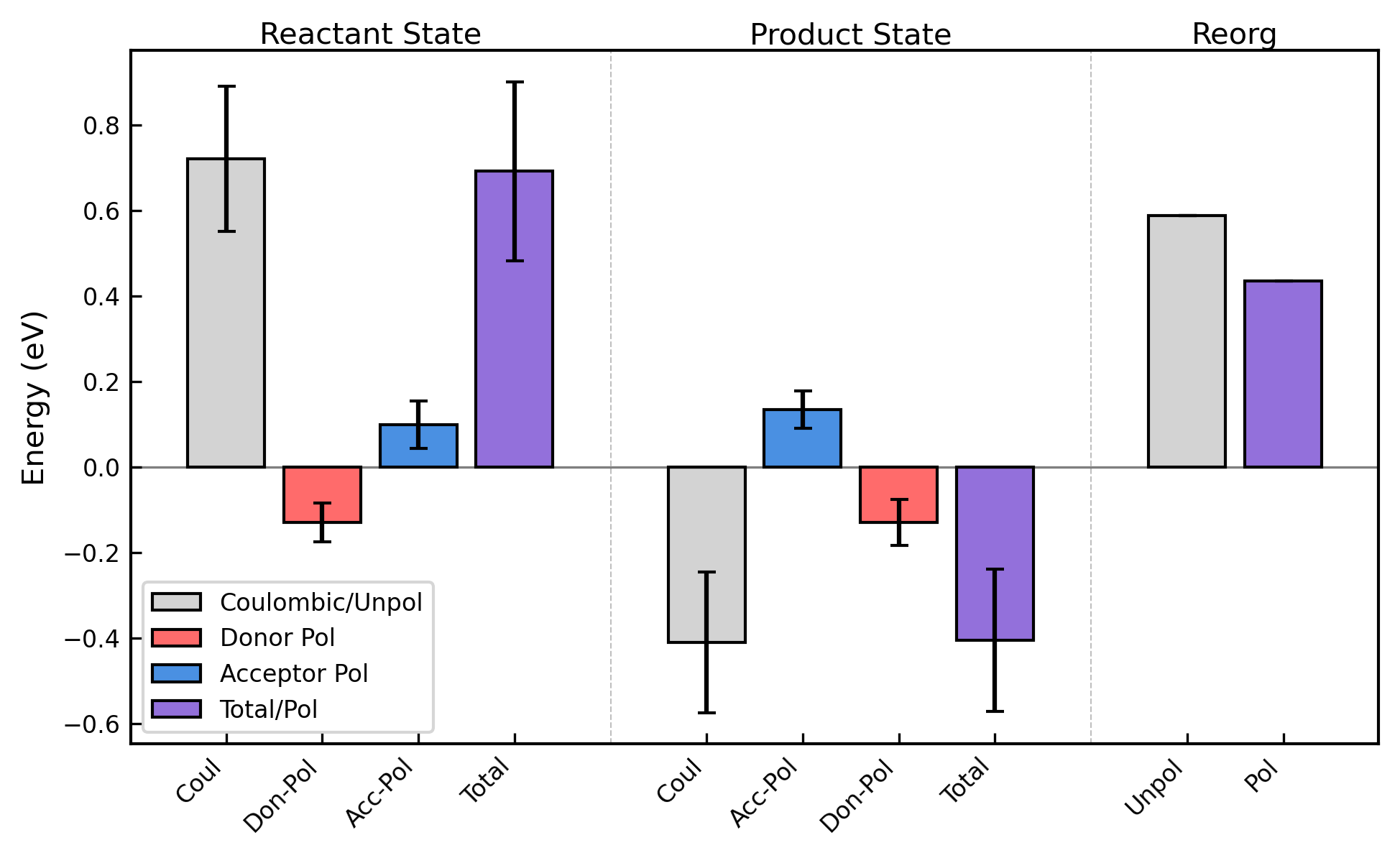

**Supplementary Figure 23.** Contributions to the vertical energy gap for the Heme 3 → 4 (forward) or 4 → 3 (reverse) electron transfer in OmcS from Coulombic interactions, as well as donor (Don) and acceptor (Acc) polarization (Pol). The effect of considering the polarization contributions on the reorganization energy is also shown.

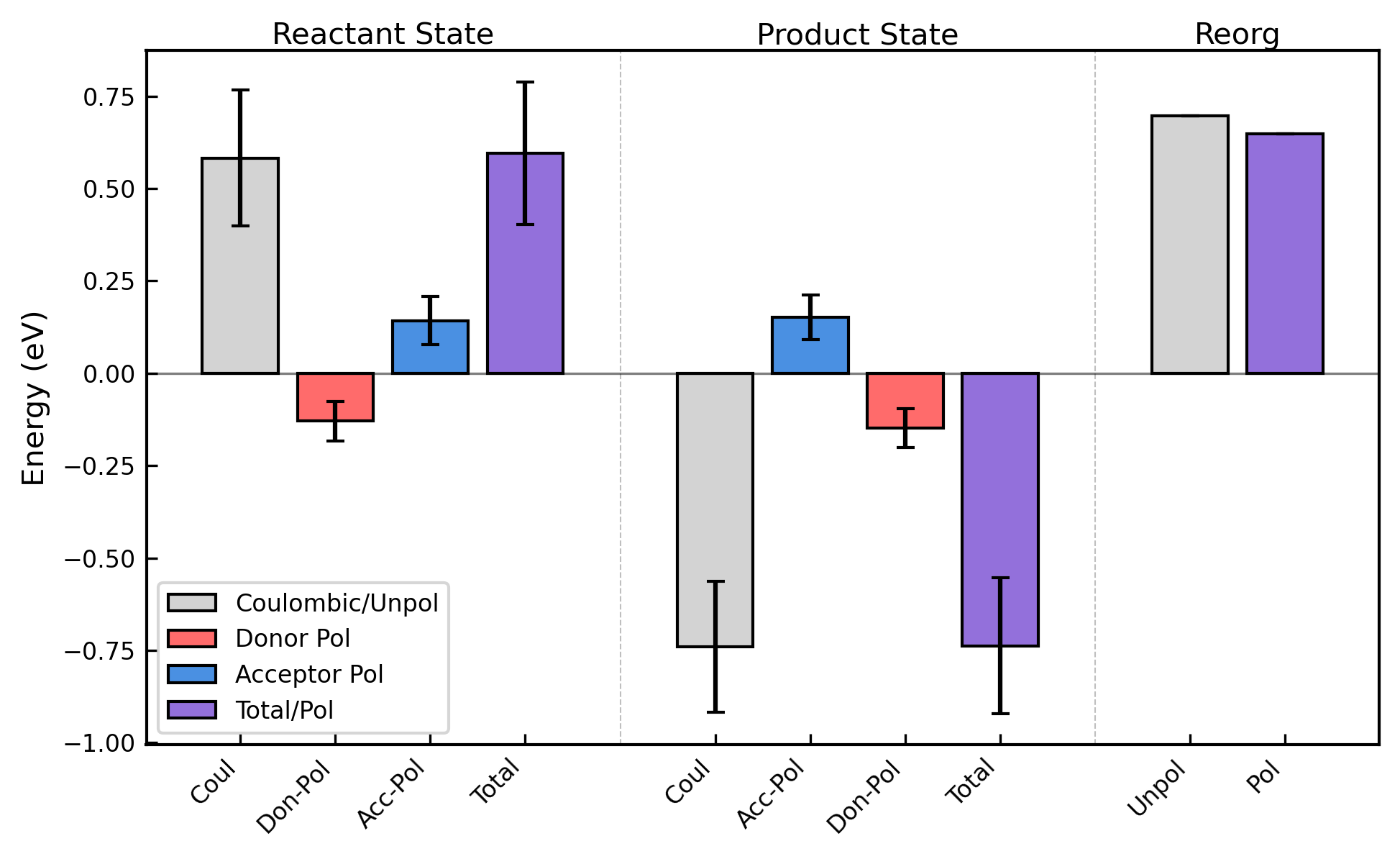

**Supplementary Figure 24.** Contributions to the vertical energy gap for the Heme 4 → 5 (forward) or 5 → 4 (reverse) electron transfer in OmcS from Coulombic interactions, as well as donor (Don) and acceptor (Acc) polarization (Pol). The effect of considering the polarization contributions on the reorganization energy is also shown.

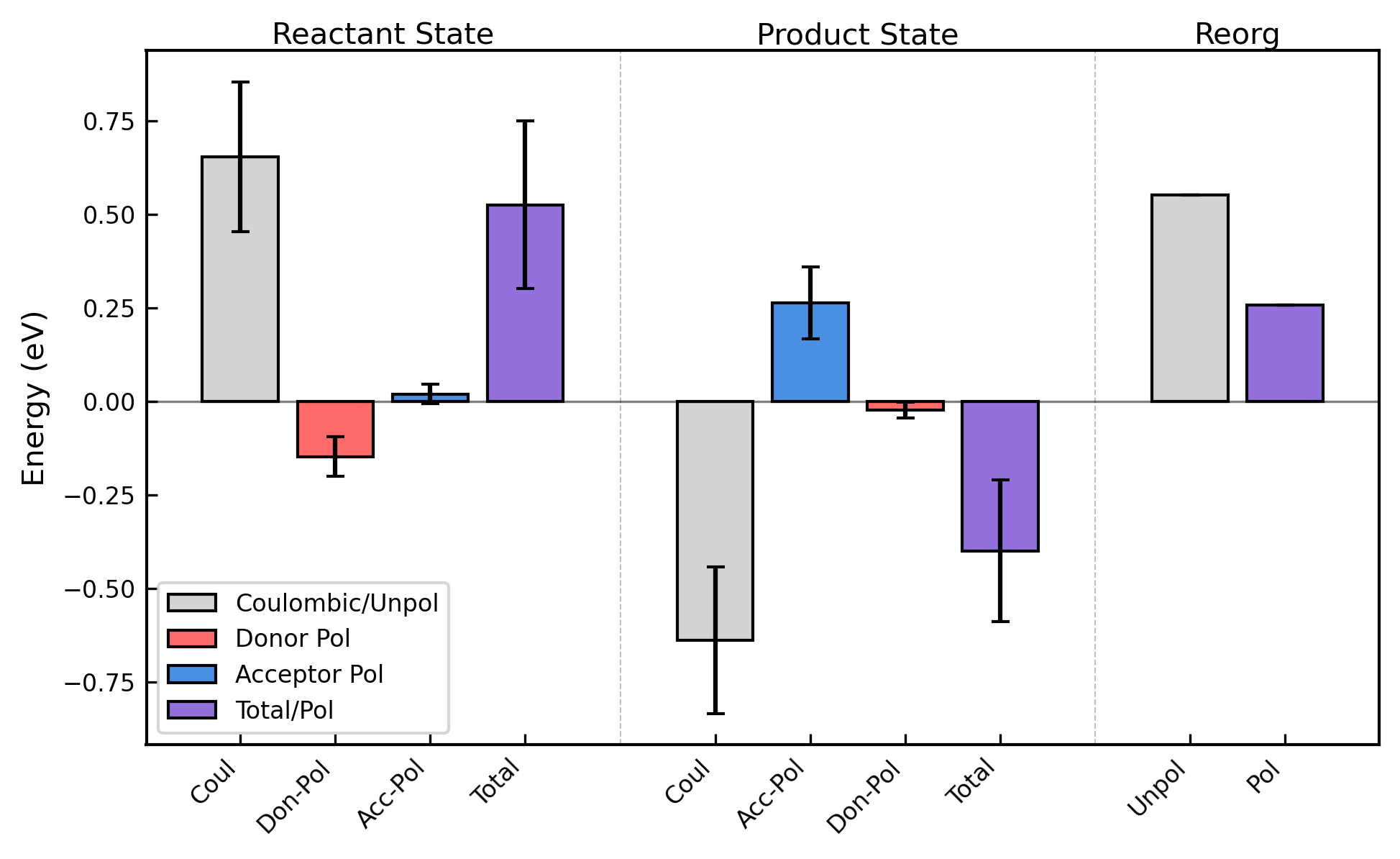

**Supplementary Figure 25.** Contributions to the vertical energy gap for the Heme 5 → 6 (forward) or 6 → 5 (reverse) electron transfer in OmcS from Coulombic interactions, as well as donor (Don) and acceptor (Acc) polarization (Pol). The effect of considering the polarization contributions on the reorganization energy is also shown.

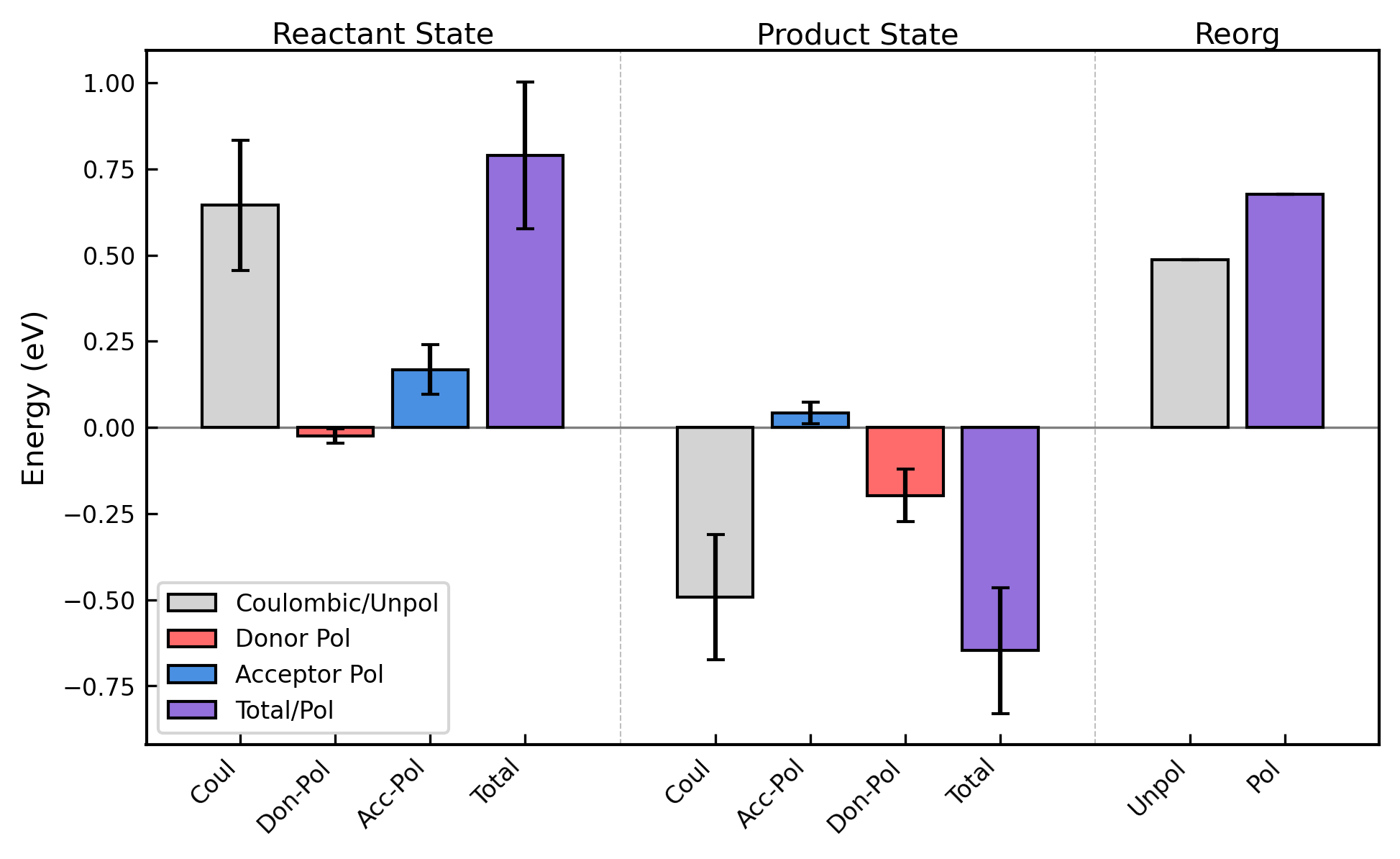

**Supplementary Figure 26.** Contributions to the vertical energy gap for the Heme 6 → 1’ (forward) or 1’ → 6 (reverse) electron transfer in OmcS from Coulombic interactions, as well as donor (Don) and acceptor (Acc) polarization (Pol). The effect of considering the polarization contributions on the reorganization energy is also shown.

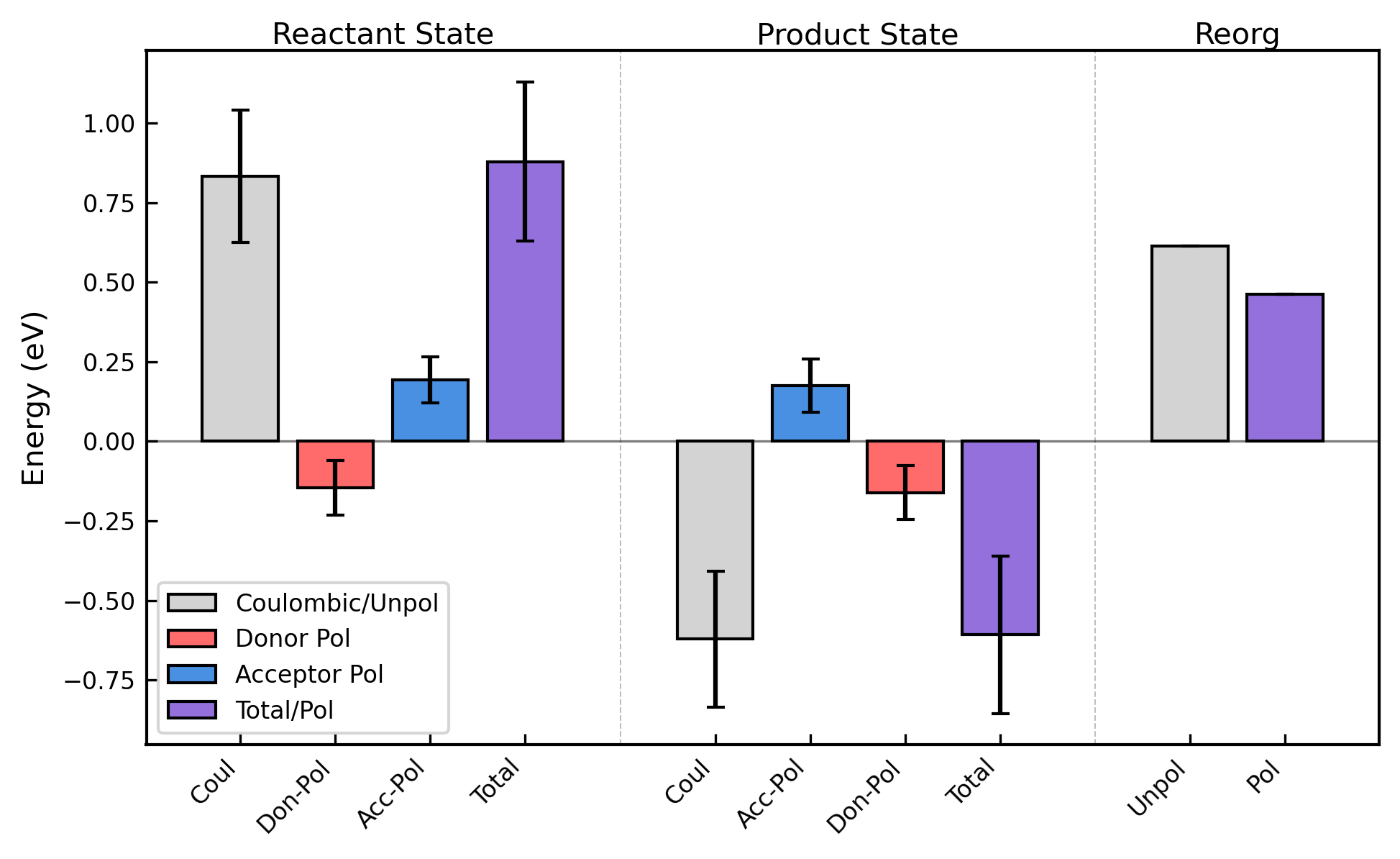

**Supplementary Figure 27.** Contributions to the vertical energy gap for the Heme 1 → 2 (forward) or 2 → 1 (reverse) electron transfer in OmcZ from Coulombic interactions, as well as donor (Don) and acceptor (Acc) polarization (Pol). The effect of considering the polarization contributions on the reorganization energy is also shown.

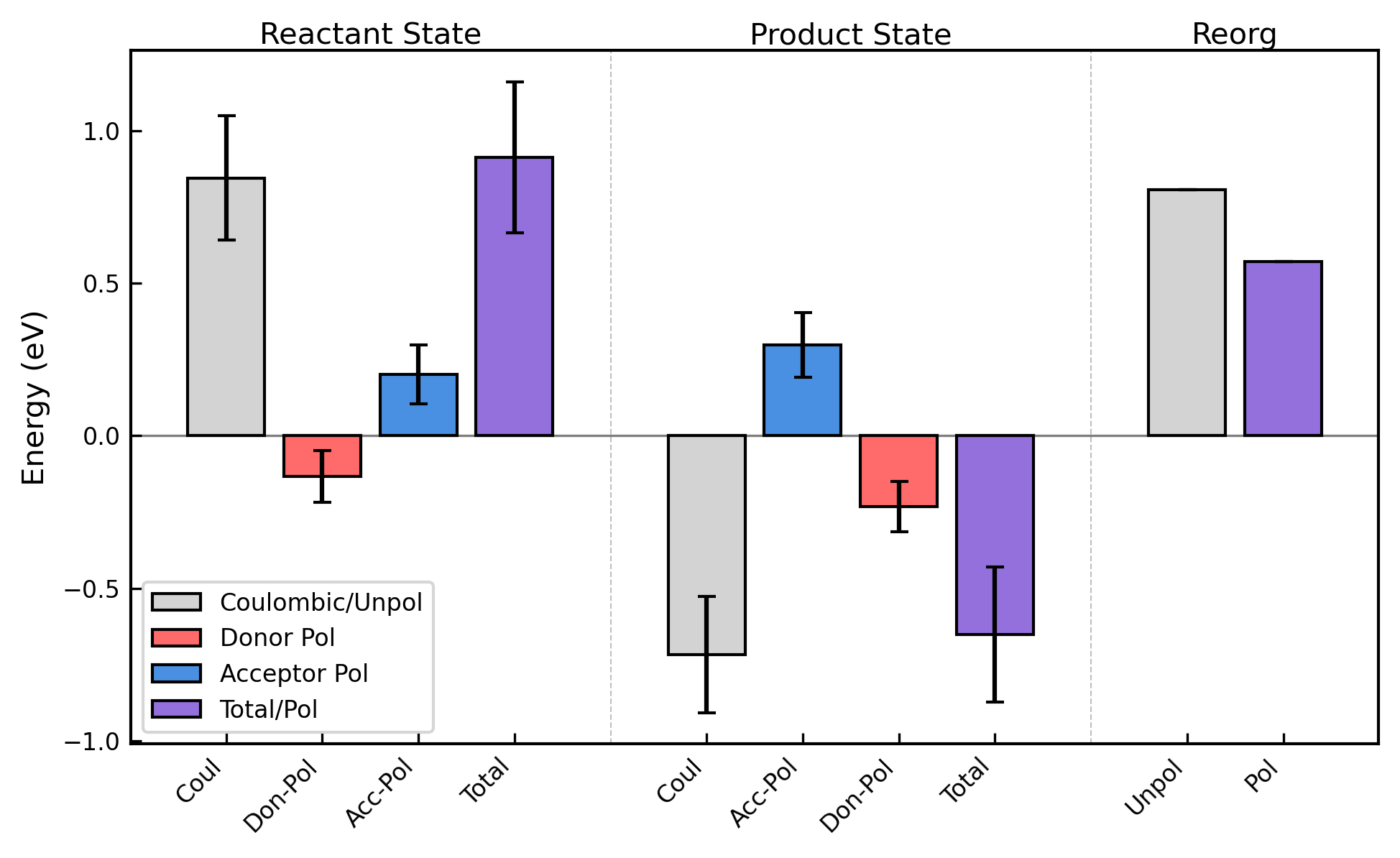

**Supplementary Figure 28.** Contributions to the vertical energy gap for the Heme 2 → 3 (forward) or 3 → 2 (reverse) electron transfer in OmcZ from Coulombic interactions, as well as donor (Don) and acceptor (Acc) polarization (Pol). The effect of considering the polarization contributions on the reorganization energy is also shown.

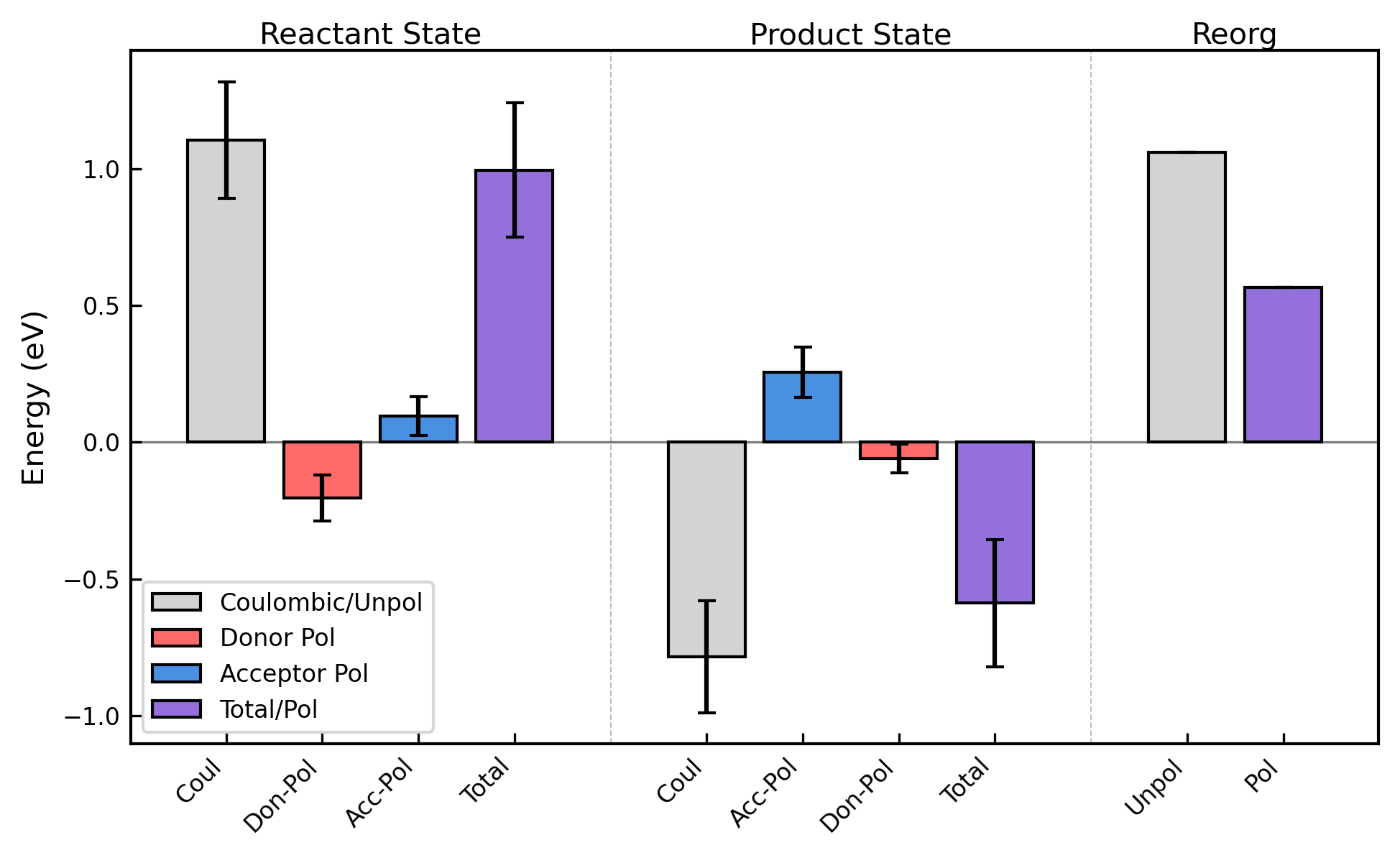

**Supplementary Figure 29.** Contributions to the vertical energy gap for the Heme 3 → 4 (forward) or 4 → 3 (reverse) electron transfer in OmcZ from Coulombic interactions, as well as donor (Don) and acceptor (Acc) polarization (Pol). The effect of considering the polarization contributions on the reorganization energy is also shown.

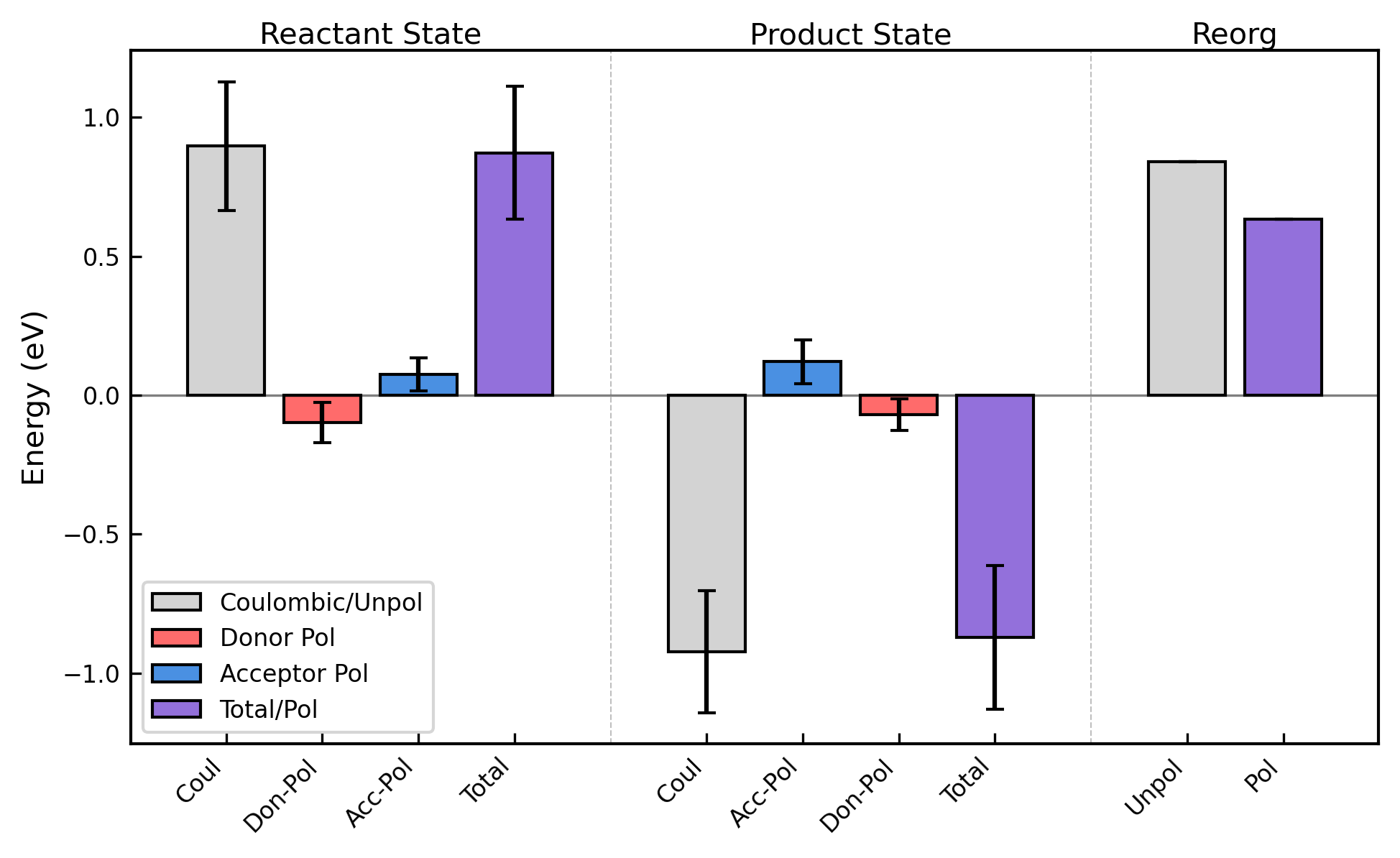

**Supplementary Figure 30.** Contributions to the vertical energy gap for the Heme 4 → 5 (forward) or 5 → 4 (reverse) electron transfer in OmcZ from Coulombic interactions, as well as donor (Don) and acceptor (Acc) polarization (Pol). The effect of considering the polarization contributions on the reorganization energy is also shown.

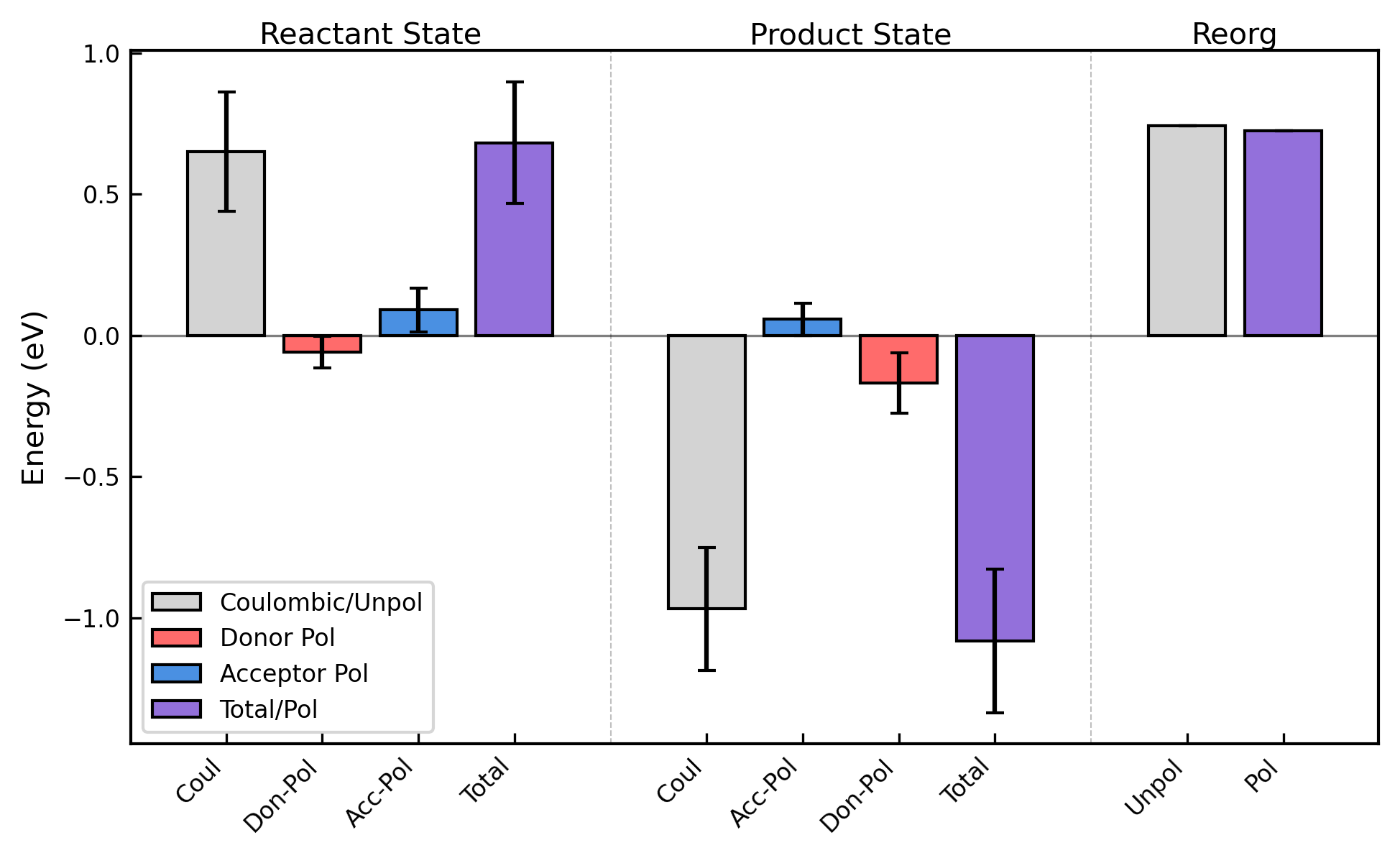

**Supplementary Figure 31.** Contributions to the vertical energy gap for the Heme 5 → 6 (forward) or 6 → 5 (reverse) electron transfer in OmcZ from Coulombic interactions, as well as donor (Don) and acceptor (Acc) polarization (Pol). The effect of considering the polarization contributions on the reorganization energy is also shown.

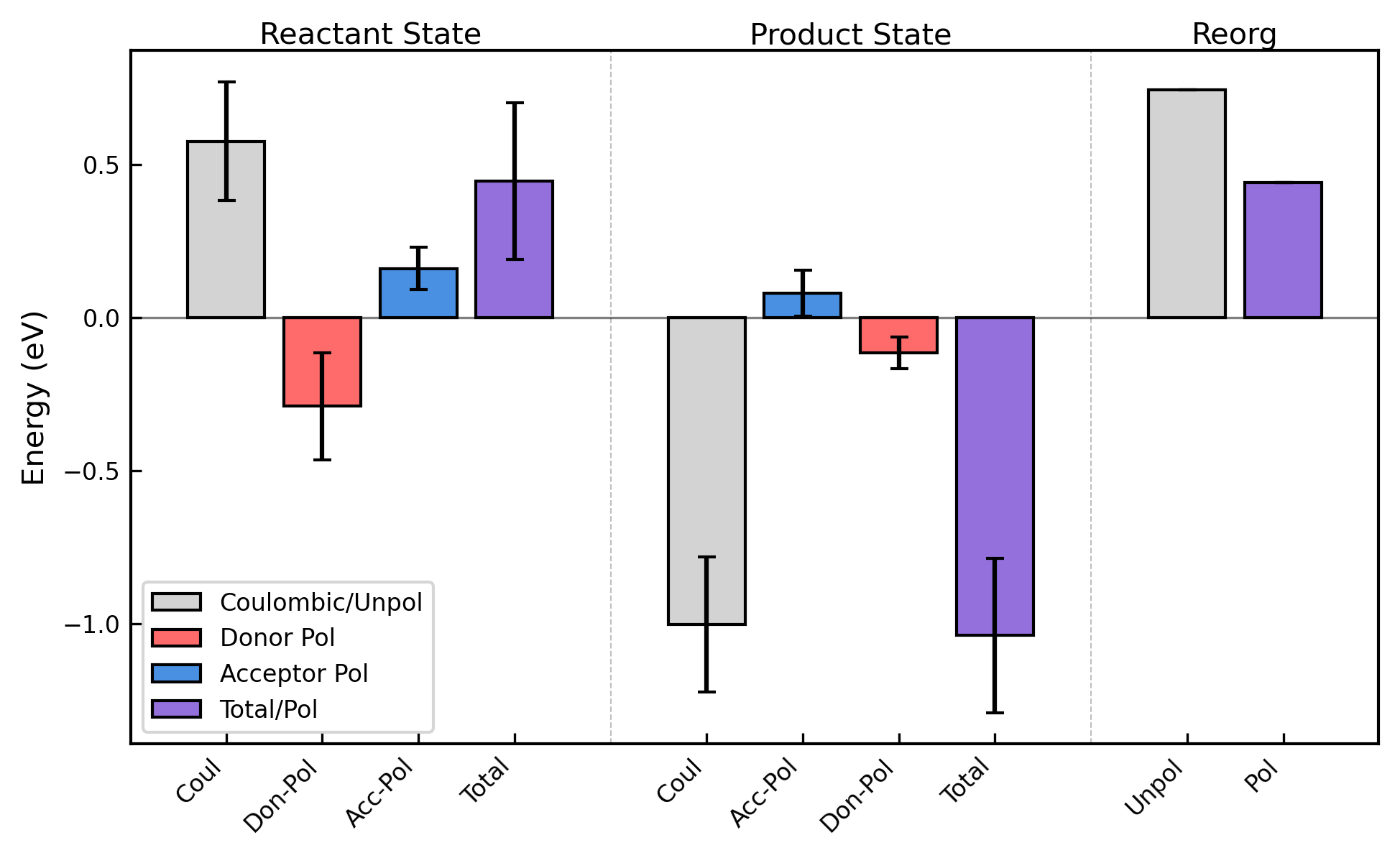

**Supplementary Figure 32.** Contributions to the vertical energy gap for the Heme 6 → 7 (forward) or 7 → 6 (reverse) electron transfer in OmcZ from Coulombic interactions, as well as donor (Don) and acceptor (Acc) polarization (Pol). The effect of considering the polarization contributions on the reorganization energy is also shown.

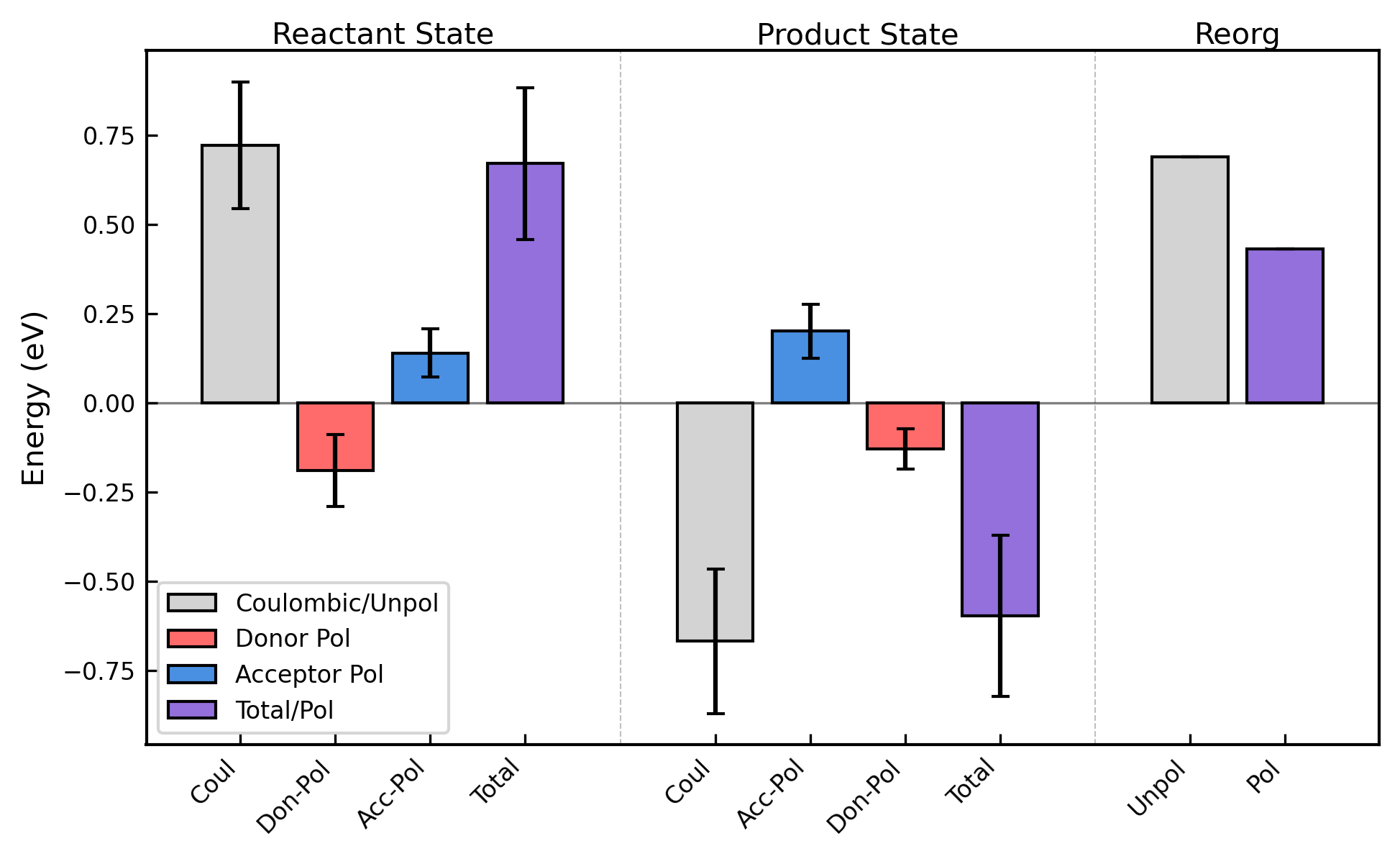

**Supplementary Figure 33.** Contributions to the vertical energy gap for the Heme 7 → 1’ (forward) or 1’ → 7 (reverse) electron transfer in OmcZ from Coulombic interactions, as well as donor (Don) and acceptor (Acc) polarization (Pol). The effect of considering the polarization contributions on the reorganization energy is also shown.

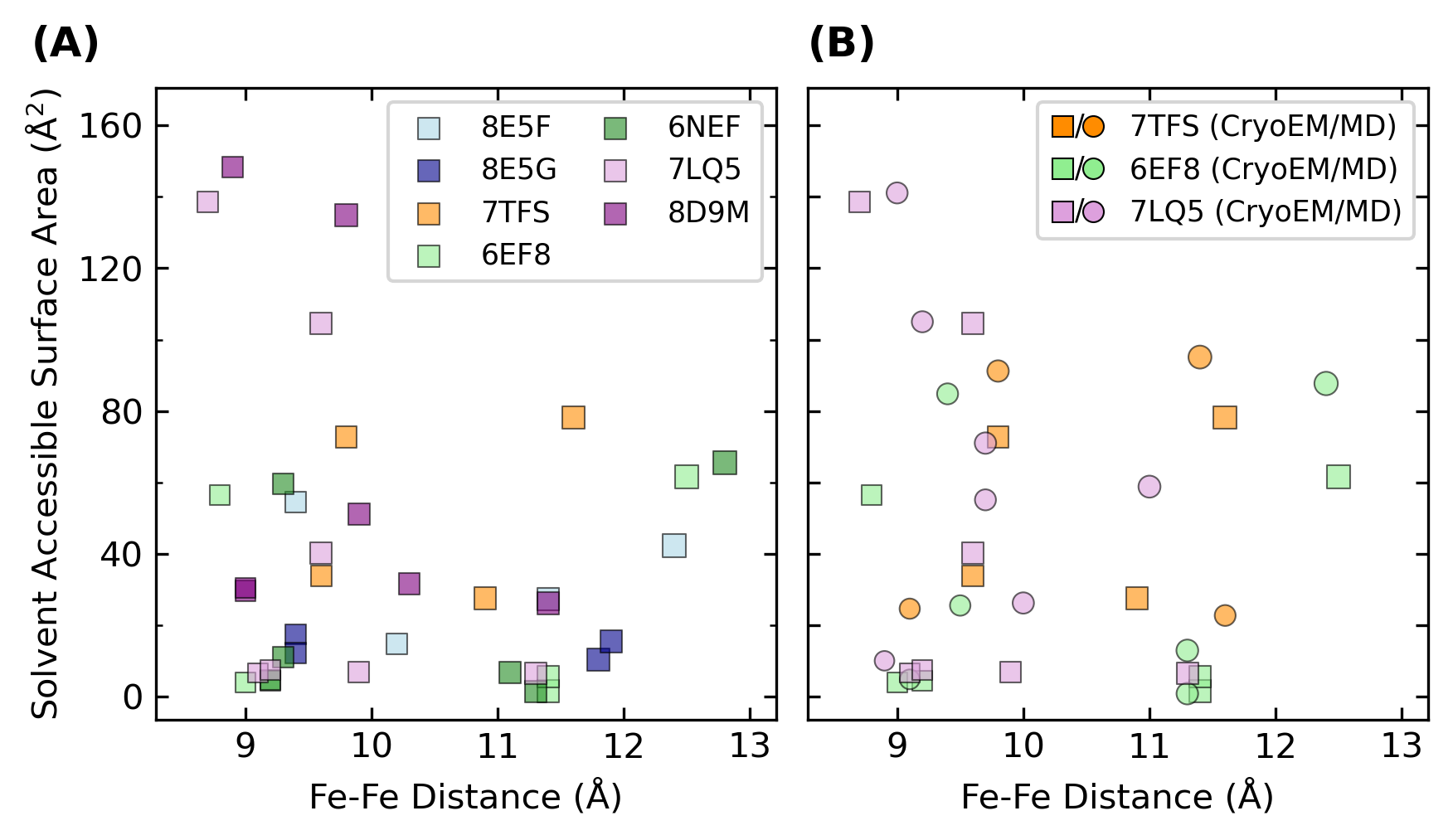

**Supplementary Figure 34.** Plots of the total donor+acceptor solvent accessible surface area versus the Fe-to-Fe distance, with the marker sizes proportional to the outer-sphere reorganization energy computed by Eq. 4 in the main text. Panel A shows the data for all filaments based on their CryoEM structural models. Panel B shows the data for three of the filaments for which molecular dynamics simulations were previously reported. The proteins are referenced by their PDB codes in the figure.

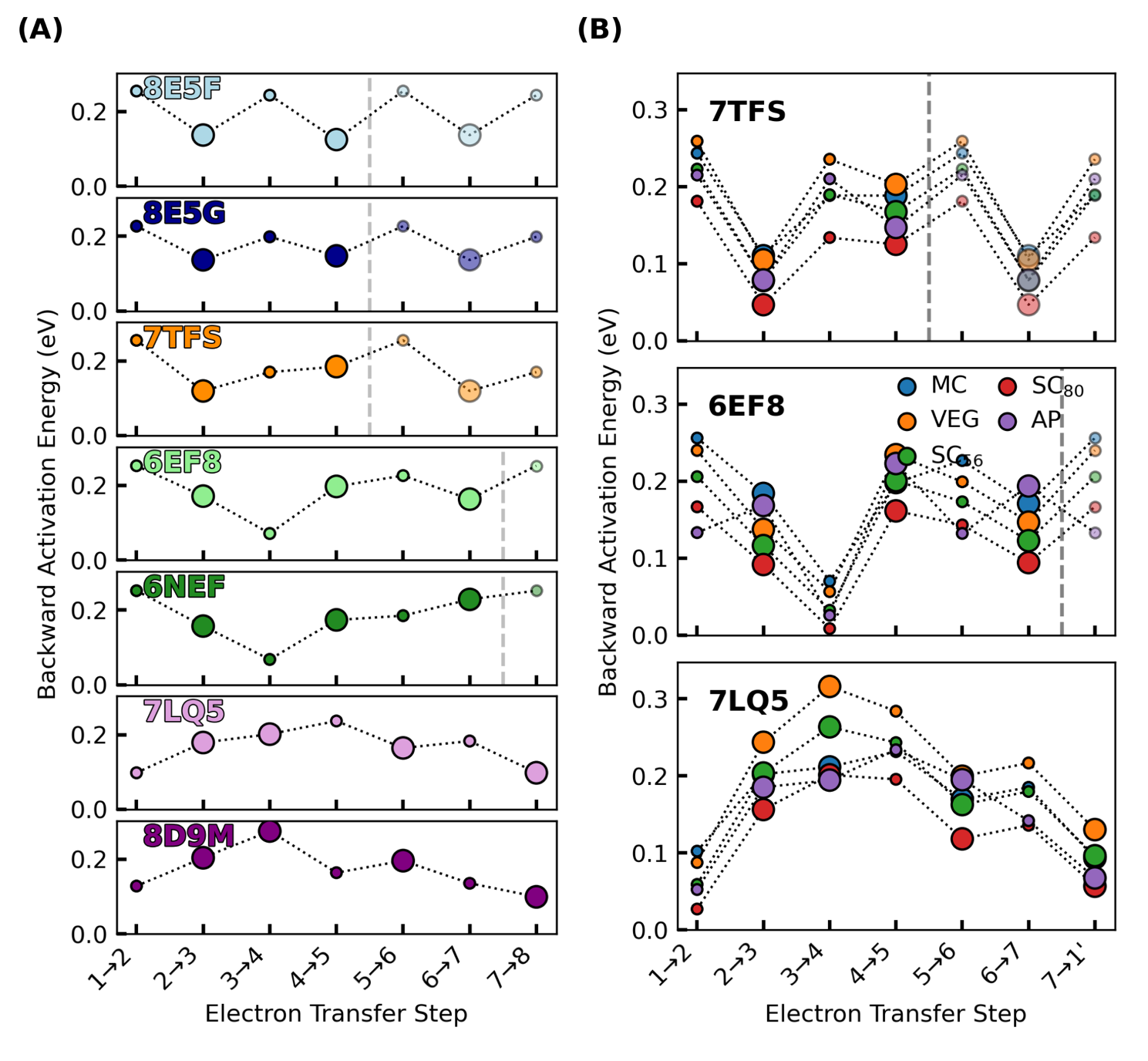

**Supplementary Figure 35.** Activation energies for electron transfers in the known cytochrome ‘nanowires’ are always ≤0.3 eV, regardless of if or how electronic polarization is included in the estimate of outer-sphere reorganization energy. Panel **(A)** shows the cytochrome ‘nanowires’ by PDB accession code: 8E5F, 8E5G, 7TFS, 6EF8/6NEF, 7LQ5/8DoM = A3MW92, F2KMU8, OmcE, OmcS, and OmcZ. Panel **(B)** shows the variation in activation energy for three of these filaments for which the outer-sphere reorganization energy was computed by various methods discussed in the main text. The size of the markers in both panels is proportional to the electronic coupling. Activation energies for the backward reactions are shown; main text Figure 11 shows the activation energies for the forward reactions. All data is presented in Supplementary Table 8 and 9.

### 2. Supplementary Tables

**Supplementary Table 1.** Iso- and anisotropic polarizabilities $\left( \alpha\right)$, and the anisotropy parameter $\left( \kappa=\frac{\alpha_{\mathrm{aniso}}}{3*\alpha_{\mathrm{iso}}} \right)$ computed in vacuum for His-Met and His-His ligated *c*-type heme models with various approximate density functionals and basis sets. All calculations were performed using Gaussian 16, Revision A.03. A common vacuum optimized geometry in a given redox state was used for all polarizability calculations.

| Model Chemistry | $\alpha_{iso,red}$ | $\alpha_{iso,ox}$ | $\Delta\alpha_{\mathrm{iso}}$ | $\alpha_{aniso,red}$ | $\alpha_{aniso,ox}$ | $\Delta\alpha_{\mathrm{aniso}}$ | $\kappa_{\mathrm{red}}$ | $\kappa_{\mathrm{ox}}$ |
| --- | --- | --- | --- | --- | --- | --- | --- | --- |
| His-His Ligated Heme | | | | | | | | |
| CAM-B3LYP/BS1 | 86.55 | 86.94 | 0.39 | 39.80 | 39.31 | -0.49 | 0.15 | 0.15 |
| B3LYP/BS1 | 88.84 | 89.62 | 0.78 | 40.30 | 40.28 | -0.02 | 0.15 | 0.15 |
| BLYP/BS1 | 91.92 | 98.46 | 6.54 | 39.37 | 49.25 | 9.88 | 0.14 | 0.17 |
| PBE/BS1 | 92.12 | 98.09 | 5.97 | 39.10 | 48.26 | 9.16 | 0.14 | 0.16 |
| BLYP/6-31G(d) | 92.25 | 96.24 | 3.99 | 38.29 | 46.46 | 8.17 | 0.14 | 0.16 |
| BLYP/6-31+G(d) | 102.60 | 107.53 | 4.93 | 39.59 | 50.48 | 10.89 | 0.13 | 0.16 |
| BLYP/6-311+G(d) | 102.99 | 108.45 | 5.46 | 39.59 | 51.08 | 11.49 | 0.13 | 0.16 |
| BLYP/6-311+G(d,p) | 103.67 | 109.07 | 5.40 | 39.60 | 51.15 | 11.55 | 0.13 | 0.16 |
| BLYP/6-311+G(2d,p) | 104.67 | 109.89 | 5.22 | 39.69 | 51.12 | 11.43 | 0.13 | 0.16 |
| His-Met Ligated Heme | | | | | | | | |
| CAM-B3LYP/BS1 | 85.76 | 87.03 | 1.27 | 43.30 | 40.97 | -2.33 | 0.17 | 0.16 |
| B3LYP/BS1 | 87.99 | 89.64 | 1.65 | 44.01 | 42.74 | -1.27 | 0.17 | 0.16 |
| BLYP/BS1 | 90.99 | 99.76 | 8.77 | 43.29 | 54.62 | 11.33 | 0.16 | 0.18 |
| PBE/BS1 | 91.26 | 99.23 | 7.97 | 42.91 | 53.24 | 10.33 | 0.16 | 0.18 |
| BLYP/6-31G(d) | 91.16 | 96.76 | 5.60 | 42.51 | 50.89 | 8.38 | 0.16 | 0.18 |
| BLYP/6-31+G(d) | 101.38 | 108.62 | 7.24 | 43.59 | 56.02 | 12.43 | 0.14 | 0.17 |
| BLYP/6-311+G(d) | 101.85 | 109.68 | 7.83 | 43.64 | 56.76 | 13.12 | 0.14 | 0.17 |
| BLYP/6-311+G(d,p) | 102.51 | 110.24 | 7.73 | 43.74 | 56.79 | 13.05 | 0.14 | 0.17 |
| BLYP/6-311+G(2d,p) | 103.50 | 110.95 | 7.45 | 43.87 | 56.64 | 12.77 | 0.14 | 0.17 |

**Supplementary Table 2.** Iso- and anisotropic polarizabilities $\left( \alpha\right)$ computed for the His-His and His-Met ligated hemes of cytochrome *c* and OmcS, computed in vacuum or within the protein-water matrix. The calculations were performed over 200 (cytochrome *c*) or 250 (OmcS) configurations from either the reduced or oxidized ensemble (Ensb.) from classical molecular dynamics for the heme being analyzed. The polarizability was computed using the BLYP function with either the BS1 basis set (LANL2DZ for Fe and 6-31G(d) for H, C, N, and S) (cytochrome *c*) or the 6-31G(d) basis set for all atoms (OmcS). The latter basis set underestimates the oxidized-reduced polarizability difference predicted by the former basis set by 10-30% (Supplementary Table S1). All polarizability calculations were performed with Gaussian 16, Revision A.03 and analyzed with MultiWFN.

| Structure | $\alpha_{iso,red}$ | $\alpha_{iso,ox}$ | $\Delta\alpha_{\mathrm{iso}}$ | $\alpha_{aniso,red}$ | $\alpha_{aniso,ox}$ | $\Delta\alpha_{\mathrm{aniso}}$ |
| --- | --- | --- | --- | --- | --- | --- |
| Cyt. *c* (Red. Ensb., vac.) | 88.8±0.5 | 96.4±2.5 | 7.6±2.6 | 41.4±1.3 | 49.9±4.0 | 8.5±4.0 |
| Cyt. *c* (Ox. Ensb. Vac.) | 88.7±0.6 | 95.9±2.2 | 7.3±2.3 | 40.9±1.1 | 48.9±3.2 | 8.8±3.4 |
| Cyt. *c* (Red. Ensb., Env.) | 88.5±0.6 | 94.6±2.4 | 6.1±2.5 | 41.6±1.1 | 47.8±3.8 | 6.2±4.0 |
| Cyt. *c* (Ox. Ensb. Env.) | 88.8±0.6 | 93.1±2.0 | 4.3±2.1 | 41.3±1.2 | 44.7±3.0 | 3.4±3.2 |
| OmcS #1 (Red. Ensb. Env.) | 90.3±0.7 | 91.7±0.8 | 1.4±1.8 | 38.1±1.2 | 40.5±1.3 | 2.4±0.8 |
| OmcS #2 (Red. Ensb. Env.) | 90.3±0.6 | 92.1±1.0 | 1.8±2.0 | 39.1±1.2 | 43.0±1.6 | 4.0±1.0 |
| OmcS #3 (Red. Ensb. Env.) | 90.0±0.6 | 91.3±0.8 | 1.3±1.8 | 37.9±1.2 | 39.2±1.4 | 1.3±0.8 |
| OmcS #4 (Red. Ensb. Env.) | 90.4±0.6 | 92.3±0.9 | 1.9±2.0 | 39.2±1.2 | 42.0±1.6 | 2.8±0.9 |
| OmcS #5 (Red. Ensb. Env.) | 90.3±0.6 | 91.6±0.8 | 1.3±2.4 | 38.5±1.6 | 41.5±1.8 | 3.0±0.8 |
| OmcS #6 (Red. Ensb. Env.) | 90.3±0.7 | 91.9±0.9 | 1.6±2.1 | 38.1±1.4 | 41.0±1.6 | 2.9±0.9 |

**Supplementary Table 3.** Outer-sphere reorganization energies $\left( \lambda_{\mathrm{out}} \right)$ in eV were assessed via Eqs. 5–10 in the main text by computing the Coulombic vertical energy gaps, the redox-linked change in the polarizability tensor of the heme group, the electric field acting at the heme-Fe centers, and the polarization energies of the donor and the acceptor.

|  | Unpolarized | | | | | Polarized | | | | |
| --- | --- | --- | --- | --- | --- | --- | --- | --- | --- | --- |
| Pair | $\lambda^{\mathrm{st}}$ | $\lambda^{var,f}$ | $\lambda^{var,b}$ | $\lambda^{rxn}$ | $\chi_{g}$ | $\lambda^{\mathrm{st}}$ | $\lambda^{var,f}$ | $\lambda^{var,b}$ | $\lambda^{rxn}$ | $\chi_{g}$ |
| OmcE | | | | | | | | | | |
| 1↔2 | 0.772 | 0.824 | 0.767 | 0.749 | 1.031 | 0.714 | 1.021 | 0.779 | 0.566 | 1.261 |
| 2↔3 | 0.628 | 0.655 | 0.796 | 0.544 | 1.155 | 0.616 | 0.794 | 0.939 | 0.438 | 1.406 |
| 3↔4 | 0.936 | 0.922 | 0.973 | 0.925 | 1.012 | 1.053 | 1.315 | 1.381 | 0.823 | 1.280 |
| 4↔1’ | 0.701 | 0.694 | 0.686 | 0.713 | 0.984 | 0.721 | 1.060 | 1.072 | 0.488 | 1.477 |
| OmcS | | | | | | | | | | |
| 1↔2 | 0.813 | 0.912 | 0.974 | 0.701 | 1.160 | 0.552 | 1.147 | 1.444 | 0.236 | 2.345 |
| 2↔3 | 0.470 | 0.512 | 0.517 | 0.429 | 1.095 | 0.572 | 0.577 | 0.605 | 0.554 | 1.033 |
| 3↔4 | 0.566 | 0.562 | 0.525 | 0.589 | 0.960 | 0.549 | 0.850 | 0.532 | 0.436 | 1.259 |
| 4↔5 | 0.662 | 0.650 | 0.607 | 0.696 | 0.950 | 0.666 | 0.720 | 0.649 | 0.649 | 1.027 |
| 5↔6 | 0.646 | 0.776 | 0.739 | 0.551 | 1.173 | 0.462 | 0.970 | 0.696 | 0.257 | 1.801 |
| 6↔1’ | 0.569 | 0.690 | 0.638 | 0.487 | 1.167 | 0.718 | 0.880 | 0.645 | 0.676 | 1.062 |
| OmcZ | | | | | | | | | | |
| 1↔2 | 0.727 | 0.841 | 0.884 | 0.613 | 1.186 | 0.744 | 1.214 | 1.183 | 0.461 | 1.612 |
| 2↔3 | 0.782 | 0.811 | 0.701 | 0.808 | 0.967 | 0.782 | 1.191 | 0.950 | 0.572 | 1.369 |
| 3↔4 | 0.944 | 0.879 | 0.805 | 1.059 | 0.892 | 0.792 | 1.164 | 1.053 | 0.565 | 1.401 |
| 4↔5 | 0.910 | 1.036 | 0.932 | 0.841 | 1.081 | 0.872 | 1.108 | 1.291 | 0.634 | 1.375 |
| 5↔6 | 0.810 | 0.864 | 0.907 | 0.741 | 1.093 | 0.881 | 0.898 | 1.251 | 0.723 | 1.219 |
| 6↔7 | 0.789 | 0.729 | 0.943 | 0.744 | 1.060 | 0.741 | 1.263 | 1.231 | 0.441 | 1.681 |
| 7↔1’ | 0.695 | 0.608 | 0.794 | 0.689 | 1.009 | 0.633 | 0.875 | 0.985 | 0.431 | 1.468 |

**Supplementary Table 4.** Outer-sphere reorganization energies $\left( \lambda_{\mathrm{out}} \right)$ in eV were assessed via Eqs. 3 and 4 in the main text by computing the solvent accessible surface area (Å^2^) for the donor and acceptor and Fe-to-Fe distance (Å) from the CryoEM structures indicated by PDB code.

| PDB ID | D → A  Residue ID | Donor SASA | Acceptor  SASA | Total SASA | $\epsilon_{s}$ | Fe-Fe  Distance | $\lambda_{\mathrm{out}}$ |
| --- | --- | --- | --- | --- | --- | --- | --- |
| 7TFS | 651→654 | 63.6 | 14.6 | 78.2 | 6.4 | 11.6 | 0.733 |
| 7TFS | 654→657 | 14.6 | 19.2 | 33.8 | 5.7 | 9.6 | 0.603 |
| 7TFS | 657→660 | 19.2 | 8.3 | 27.5 | 5.6 | 10.9 | 0.663 |
| 7TFS | 660→639 | 8.3 | 64.5 | 72.8 | 6.3 | 9.8 | 0.640 |
| 8E5F | 933→936 | 1.3 | 41.0 | 42.3 | 5.9 | 12.4 | 0.734 |
| 8E5F | 936→939 | 41.0 | 13.4 | 54.4 | 6.1 | 9.4 | 0.605 |
| 8E5F | 939→942 | 13.4 | 14.0 | 27.3 | 5.6 | 11.4 | 0.683 |
| 8E5F | 942→921 | 14.0 | 0.8 | 14.8 | 5.4 | 10.2 | 0.618 |
| 8E5G | 1125→1128 | 2.2 | 8.1 | 10.3 | 5.3 | 11.8 | 0.682 |
| 8E5G | 1128→1131 | 8.1 | 9.3 | 17.4 | 5.5 | 9.4 | 0.579 |
| 8E5G | 1131→1134 | 9.3 | 6.2 | 15.5 | 5.4 | 11.9 | 0.691 |
| 8E5G | 1134→1137 | 6.2 | 6.0 | 12.2 | 5.4 | 9.4 | 0.573 |
| 6EF8 | 1272→1266 | 5.2 | 56.4 | 61.6 | 6.2 | 12.5 | 0.755 |
| 6EF8 | 1266→1269 | 56.4 | 0.0 | 56.4 | 6.1 | 8.8 | 0.567 |
| 6EF8 | 1269→1263 | 0.0 | 1.5 | 1.5 | 5.2 | 11.4 | 0.657 |
| 6EF8 | 1263→1260 | 1.5 | 2.5 | 4.0 | 5.2 | 9.0 | 0.543 |
| 6EF8 | 1260→1275 | 2.5 | 3.0 | 5.6 | 5.3 | 11.4 | 0.663 |
| 6EF8 | 1275→1254 | 3.0 | 1.4 | 4.4 | 5.3 | 9.2 | 0.554 |
| 6NEF | 1260→1263 | 6.0 | 59.5 | 65.5 | 6.2 | 12.8 | 0.770 |
| 6NEF | 1263→1266 | 59.5 | 0.0 | 59.5 | 6.1 | 9.3 | 0.601 |
| 6NEF | 1266→1269 | 0.0 | 1.3 | 1.3 | 5.2 | 11.3 | 0.654 |
| 6NEF | 1269→1272 | 1.3 | 3.4 | 4.7 | 5.3 | 9.2 | 0.553 |
| 6NEF | 1272→1275 | 3.4 | 3.4 | 6.8 | 5.3 | 11.1 | 0.652 |
| 6NEF | 1275→1242 | 3.4 | 7.5 | 11.0 | 5.4 | 9.3 | 0.565 |
| 7LQ5 | 562→553 | 2.3 | 4.2 | 6.4 | 5.3 | 11.3 | 0.658 |
| 7LQ5 | 553→559 | 4.2 | 2.5 | 6.6 | 5.3 | 9.1 | 0.547 |
| 7LQ5 | 559→550 | 2.5 | 4.5 | 7.0 | 5.3 | 9.9 | 0.594 |
| 7LQ5 | 550→541 | 4.5 | 100.0 | 104.5 | 6.9 | 9.6 | 0.646 |
| 7LQ5 | 541→547 | 100.0 | 38.5 | 138.5 | 7.4 | 8.7 | 0.601 |
| 7LQ5 | 547→544 | 38.5 | 1.6 | 40.2 | 5.8 | 9.6 | 0.609 |
| 7LQ5 | 544→844 | 1.6 | 5.7 | 7.4 | 5.3 | 9.2 | 0.558 |
| 8D9M | 562→559 | 10.1 | 16.0 | 26.1 | 5.6 | 11.4 | 0.683 |
| 8D9M | 559→553 | 16.0 | 13.5 | 29.6 | 5.7 | 9.0 | 0.563 |
| 8D9M | 553→550 | 13.5 | 18.1 | 31.6 | 5.7 | 10.3 | 0.640 |
| 8D9M | 550→547 | 18.1 | 116.8 | 134.8 | 7.3 | 9.9 | 0.681 |
| 8D9M | 547→544 | 116.8 | 31.7 | 148.4 | 7.6 | 8.9 | 0.622 |
| 8D9M | 544→541 | 31.7 | 19.4 | 51.1 | 6.0 | 9.9 | 0.630 |
| 8D9M | 541→844 | 19.4 | 11.1 | 30.5 | 5.7 | 9.0 | 0.564 |

Supplementary Table 5. Outer-sphere reorganization energies $\left( \lambda_{\mathrm{out}} \right)$ in eV were assessed via Eqs. 3 and 4 in the main text by computing the solvent accessible surface area (Å^2^) for the donor and acceptor and Fe-to-Fe distance (Å) from MD simulations previously performed for three of the cytochrome filaments ^1^.

| Omc  type | PDB ID | D → A  Residue ID | Donor SASA | Acceptor  SASA | Total SASA | $\epsilon_{s}$ | Fe-Fe  Distance | $\lambda_{\mathrm{out}}$ |
| --- | --- | --- | --- | --- | --- | --- | --- | --- |
| E | 7TFS | 659→656 | 11.9 | 12.8 | 24.7 | 5.6 | 11.5 | 0.683 |
| E | 7TFS | 656→659 | 14.7 | 6.2 | 20.9 | 5.5 | 11.7 | 0.687 |
| E | 7TFS | 656→653 | 14.7 | 12.6 | 27.4 | 5.6 | 9.2 | 0.575 |
| E | 7TFS | 653→656 | 12.7 | 9.1 | 21.8 | 5.5 | 9.0 | 0.556 |
| E | 7TFS | 653→650 | 12.7 | 86.6 | 99.3 | 6.8 | 11.4 | 0.740 |
| E | 7TFS | 650→653 | 71.9 | 19.1 | 91.0 | 6.6 | 11.3 | 0.731 |
| E | 7TFS | 650→671 | 71.9 | 5.3 | 77.2 | 6.4 | 9.8 | 0.644 |
| E | 7TFS | 671→650 | 22.6 | 82.7 | 105.4 | 6.9 | 9.8 | 0.663 |
| S | 6EF8 | 1280→1274 | 4.5 | 81.4 | 85.8 | 6.6 | 12.3 | 0.768 |
| S | 6EF8 | 1274→1280 | 82.7 | 6.9 | 89.6 | 6.6 | 12.4 | 0.772 |
| S | 6EF8 | 1274→1277 | 82.7 | 0.1 | 82.8 | 6.5 | 9.3 | 0.618 |
| S | 6EF8 | 1277→î1274 | 0.1 | 86.6 | 86.7 | 6.6 | 9.4 | 0.625 |
| S | 6EF8 | 1277→1271 | 0.1 | 0.5 | 0.6 | 5.2 | 11.3 | 0.651 |
| S | 6EF8 | 1271→1277 | 0.8 | 0.2 | 1.0 | 5.2 | 11.2 | 0.648 |
| S | 6EF8 | 1271→1268 | 0.8 | 4.1 | 4.9 | 5.3 | 9.1 | 0.546 |
| S | 6EF8 | 1268→1271 | 4.2 | 0.5 | 4.7 | 5.3 | 9.0 | 0.543 |
| S | 6EF8 | 1268→1283 | 4.2 | 8.9 | 13.1 | 5.4 | 11.3 | 0.665 |
| S | 6EF8 | 1283→1268 | 8.7 | 4.0 | 12.6 | 5.4 | 11.3 | 0.663 |
| S | 6EF8 | 1283→1298 | 8.7 | 13.0 | 21.6 | 5.5 | 9.5 | 0.584 |
| S | 6EF8 | 1298→1283 | 13.5 | 16.0 | 29.5 | 5.7 | 9.6 | 0.601 |
| Z | 7LQ5 | 562→553 | 3.7 | 32.8 | 36.5 | 5.8 | 11.0 | 0.676 |
| Z | 7LQ5 | 553→562 | 71.1 | 10.0 | 81.1 | 6.5 | 11.0 | 0.708 |
| Z | 7LQ5 | 553→559 | 71.1 | 8.3 | 79.4 | 6.5 | 9.7 | 0.641 |
| Z | 7LQ5 | 559→553 | 15.1 | 47.3 | 62.5 | 6.2 | 9.6 | 0.622 |
| Z | 7LQ5 | 559→550 | 15.1 | 24.3 | 39.5 | 5.8 | 10.1 | 0.635 |
| Z | 7LQ5 | 550→559 | 4.6 | 8.5 | 13.0 | 5.4 | 9.9 | 0.601 |
| Z | 7LQ5 | 550→541 | 4.6 | 97.0 | 101.6 | 6.8 | 9.2 | 0.624 |
| Z | 7LQ5 | 541→550 | 98.4 | 10.1 | 108.5 | 6.9 | 9.2 | 0.624 |
| Z | 7LQ5 | 541→547 | 98.4 | 40.0 | 138.5 | 7.4 | 8.9 | 0.621 |
| Z | 7LQ5 | 547→541 | 46.3 | 97.5 | 143.8 | 7.5 | 9.1 | 0.637 |
| Z | 7LQ5 | 547→544 | 46.3 | 1.1 | 47.5 | 5.9 | 9.7 | 0.617 |
| Z | 7LQ5 | 544→547 | 3.7 | 59.1 | 62.8 | 6.2 | 9.7 | 0.628 |
| Z | 7LQ5 | 544→844 | 3.7 | 6.0 | 9.7 | 5.3 | 8.9 | 0.538 |
| Z | 7LQ5 | 844→544 | 8.2 | 2.2 | 10.3 | 5.3 | 8.9 | 0.541 |

**Supplementary Table 6.** Electric field magnitude measured at the heme-Fe centers in the minimized CryoEM structures at a 1.0 Å solvent cutoff

| PDB ID | Residue | Electric Field Magnitude (V/Å)  at Heme-Fe Center |
| --- | --- | --- |
| 7TFS | 651 | 0.520 |
| 7TFS | 654 | 0.360 |
| 7TFS | 657 | 0.436 |
| 7TFS | 660 | 0.612 |
| 7TFS | 639 | 0.223 |
| 8E5F | 933 | 0.443 |
| 8E5F | 936 | 0.665 |
| 8E5F | 939 | 0.257 |
| 8E5F | 942 | 0.180 |
| 8E5F | 921 | 0.249 |
| 8E5G | 1125 | 0.584 |
| 8E5G | 1128 | 0.621 |
| 8E5G | 1131 | 0.307 |
| 8E5G | 1134 | 0.539 |
| 8E5G | 1137 | 0.568 |
| 6EF8 | 1272 | 0.229 |
| 6EF8 | 1266 | 0.099 |
| 6EF8 | 1269 | 0.459 |
| 6EF8 | 1263 | 0.714 |
| 6EF8 | 1260 | 0.437 |
| 6EF8 | 1275 | 0.666 |
| 6EF8 | 1254 | 0.302 |
| 6NEF | 1260 | 0.070 |
| 6NEF | 1263 | 0.255 |
| 6NEF | 1266 | 0.720 |
| 6NEF | 1269 | 0.565 |
| 6NEF | 1272 | 0.400 |
| 6NEF | 1275 | 0.305 |
| 6NEF | 1242 | 0.196 |
| 7LQ5 | 562 | 0.353 |
| 7LQ5 | 553 | 0.521 |
| 7LQ5 | 559 | 0.510 |
| 7LQ5 | 550 | 0.333 |
| 7LQ5 | 541 | 0.329 |
| 7LQ5 | 547 | 0.346 |
| 7LQ5 | 544 | 0.170 |
| 7LQ5 | 844 | 0.194 |
| 8D9M | 562 | 0.411 |
| 8D9M | 559 | 0.418 |
| 8D9M | 553 | 0.440 |
| 8D9M | 550 | 0.297 |
| 8D9M | 547 | 0.382 |
| 8D9M | 544 | 0.285 |
| 8D9M | 541 | 0.190 |
| 8D9M | 844 | 0.520 |

**Supplementary Table 7.** Electric field magnitude (Average ± Standard Deviation) measured at the heme-Fe centers for 1.0 and 29.0 Å solvent cutoffs over molecular dynamics trajectory in which the donor and acceptor were respectively in the reduced and oxidized state.

| Step | Donor | \|E\| (V/Å)  @ 1.0 Å cutoff | \|E\| (VÅ)  @ 29 Å cutoff | Acceptor | \|E\| (V/Å)  @ 1.0 Å cutoff | \|E\| (VÅ)  @ 29 Å cutoff |
| --- | --- | --- | --- | --- | --- | --- |
| OmcE (PDB 7TFS) | | | | | | |
| 1 | 659 | 0.425±0.054 | 0.158±0.036 | 656 | 0.717±0.093 | 0.215±0.052 |
| 2 | 656 | 0.720±0.085 | 0.128±0.050 | 653 | 0.504±0.042 | 0.185±0.043 |
| 3 | 653 | 0.440±0.060 | 0.156±0.043 | 650 | 0.507±0.097 | 0.299±0.056 |
| 4 | 650 | 0.491±0.058 | 0.220±0.058 | 671 | 0.684±0.051 | 0.301±0.046 |
| OmcS (PDB 6EF8) | | | | | | |
| 1 | 1280 | 0.450±0.078 | 0.288±0.047 | 1274 | 0.150±0.043 | 0.131±0.040 |
| 2 | 1274 | 0.181±0.042 | 0.146±0.032 | 1277 | 0.198±0.040 | 0.219±0.034 |
| 3 | 1277 | 0.184±0.040 | 0.207±0.035 | 1271 | 0.403±0.037 | 0.252±0.048 |
| 4 | 1271 | 0.451±0.036 | 0.222±0.041 | 1268 | 0.647±0.052 | 0.207±0.046 |
| 5 | 1268 | 0.594±0.044 | 0.215±0.038 | 1283 | 0.259±0.044 | 0.097±0.036 |
| 6 | 1283 | 0.264±0.040 | 0.091±0.031 | 1298 | 0.381±0.063 | 0.231±0.045 |
| OmcZ (PDB 7LQ5) | | | | | | |
| 1 | 562 | 0.329±0.040 | 0.197±0.037 | 553 | 0.394±0.071 | 0.224±0.041 |
| 2 | 553 | 0.451±0.077 | 0.185±0.045 | 559 | 0.367±0.071 | 0.227±0.052 |
| 3 | 559 | 0.500±0.090 | 0.254±0.045 | 550 | 0.440±0.060 | 0.178±0.053 |
| 4 | 550 | 0.355±0.059 | 0.167±0.052 | 541 | 0.305±0.077 | 0.140±0.053 |
| 5 | 541 | 0.229±0.068 | 0.152±0.049 | 547 | 0.371±0.059 | 0.177±0.060 |
| 6 | 547 | 0.305±0.064 | 0.180±0.052 | 544 | 0.420±0.043 | 0.253±0.046 |
| 7 | 544 | 0.357±0.041 | 0.189±0.035 | 844 | 0.294±0.042 | 0.187±0.047 |

**Supplementary Table 8.** Energetic and kinetic parameters for the structurally characterized cytochrome ‘nanowires’ obtained with the BioDC program. Geom. Is the classification of the donor (D)-acceptor (A) pair as slip-stacked (S) or T-stacked (T). H_da_ (in meV) is the electronic coupling and is assigned a value of 8 or 2 meV for S and T geometries, respectively. $\Delta G^{\circ}$ (in eV) is the reaction free energy obtained by oxidizing hemes in the linear sequence prescribed by the filament topology while including heme-heme interactions. $\lambda$ (in eV) is the outer-sphere reorganization energy estimated by the empirically parameterized Marcus Continuum approach described in the main text. $E_{a,f}$, and $E_{a,b}$ (in eV) are the activation energies for the forward and backward reactions, respectively. $k_{et,f}$ and $k_{et,b}$ (in s^-1^) are the forward and backward Marcus rates.

| PDB ID | D-A ID | Geom. | H_da_ | $\Delta G^{\circ}$ | $\lambda$ | $E_{a,f}$ | $E_{a,b}$ | $k_{et,f}$ | $k_{et,b}$ |
| --- | --- | --- | --- | --- | --- | --- | --- | --- | --- |
| 7TFS | 651-654 | T | 2.000 | -0.132 | 0.733 | 0.123 | 0.255 | 6.7E+08 | 4.0E+06 |
| 7TFS | 654-657 | S | 8.000 | 0.066 | 0.603 | 0.186 | 0.120 | 1.1E+09 | 1.4E+10 |
| 7TFS | 657-660 | T | 2.000 | -0.009 | 0.663 | 0.161 | 0.170 | 1.6E+08 | 1.1E+08 |
| 7TFS | 660-639 | S | 8.000 | -0.048 | 0.640 | 0.137 | 0.185 | 6.7E+09 | 1.1E+09 |
| 8E5F | 933-936 | T | 2.000 | -0.132 | 0.734 | 0.123 | 0.255 | 6.6E+08 | 4.0E+06 |
| 8E5F | 936-939 | S | 8.000 | 0.028 | 0.605 | 0.166 | 0.138 | 2.3E+09 | 6.7E+09 |
| 8E5F | 939-942 | T | 2.000 | -0.134 | 0.683 | 0.110 | 0.244 | 1.1E+09 | 6.4E+06 |
| 8E5F | 942-921 | S | 8.000 | 0.061 | 0.618 | 0.187 | 0.126 | 1.0E+09 | 1.1E+10 |
| 8E5G | 1125-1128 | T | 2.000 | -0.105 | 0.682 | 0.122 | 0.227 | 7.2E+08 | 1.2E+07 |
| 8E5G | 1128-1131 | S | 8.000 | 0.016 | 0.579 | 0.153 | 0.137 | 3.8E+09 | 7.1E+09 |
| 8E5G | 1131-1134 | T | 2.000 | -0.050 | 0.691 | 0.149 | 0.199 | 2.6E+08 | 3.7E+07 |
| 8E5G | 1134-1137 | S | 8.000 | -0.009 | 0.573 | 0.139 | 0.148 | 6.6E+09 | 4.7E+09 |
| 6EF8 | 1272-1266 | T | 2.000 | -0.119 | 0.755 | 0.134 | 0.253 | 4.3E+08 | 4.3E+06 |
| 6EF8 | 1266-1269 | S | 8.000 | -0.057 | 0.567 | 0.115 | 0.172 | 1.7E+10 | 1.9E+09 |
| 6EF8 | 1269-1263 | T | 2.000 | 0.223 | 0.657 | 0.295 | 0.072 | 9.3E+05 | 5.2E+09 |
| 6EF8 | 1263-1260 | S | 8.000 | -0.112 | 0.543 | 0.086 | 0.198 | 5.3E+10 | 7.0E+08 |
| 6EF8 | 1260-1275 | T | 2.000 | -0.112 | 0.663 | 0.114 | 0.226 | 9.8E+08 | 1.3E+07 |
| 6EF8 | 1275-1254 | S | 8.000 | -0.048 | 0.554 | 0.116 | 0.164 | 1.7E+10 | 2.6E+09 |
| 6NEF | 1260-1263 | T | 2.000 | -0.111 | 0.770 | 0.141 | 0.252 | 3.3E+08 | 4.5E+06 |
| 6NEF | 1263-1266 | S | 8.000 | -0.014 | 0.601 | 0.143 | 0.157 | 5.4E+09 | 3.2E+09 |
| 6NEF | 1266-1269 | T | 2.000 | 0.232 | 0.654 | 0.300 | 0.068 | 7.5E+05 | 6.0E+09 |
| 6NEF | 1269-1272 | S | 8.000 | -0.067 | 0.553 | 0.107 | 0.174 | 2.3E+10 | 1.7E+09 |
| 6NEF | 1272-1275 | T | 2.000 | -0.043 | 0.652 | 0.142 | 0.185 | 3.4E+08 | 6.4E+07 |
| 6NEF | 1275-1242 | S | 8.000 | -0.154 | 0.565 | 0.075 | 0.229 | 7.9E+10 | 2.1E+08 |
| 7LQ5 | 562-553 | T | 2.000 | 0.150 | 0.658 | 0.248 | 0.098 | 5.6E+06 | 1.9E+09 |
| 7LQ5 | 553-559 | S | 8.000 | -0.079 | 0.547 | 0.100 | 0.179 | 3.0E+10 | 1.4E+09 |
| 7LQ5 | 559-550 | S | 8.000 | -0.098 | 0.594 | 0.104 | 0.202 | 2.5E+10 | 5.7E+08 |
| 7LQ5 | 550-541 | T | 2.000 | -0.136 | 0.646 | 0.101 | 0.237 | 1.7E+09 | 8.8E+06 |
| 7LQ5 | 541-547 | S | 8.000 | -0.028 | 0.601 | 0.137 | 0.165 | 7.0E+09 | 2.4E+09 |
| 7LQ5 | 547-544 | T | 2.000 | -0.059 | 0.609 | 0.124 | 0.183 | 7.0E+08 | 7.2E+07 |
| 7LQ5 | 544-844 | S | 8.000 | 0.090 | 0.558 | 0.188 | 0.098 | 9.9E+08 | 3.2E+10 |
| 8D9M | 562-559 | T | 2.000 | 0.091 | 0.683 | 0.219 | 0.128 | 1.7E+07 | 5.7E+08 |
| 8D9M | 559-553 | S | 8.000 | -0.115 | 0.563 | 0.089 | 0.204 | 4.6E+10 | 5.3E+08 |
| 8D9M | 553-550 | S | 8.000 | -0.200 | 0.640 | 0.076 | 0.276 | 7.2E+10 | 3.1E+07 |
| 8D9M | 550-547 | T | 2.000 | 0.012 | 0.681 | 0.176 | 0.164 | 8.9E+07 | 1.4E+08 |
| 8D9M | 547-544 | S | 8.000 | -0.077 | 0.622 | 0.119 | 0.196 | 1.3E+10 | 6.8E+08 |
| 8D9M | 544-541 | T | 2.000 | 0.045 | 0.630 | 0.181 | 0.136 | 7.7E+07 | 4.4E+08 |
| 8D9M | 541-844 | S | 8.000 | 0.090 | 0.564 | 0.190 | 0.100 | 9.3E+08 | 3.0E+10 |

**Supplementary Table 9.** Energetic parameters for Omc- E S and Z ‘nanowires’ obtained with the BioDC program. H_da_ (in meV) is the electronic coupling and is assigned a value of 8 or 2 meV for S and T geometries, respectively. $\Delta G^{\circ}$ (in eV) is the reaction free energy obtained by oxidizing hemes in the linear sequence prescribed by the filament topology while including heme-heme interactions. $\lambda$ (in eV) is the outer-sphere reorganization energy estimated by the MC, VEG, SC80, SC56, and AP methods, as defined in the main text. $E_{a,f}$, and $E_{a,b}$ (in eV) are the activation energies for the forward and backward reactions, respectively

| PDB ID | Method | Step | D-A ID | H_da_ | $\Delta G^{\circ}$ | $\lambda$ | $E_{a,f}$ | $E_{a,b}$ |
| --- | --- | --- | --- | --- | --- | --- | --- | --- |
| 7TFS | MC | 1 | 651-654 | 2.000 | -0.132 | 0.685 | 0.112 | 0.244 |
| 7TFS | MC | 2 | 654-657 | 8.000 | 0.066 | 0.566 | 0.176 | 0.110 |
| 7TFS | MC | 3 | 657-660 | 2.000 | -0.009 | 0.736 | 0.180 | 0.189 |
| 7TFS | MC | 4 | 660-639 | 8.000 | -0.048 | 0.654 | 0.140 | 0.188 |
| 7TFS | VEG | 1 | 651-654 | 2.000 | -0.132 | 0.749 | 0.127 | 0.259 |
| 7TFS | VEG | 2 | 654-657 | 8.000 | 0.066 | 0.544 | 0.171 | 0.105 |
| 7TFS | VEG | 3 | 657-660 | 2.000 | -0.009 | 0.925 | 0.227 | 0.236 |
| 7TFS | VEG | 4 | 660-639 | 8.000 | -0.048 | 0.713 | 0.155 | 0.203 |
| 7TFS | SC80 | 1 | 651-654 | 2.000 | -0.132 | 0.599 | 0.091 | 0.223 |
| 7TFS | SC80 | 2 | 654-657 | 8.000 | 0.066 | 0.435 | 0.144 | 0.078 |
| 7TFS | SC80 | 3 | 657-660 | 2.000 | -0.009 | 0.740 | 0.181 | 0.190 |
| 7TFS | SC80 | 4 | 660-639 | 8.000 | -0.048 | 0.570 | 0.120 | 0.168 |
| 7TFS | SC56 | 1 | 651-654 | 2.000 | -0.132 | 0.419 | 0.049 | 0.181 |
| 7TFS | SC56 | 2 | 654-657 | 8.000 | 0.066 | 0.305 | 0.113 | 0.047 |
| 7TFS | SC56 | 3 | 657-660 | 2.000 | -0.009 | 0.518 | 0.125 | 0.134 |
| 7TFS | SC56 | 4 | 660-639 | 8.000 | -0.048 | 0.399 | 0.077 | 0.125 |
| 7TFS | AP | 1 | 651-654 | 2.000 | -0.132 | 0.566 | 0.083 | 0.215 |
| 7TFS | AP | 2 | 654-657 | 8.000 | 0.066 | 0.438 | 0.145 | 0.079 |
| 7TFS | AP | 3 | 657-660 | 2.000 | -0.009 | 0.823 | 0.201 | 0.210 |
| 7TFS | AP | 4 | 660-639 | 8.000 | -0.048 | 0.488 | 0.099 | 0.147 |
| 6EF8 | MC | 1 | 1272-1266 | 2.000 | -0.119 | 0.768 | 0.137 | 0.256 |
| 6EF8 | MC | 2 | 1266-1269 | 8.000 | -0.057 | 0.618 | 0.127 | 0.184 |
| 6EF8 | MC | 3 | 1269-1263 | 2.000 | 0.223 | 0.651 | 0.293 | 0.070 |
| 6EF8 | MC | 4 | 1263-1260 | 8.000 | -0.112 | 0.546 | 0.086 | 0.198 |
| 6EF8 | MC | 5 | 1260-1275 | 2.000 | -0.112 | 0.665 | 0.115 | 0.227 |
| 6EF8 | MC | 6 | 1275-1254 | 8.000 | -0.048 | 0.584 | 0.123 | 0.171 |
| 6EF8 | VEG | 1 | 1272-1266 | 2.000 | -0.119 | 0.702 | 0.121 | 0.240 |
| 6EF8 | VEG | 2 | 1266-1269 | 8.000 | -0.057 | 0.429 | 0.081 | 0.138 |
| 6EF8 | VEG | 3 | 1269-1263 | 2.000 | 0.223 | 0.589 | 0.280 | 0.057 |
| 6EF8 | VEG | 4 | 1263-1260 | 8.000 | -0.112 | 0.696 | 0.123 | 0.235 |
| 6EF8 | VEG | 5 | 1260-1275 | 2.000 | -0.112 | 0.551 | 0.087 | 0.199 |
| 6EF8 | VEG | 6 | 1275-1254 | 8.000 | -0.048 | 0.487 | 0.099 | 0.147 |
| 6EF8 | SC80 | 1 | 1272-1266 | 2.000 | -0.119 | 0.561 | 0.087 | 0.206 |
| 6EF8 | SC80 | 2 | 1266-1269 | 8.000 | -0.057 | 0.343 | 0.060 | 0.117 |
| 6EF8 | SC80 | 3 | 1269-1263 | 2.000 | 0.223 | 0.471 | 0.256 | 0.033 |
| 6EF8 | SC80 | 4 | 1263-1260 | 8.000 | -0.112 | 0.557 | 0.089 | 0.201 |
| 6EF8 | SC80 | 5 | 1260-1275 | 2.000 | -0.112 | 0.441 | 0.061 | 0.173 |
| 6EF8 | SC80 | 6 | 1275-1254 | 8.000 | -0.048 | 0.390 | 0.075 | 0.123 |
| 6EF8 | SC56 | 1 | 1272-1266 | 2.000 | -0.119 | 0.393 | 0.048 | 0.167 |
| 6EF8 | SC56 | 2 | 1266-1269 | 8.000 | -0.057 | 0.240 | 0.035 | 0.092 |
| 6EF8 | SC56 | 3 | 1269-1263 | 2.000 | 0.223 | 0.330 | 0.232 | 0.009 |
| 6EF8 | SC56 | 4 | 1263-1260 | 8.000 | -0.112 | 0.390 | 0.050 | 0.162 |
| 6EF8 | SC56 | 5 | 1260-1275 | 2.000 | -0.112 | 0.309 | 0.031 | 0.143 |
| 6EF8 | SC56 | 6 | 1275-1254 | 8.000 | -0.048 | 0.273 | 0.046 | 0.094 |
| 6EF8 | AP | 1 | 1272-1266 | 2.000 | -0.119 | 0.236 | 0.015 | 0.134 |
| 6EF8 | AP | 2 | 1266-1269 | 8.000 | -0.057 | 0.554 | 0.111 | 0.168 |
| 6EF8 | AP | 3 | 1269-1263 | 2.000 | 0.223 | 0.436 | 0.249 | 0.026 |
| 6EF8 | AP | 4 | 1263-1260 | 8.000 | -0.112 | 0.649 | 0.111 | 0.223 |
| 6EF8 | AP | 5 | 1260-1275 | 2.000 | -0.112 | 0.257 | 0.020 | 0.132 |
| 6EF8 | AP | 6 | 1275-1254 | 8.000 | -0.048 | 0.676 | 0.146 | 0.194 |
| 7LQ5 | MC | 1 | 562-553 | 2.000 | 0.150 | 0.676 | 0.252 | 0.102 |
| 7LQ5 | MC | 2 | 553-559 | 8.000 | -0.079 | 0.641 | 0.123 | 0.202 |
| 7LQ5 | MC | 3 | 559-550 | 8.000 | -0.098 | 0.635 | 0.114 | 0.212 |
| 7LQ5 | MC | 4 | 550-541 | 2.000 | -0.136 | 0.624 | 0.095 | 0.231 |
| 7LQ5 | MC | 5 | 541-547 | 8.000 | -0.028 | 0.621 | 0.142 | 0.170 |
| 7LQ5 | MC | 6 | 547-544 | 2.000 | -0.059 | 0.617 | 0.126 | 0.185 |
| 7LQ5 | MC | 7 | 544-844 | 8.000 | 0.090 | 0.538 | 0.183 | 0.093 |
| 7LQ5 | VEG | 1 | 562-553 | 2.000 | 0.150 | 0.613 | 0.237 | 0.087 |
| 7LQ5 | VEG | 2 | 553-559 | 8.000 | -0.079 | 0.809 | 0.165 | 0.244 |
| 7LQ5 | VEG | 3 | 559-550 | 8.000 | -0.098 | 1.059 | 0.218 | 0.316 |
| 7LQ5 | VEG | 4 | 550-541 | 2.000 | -0.136 | 0.842 | 0.148 | 0.284 |
| 7LQ5 | VEG | 5 | 541-547 | 8.000 | -0.028 | 0.741 | 0.172 | 0.200 |
| 7LQ5 | VEG | 6 | 547-544 | 2.000 | -0.059 | 0.744 | 0.158 | 0.217 |
| 7LQ5 | VEG | 7 | 544-844 | 8.000 | 0.090 | 0.689 | 0.220 | 0.130 |
| 7LQ5 | SC80 | 1 | 562-553 | 2.000 | 0.150 | 0.490 | 0.209 | 0.059 |
| 7LQ5 | SC80 | 2 | 553-559 | 8.000 | -0.079 | 0.647 | 0.125 | 0.204 |
| 7LQ5 | SC80 | 3 | 559-550 | 8.000 | -0.098 | 0.847 | 0.166 | 0.264 |
| 7LQ5 | SC80 | 4 | 550-541 | 2.000 | -0.136 | 0.673 | 0.107 | 0.243 |
| 7LQ5 | SC80 | 5 | 541-547 | 8.000 | -0.028 | 0.593 | 0.135 | 0.163 |
| 7LQ5 | SC80 | 6 | 547-544 | 2.000 | -0.059 | 0.595 | 0.121 | 0.180 |
| 7LQ5 | SC80 | 7 | 544-844 | 8.000 | 0.090 | 0.551 | 0.186 | 0.096 |
| 7LQ5 | SC56 | 1 | 562-553 | 2.000 | 0.150 | 0.343 | 0.177 | 0.027 |
| 7LQ5 | SC56 | 2 | 553-559 | 8.000 | -0.079 | 0.453 | 0.077 | 0.156 |
| 7LQ5 | SC56 | 3 | 559-550 | 8.000 | -0.098 | 0.593 | 0.103 | 0.201 |
| 7LQ5 | SC56 | 4 | 550-541 | 2.000 | -0.136 | 0.471 | 0.060 | 0.196 |
| 7LQ5 | SC56 | 5 | 541-547 | 8.000 | -0.028 | 0.415 | 0.090 | 0.118 |
| 7LQ5 | SC56 | 6 | 547-544 | 2.000 | -0.059 | 0.417 | 0.077 | 0.136 |
| 7LQ5 | SC56 | 7 | 544-844 | 8.000 | 0.090 | 0.386 | 0.147 | 0.057 |
| 7LQ5 | AP | 1 | 562-553 | 2.000 | 0.150 | 0.461 | 0.202 | 0.052 |
| 7LQ5 | AP | 2 | 553-559 | 8.000 | -0.079 | 0.572 | 0.106 | 0.185 |
| 7LQ5 | AP | 3 | 559-550 | 8.000 | -0.098 | 0.565 | 0.096 | 0.194 |
| 7LQ5 | AP | 4 | 550-541 | 2.000 | -0.136 | 0.634 | 0.098 | 0.234 |
| 7LQ5 | AP | 5 | 541-547 | 8.000 | -0.028 | 0.723 | 0.167 | 0.195 |
| 7LQ5 | AP | 6 | 547-544 | 2.000 | -0.059 | 0.441 | 0.083 | 0.142 |
| 7LQ5 | AP | 7 | 544-844 | 8.000 | 0.090 | 0.431 | 0.157 | 0.067 |
